# Astrocyte modulation of neural activity during intracortical microstimulation

**DOI:** 10.64898/2026.10.25.759836

**Authors:** Kevin C. Stieger, Alberto Vazquez, Takashi D.Y. Kozai

## Abstract

Electrical manipulation of neural circuits has advanced our understanding of brain function, but the mechanisms that shape neural activity during stimulation remain incompletely understood. This gap reflects a persistent neuron-centric view of stimulation that overlooks the regulatory influence of glial cells, especially for long-duration stimuli. Here we investigated whether astrocytes actively shape the brain’s response to different patterns of stimulation. Using two-photon and mesoscale calcium imaging of the visual cortex in awake mice, we bidirectionally manipulated astrocyte calcium signaling during intracortical microstimulation (ICMS) via chemogenetic (hM3Dq) and optogenetic (melanopsin) tools. We show that astrocytes respond to stimulation in a pattern-dependent manner and shape ongoing neural activity rather than simply following it. Near the electrode, neurons responded to astrocyte modulation in mixed directions, some increasing and some decreasing. Instead of uniformly raising or lowering activity, modulating astrocyte calcium pulled neuronal responses toward an intermediate state, with the direction determined by each neuron’s starting response and its distance from the electrode. Our results identify astrocytes as a modulator of sustained neuronal activity during electrical stimulation and highlight the need to consider neuron-glia dynamics in the design of therapeutic neurostimulation.

## 1. Introduction

Electrical stimulation of the nervous system has played a central role in revealing fundamental aspects of brain activity and has led to important therapies for neurological disorders and sensory neuroprostheses^1–5^. Following the seminal work of Fritsch and Hitzig^6^ and later Penfield and colleagues^5^, researchers used electrical stimulation to map brain function and identify how specific brain regions and cell populations contribute to behavior and perception. For example, stimulating the middle temporal (MT) area in non-human primates (NHPs) biased how animals perceived the direction of moving dots, depending on the directional tuning properties of specific neurons in that region, linking it to motion perception^1^. Additionally, NHPs trained to discriminate tactile stimuli could also distinguish different frequencies of intracortical microstimulation (ICMS) with similar accuracy^7^. This finding highlighted that patterned electrical activity in somatosensory cortex is sufficient to encode tactile sensations, which is a foundational principle underlying sensory neuroprostheses^7^. Finally, the discovery that stimulation of the thalamus at specific frequencies reduces tremors in Parkinson’s Disease (PD) led to the development of deep brain stimulation (DBS) therapies, which are now widely used to treat Parkinson’s Disease, epilepsy, depression, and other neurological disorders by disrupting abnormal neuronal activity patterns^2,8^. More recently, bursting stimulation targeted to basal ganglia circuits was shown to reduce ataxia in a mouse model of PD by selectively modulating different cell types^9^. Unlike conventional high frequency stimulation, which only works during stimulation, this bursting pattern produced lasting improvements after stimulation stopped. Together, these studies highlight that effective brain stimulation depends on where it is delivered, its temporal pattern, and which circuits and cell types it engages.

Despite these advances, the full therapeutic potential of brain stimulation remains limited because it is often unclear which cell types are activated by the stimulus and how they coordinate to shape the overall network response^10^. Identifying how specific neuronal and glial populations respond to stimulation, either through direct activation or via synaptic recruitment, is essential to improve stimulation-based therapies. Sensory processing itself depends on coordinated interactions between neurons and glial cells, which together shape perception^11–13^. For sensory neuroprostheses to be effective, artificial percepts must be stable and not rapidly fade over time^14,15^. Therefore, understanding how ICMS recruits different cell types and how their interactions influence network dynamics is critical^10^. One major challenge is perceptual fading, where artificial sensations weaken quickly during continuous stimulation^14,15^. Notably, calcium imaging studies in mice showed that neurons farther from the electrode exhibit brief activation followed by rapid adaptation, occurring on a similar time scale as the perceptual fading observed in humans^15,16^. Using short bursts of stimulation instead of continuous high frequency trains can repeatedly activate neurons in mice, reducing neuronal adaptation, and can reduce perceptual fading in humans^17,18^.

This adaptation likely arises from multiple interacting mechanisms^19^, including inhibitory network stabilization^20^, neurotransmitter or metabolic depletion^21,22^, glial signaling^13,23–25^, mural cell vascular coordination^26–28^, and intrinsic neuronal properties^21^. In particular, astrocytes have long been proposed as key contributors to the therapeutic effects of deep brain stimulation^29–31^. For example, astrocytes show increases in calcium activity during DBS^23^, which correlates with elevated extracellular adenosine levels that are necessary for tremor reduction in rodent models of PD. Within the hippocampus, astrocytes release ATP following prolonged or high frequency stimulation of interneurons, leading to reduced synaptic transmission^32^. Furthermore, disrupting astrocyte calcium signaling reduces neuronal adaptation during sensory stimulation, and in a separate model, impairs fine but not coarse texture discrimination, supporting roles for astrocytes in both sensory response dynamics and perceptual precision^13,33^. Astrocytes in spinal sensory circuits can exert opposing effects on nociceptive processing depending on the way in which they are activated. Prolonged depolarizing optogenetic activation of spinal astrocytes induced pain hypersensitivity in rats^34^, whereas Aβ-fiber stimulation recruits astrocytes that contribute to adenosine-dependent suppression of nociceptive transmission and antinociception^35^. Because epidural spinal cord stimulation recruits large-diameter Aβ sensory fibers^36,37^, it may engage similar astrocyte-mediated inhibitory mechanisms. These findings highlight the importance of determining how astrocytes are recruited by electrical stimulation and how they shape the resulting neuronal responses, yet these interactions remain understudied in the cortex^38,39^.

This study asks whether cortical astrocytes respond to intracortical microstimulation and whether their calcium signaling shapes stimulation-evoked neural activity. We combined genetically encoded calcium indicators in astrocytes and neurons with chemogenetic and optogenetic control of astrocyte calcium during ICMS, and imaged at two spatial scales to separate directly recruited neurons near the electrode from synaptically recruited neurons across the wider network. By raising and lowering astrocyte calcium during stimulation while tracking neuronal responses, we tested whether astrocytes actively shape ongoing activity and, if so, where and under which stimulation patterns.

## 2. Results

To investigate whether astrocytes actively shape the temporal progression of stimulation-induced neural activity we performed in vivo two-photon and mesoscale calcium imaging in the visual cortex of awake mice. We monitored astrocyte calcium transients because calcium-dependent signaling can trigger gliotransmitter release and thereby influence neural activity^40,41^. Neuronal calcium responses to stimulation were monitored, which our previous work showed evokes pattern-dependent activity that parallel changes in perceptual magnitude reported in humans during clinical ICMS^15,42,43^. Finally, to evaluate how astrocytic influence scales with neuronal adaptation versus facilitation, we recorded these dynamics across a 30-s train of intracortical microstimulation (ICMS) using four distinct temporal paradigms: (1) continuous 100 Hz, which drives rapid neural adaptation; (2) continuous 10 Hz, which promotes sustained neural activity; (3) a bursting paradigm consisting of 10 pulses delivered at 100 Hz every second (10Hz-burst), which is associated with sustained sensory perception; and (4) theta-burst stimulation (TBS) consisting of 2 pulses delivered at 100 Hz repeated at 5 Hz every second, a paradigm commonly used to drive circuit-level plasticity.

### 2.1 Astrocytes Display Calcium Activity During ICMS On Relevant Timescales to Modulate Neural Activity

We first asked whether astrocytes become active during ICMS on spatial and temporal scales that would allow them to affect neurons. AAVs were used to label astrocytes and neurons with different calcium sensors (Aldh1l1-targeted astrocytes expressed GCaMP8m, and syn-targeted neurons expressed jRGECO1a) to measure their stimulation induced activity simultaneously (Fig. 1a). On the day of experimentation, a single shank Michigan-style electrode was implanted into layer II/III of the visual cortex, under isoflurane anesthesia. The window was sealed and the mouse was allowed to wake up and recover for at least one hour in its home cage before stimulation experiments began.

**Figure 1.**
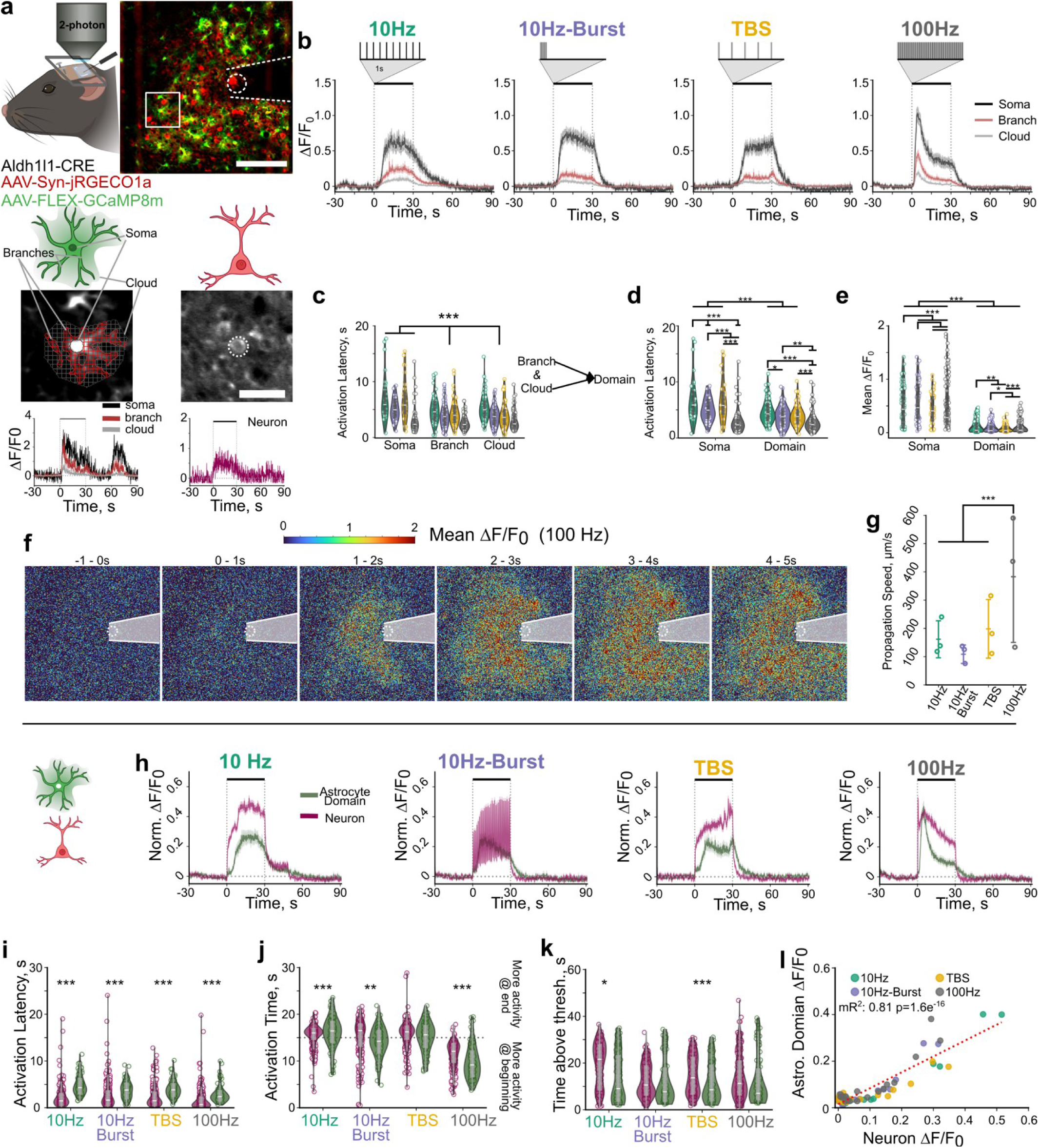
ICMS elicits calcium activity within astrocytes on comparable time and spatial scale as neurons under two-photon microscopy. **A.** (top) Aldh1l1-CRE mice expressing GCaMP8m in astrocytes and jRGeCO1a in neurons were implanted with an electrode in the visual cortex and head fixed under a two-photon microscope. Representative image highlighting astrocyte morphology (green) (See Fig. S1) and neurons (red). Scale bar = 100 µm. (Bottom) Astrocytes were segmented into the soma, main branches and cloudy region around the astrocyte. Representative traces of astrocyte regions (left) and a neuron (right) during 100 Hz ICMS **B.** Astrocytes display diverse calcium responses between soma, branch, and cloud regions to different stimulation paradigms. **C.** Astrocyte branches and cloud domains activate at a similar latency to stimulation, and earlier than the soma (LMM; Stim: 1.7e-82, ROI type: p=5.7e-10, Stim x ROI type: p=4.1e-12). Branches and cloud region were averaged together for further analysis. **D.** Stimulation parameters influence the activation latency of astrocyte somata and domains (LMM; Stim: p=1.1e-58, ROI type: p=3.2e-8, Stim x ROI type : p=1.1e-11, trial: p = 0.82). **E**. Astrocyte soma displays larger mean calcium activity during 30s ICMS, and evoked activity depends on stimulation pattern (LMM; Stim: 9.8e-87, ROI type: p=1.4e-9, Stim x ROI type: p=1.9e-50, trial: p = 5.6e-4). **F.** During ICMS calcium activity tends to propagate away from the stimulation site. Images represent mean ΔF/F0 across corresponding time ranges. **G**. Calcium activity spreads fastest during 100 Hz stimulation Hz compared to other patterns (LMM: stim: 1.3e-6). **H**. Max normalized calcium traces ± standard error highlight diverse temporal distributions of astrocyte domains (green) and neuron (red) calcium activity depending on stimulation pattern. **I**. Astrocyte domains activate later than neurons (LMM; Stim: p=1.2e-48, cell type: p=3.7e-14, interaction: p=4.3e-3, trial: p=0.49). **J**. Activation Time of neurons and astrocytes shows temporal distribution of calcium activity depends on stimulation and differs between cell types (LMM; Stim: p=2.6e-251, cell type: p=0.57, interaction: p=9.5e-21, trial p=0.05). **K**. Time spent above threshold depends on stimulation pattern and cell type (LMM; Stim: p=2.1e-57, cell type: p=0.15, interaction: p=4.3e-7, trial: p=0.01). **L**. The magnitude of population-level astrocyte activity correlates with neuronal activity (LMM; p = 1.6e-16) For c-e, N=3, n=31-82, 6 trials. I-L: N=3 animals, 6 trials. neurons: n=215-280, astrocytes 72-82 *: P<0.05, **P<0.01, *** p<0.001.

#### 2.1.1 Astrocyte Compartment-Specific Responses

We wanted to determine how astrocytes respond to different stimulation parameters before comparing their responses to neurons. The strong baseline fluorescence of astrocytic GCaMP8m enabled segmentation of individual astrocytes into somata, main branches and surrounding fine-process-enriched “cloudy” regions with clear separation from the neurons (Fig. 1a, Fig. S1). Using a semi-automated approach similar to CalSee^33^, branches and cloudy regions were further divided into 3 x 3 µm spatial domains for analysis. This compartmentalization was important because astrocytic calcium signaling is spatially heterogeneous, with branches and fine processes exhibiting distinct calcium dynamics and close functional relationships with synaptic activity^44,45^. For each astrocyte, calcium signals from all active domains within a given region were averaged to represent that region in subsequent analyses (Fig. S2, Fig. 1a).

Astrocyte somata consistently showed the greatest increases in calcium during stimulation (Fig.1B, Fig. S3) and activated later than branches and cloudy regions (Fig.S3). In contrast, the cloudy region exhibited the shortest activation latency, consistent with the idea that perisynaptic astrocyte processes detect neural activity first^45–47^. This pattern is consistent with models in which astrocytes integrate calcium signals originating in fine processes before generating a larger somatic calcium event, with the perisynaptic processes responding before the soma and tracking the timing of neuronal activity more closely^46^. For the rest of our analyses, calcium activity from the branch and cloudy domains was averaged together and referred to as the astrocyte ‘domain’ signal, providing a concise representation of perisynaptic and branch activity.

We then asked whether stimulation pattern influences the timing and strength of astrocyte calcium activity. Consistent with the above results, the soma activated later than distal domains, and depended on stimulation pattern (Fig. 1d). 100Hz elicited the largest calcium transients (Fig. 1e) and its rise was faster compared to other stimulation patterns (Fig. S3). However, 10Hz stimulation elicited above-threshold increases for a comparable duration as 100 Hz (Fig. S3). Interestingly, astrocyte calcium activity often persisted after stimulation ended, and 10 Hz stimulation resulted in the longest time for domain activity to return to baseline (Fig.S3). These results indicate that both the speed and persistence of astrocyte calcium changes are strongly dependent on the temporal structure of stimulation, suggesting that different patterns may engage distinct modulatory or integrative functions in astrocytes.

#### 2.1.2 Spatial Propagation of Calcium Activity

Neurons within ∼100 µm of the electrode are likely directly activated by ICMS, while neurons beyond ∼250 µm are more likely to be recruited synaptically^42^, allowing us to probe astrocyte influence across direct and network-mediated pathways^42^. By examining the spatial patterns of astrocyte activity, we noticed that calcium signals often appeared first near the electrode and then spread outward, especially during 100 Hz stimulation (Supplemental Movie 1). We therefore asked whether the speed of calcium propagation depended on the stimulation pattern, which could reflect a coordinated astrocytic response across the network. To test this, we compared the timing of calcium activity within regions located <200 µm from the electrode to regions located >200 µm away. Our analysis showed that calcium signals appeared to propagate fastest during 100 Hz stimulation (Fig. 1f-g), suggesting that high-frequency stimulation can induce rapid, spatially organized waves of astrocytic calcium activity.

#### 2.1.3 Astrocyte-Neuron Timing Comparisons

Next, we compared stimulation-evoked calcium signals in astrocyte domains and neurons recorded simultaneously under two-photon (Fig. 1h-l). Because the domains displayed a shorter activation latency, we focused this comparison on domain activity rather than somatic signals (Fig. 1h). Across all stimulation patterns, astrocyte domain activity began later than neuronal activity (Fig. 2h-I, Supplemental Movie S2), despite the faster kinetics of the GcaMP8m sensor compared to jRGeco1a^48,49^. This temporal ordering supports the interpretation that astrocyte activity is driven by neuronal activity instead of being directly activated by the electrical stimulus^38^. The timing also provides a mechanistic framework for the distance-dependent effects observed with astrocyte suppression. Near the electrode, neurons are more strongly driven by the electrical stimulus itself, limiting the extent to which astrocyte modulation can alter their responses. Farther from the electrode, where neuronal recruitment is more dependent on synaptic propagation^42^, the delayed astrocyte response occurs within the timescale of ongoing network activity and may therefore modulate subsequent neuronal dynamics.

**Figure 2:**
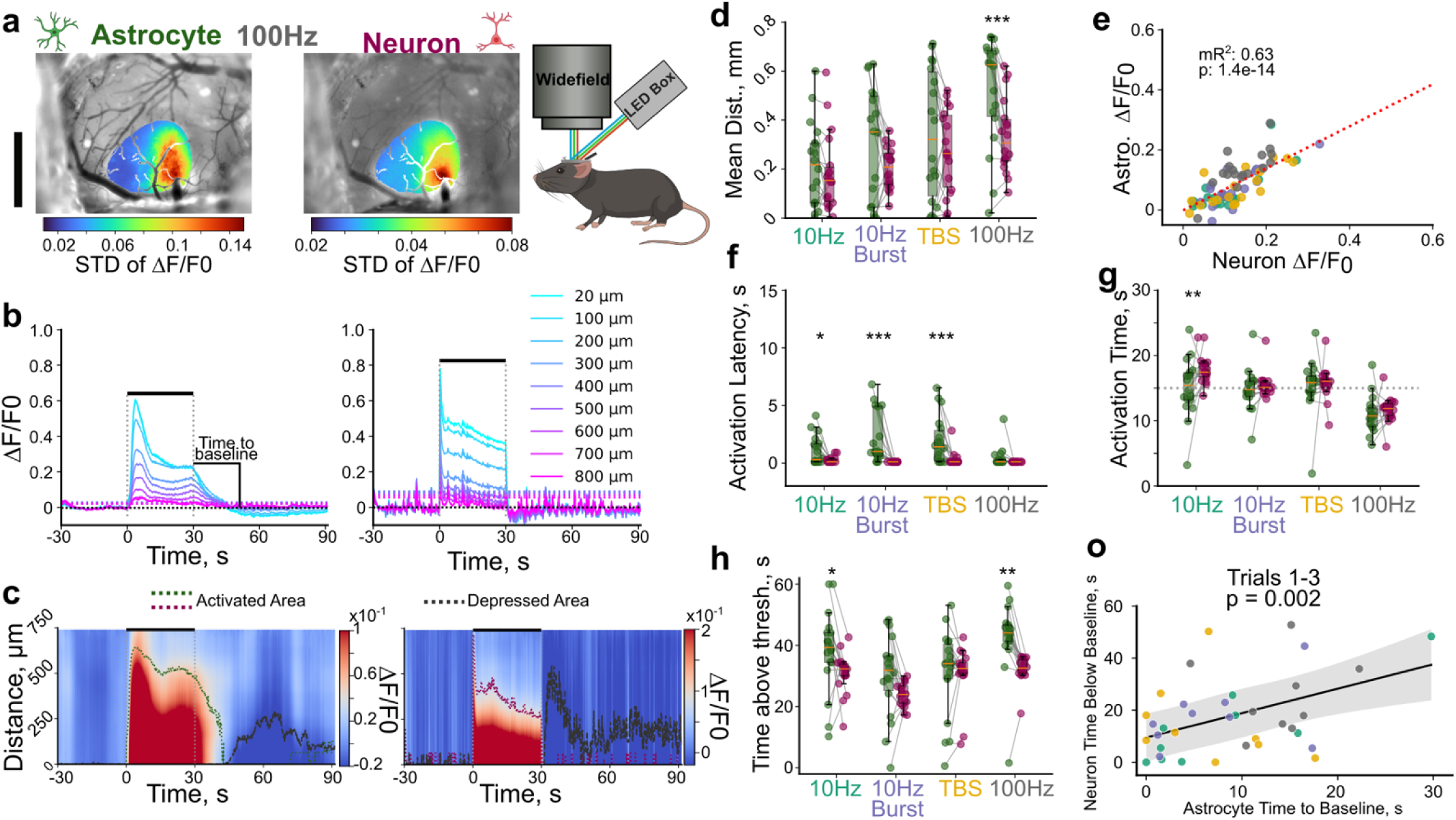
ICMS elicits calcium activity within astrocytes on comparable time and spatial scale as neurons under mesoscale imaging. **A**. Representative mesoscale image highlighting the standard deviation of astrocyte (left) or neuron (right) calcium activity during 100 Hz stimulation. **B**. Astrocyte and neuron activity decreases with distance from the center of activation. Dotted lines represent respective bin threshold levels for activation. **C.** Heatmaps of the average calcium activity and corresponding ‘activation distance’ for astrocytes (left, green dotted line) and neurons (right, red dotted line). Gray dotted line shows corresponding distance of ‘depressed’ (i.e. below baseline) activity **D.** Average distance of mesoscale astrocyte calcium activity greater than that for neurons specifically for 100 Hz (LMM; Stim: p=4.8e-7, cell type: p=1.1e-3, interaction: p=0.24, trial p=2.8e-3). **E**. Stimulation induced calcium activity correlates with the magnitude of neuronal calcium activity (lm: p = 1.4e-14). **F.** Mesoscale astrocyte activity elevates after neurons (LMM; Stim: p=5.5e-3, cell type: p=1.1e-8, interaction: p=0.01, trial p= 0.30). **G**. Activation time is significantly different between mesoscale astrocyte and neuron activity (LMM; Stim: p=1.3e-11, cell type: p=0.02, interaction: p=0.25, trial: p=0.08). **H**. Astrocyte calcium activity is above threshold significantly longer than neurons (LMM; Stim: p=1.7e-4, cell type: p=5.6e-4, interaction: p=0.18, trial: p=0.04). **I**. Depression of neuron activity after stimulation correlates with the time for astrocyte calcium activity to return to baseline for the first 3 trials (LMM astrocyte time to baseline p = 0.002). N=3 animals, 6 trials. *: p<0.05, **:p<0.01,***:p<0.001.

#### 2.1.4 Temporal Integration of Astrocyte Activity

We next asked whether astrocyte calcium activity is biased toward the beginning or the end of the stimulation period, which could provide insight into how astrocytes integrate activity over time. To determine this bias, we used a metric called ‘Activation time’, which represents the time required to reach half of the total calcium activity during the 30 s stimulation train^16^ (Fig. S2b). Calcium signals that build up over time yield activation times >15 s, while signals that adapt rapidly yield activation times <15 s (Fig. 1j). Astrocyte domain activation time depended strongly on stimulation pattern. During 10 Hz stimulation, astrocyte activity was weighted later in time than neuronal activity, while during 10 Hz burst and 100 Hz stimulation, astrocyte activity was weighted earlier than neuronal activity (Fig. 1j). The time spent above threshold also varied by stimulation pattern. These domains were above threshold for a shorter duration than neurons during 10 Hz and TBS, but were comparable during 10 Hz-burst and 100 Hz (Fig. 1k). This difference likely reflects that astrocyte domains may remain engaged to support processes such as gliotransmission that influence neuronal responses. Across every pattern tested, neurons activated before astrocytes. Even at 100 Hz, astrocytes followed several seconds later (Fig. 1i). Furthermore, this timing suggests that ICMS drives cortical neurons much as sensory input does, not through the electrical nature of the stimulus. It also places the astrocyte response within the window of ongoing network activity. Consistent with this interpretation, the magnitude of astrocyte domain calcium activity correlated with neuronal calcium activity across stimulation patterns (Fig. 1l), in agreement with previous reports^23,38^.

#### 2.1.5 Mesoscale Spread and Network-Level Correlation

While these analyses focused on astrocytes near the electrode^38^, ICMS recruits broader networks through synaptic connections, so we next asked how far astrocyte activation extends across the cortex and how it relates to network-level neuronal activity? Using mesoscale calcium imaging, we quantified the spatial spread and timing of astrocyte and neuronal activity across the cortex, specifically where the sensors are expressed (Fig. S4, Fig. 2a). Astrocytes displayed calcium activity over a larger radius than neurons, particularly during 100 Hz stimulation (Fig. 2c-d). The broader spatial spread of astrocyte activity extended beyond the primary neuronal activation zone, providing a spatial mechanism by which astrocytes could influence neurons recruited farther from the electrode through synaptic network propagation. To more effectively compare differences in astrocyte and neural activity within a stimulation condition, we computed the average calcium activity within a defined distance based on the mean and variance of the activation distance for neurons and astrocytes within each stimulation condition (Fig. S5). When comparing astrocyte and neuronal activity within equivalent spatial regions within a stimulation pattern, astrocyte calcium activity remained strongly correlated with neuronal activity (Fig. 2e). As expected from the two-photon data, astrocyte activation occurred later than neuronal activation (Fig. 2f). However, during stimulation, the overall temporal distribution of astrocyte and neuronal activity was similar across most patterns, except for 10 Hz stimulation as captured by the activation time metric (Fig. 2g).

A striking difference emerged after stimulation ended. Astrocyte calcium activity often remained elevated, while neuronal activity rapidly returned to baseline or even fell below baseline levels (Fig. 2c, h-i). As a result, astrocytes spent more time above threshold than neurons (Fig. 2h). Importantly, the duration of post-stimulation astrocyte activity associated with the duration of neuronal suppression (Fig. 2i, Fig. S5). Across all six trials this post-stimulation relationship changed significantly with trial number (Fig. S6; LMM astrocyte time to baseline p= 0.06, trial p=0.87, interaction p=0.024). In early trials, longer astrocyte activity predicted longer neuronal suppression (Fig. 2i; Trials 1-3, slope =0.94s/s, p=0.002), This indicated that the association depended on stimulation history. This relationship weakened with repeated stimulation and was absent by later trials (Trials 4-6, p=0.56 Fig. S6). While astrocytes also showed post-stimulation decreases in activity (Fig. 2i, Fig. S6), these correlations suggest that astrocyte signaling could contribute to the suppression of neuronal activity after stimulation.

#### 2.1.6 Dependence on trials

Clinical utility of intracortical microstimulation will ultimately require repeated stimulation for sensory replacement and feedback. It is therefore critical to consider whether responses attenuate over trials, and if this attenuation can be mitigated by certain stimulation parameters. Interestingly, the magnitude of astrocyte somata responses decayed more than that of domains, while the number of active domain ROIs decayed more than somata (Fig. S6). More importantly, this effect depended on stimulation pattern, where both bursting patterns resulted in the lowest attenuation in magnitude over trials in domain ROIs. The attenuation of neuronal activity over trials followed a similar pattern, consistent with the idea that astrocyte activity follows that of the neurons. Despite this decay in the magnitude of activity over trials, the temporal structure of the responses (i.e. activation time) remained largely stable over trials, with the exception of 10 Hz (Astrocytes and neurons) and TBS (neurons), indicating a larger attenuation during a stimulation train, by later trials. Moreover, these patterns were consistent in the mesoscale responses (Fig. S6).

#### 2.1.7 Summary

Taken together, these results indicate that astrocytes are not passive bystanders during ICMS. They respond to stimulation-evoked neural activity with calcium signals that are spatially extensive, temporally structured, and aligned with periods of neural adaptation or facilitation. These findings raise whether astrocytes simply reflect neural activity or actively shape the neural response to ICMS. Because spectral overlap between calcium indicators may confound correlation analyses, we next directly manipulated astrocyte calcium signaling to test its causal influence on neuronal activity.

### 2.2 Sustained GqDREADD Activation Suppresses Astrocyte Calcium and Reshapes Neuronal Responses to ICMS

Historically, reducing astrocyte calcium activity *in vivo* has been challenging^50–53^, where some models do not completely abolish calcium signaling in microdomains (IP3R -/-), and others can induce compensatory mechanisms (CalEx). Designer receptors exclusively activated by designer drugs (DREADDs) modulate astrocyte calcium signaling acutely^52^ permitting within-animal comparisons, but both Gq and Gi DREADDs have been shown to elicit increases in calcium activity shortly after agonist administration^52^. However, recent evidence suggests that sustained stimulation of the hM3Dq, could effectively exhaust intracellular calcium stores, providing a potential method to sufficiently elicit prolonged attenuation of astrocyte calcium signaling^54–56^.

Consistent with this idea, stimulation-induced astrocyte calcium activity (Flex-GCaMP8m, Aldh1l1-Cre) was virtually absent for most conditions 30 minutes after stimulating hM3Dq (GFAP-hM3Dq-mCherry) with an intraperitoneal injection of compound 21 (C21, 1 mg/kg; Fig. 3a-d, 4a-d, Fig. S7). In validation experiments, C21 alone did not broadly attenuate stimulation-induced astrocyte calcium responses across stimulation parameters in mice lacking the hM3Dq receptor (GFAP104-mCherry; Fig. S7). A reduction in the response to 100 Hz was observed, but activity clearly persisted during the other stimulation conditions (Fig. S7). These results indicate that sustained activation of the astrocyte-specific Gq DREADD with C21 effectively suppresses astrocyte calcium responses during ICMS, particularly at lower frequencies. At 100 Hz, evoked responses were significantly dampened (p = 4.2e-15) but not eliminated. This sets up the question of how such suppression of astrocyte calcium impacts neuronal activity, both locally and across synaptically recruited networks.

**Figure 3.**
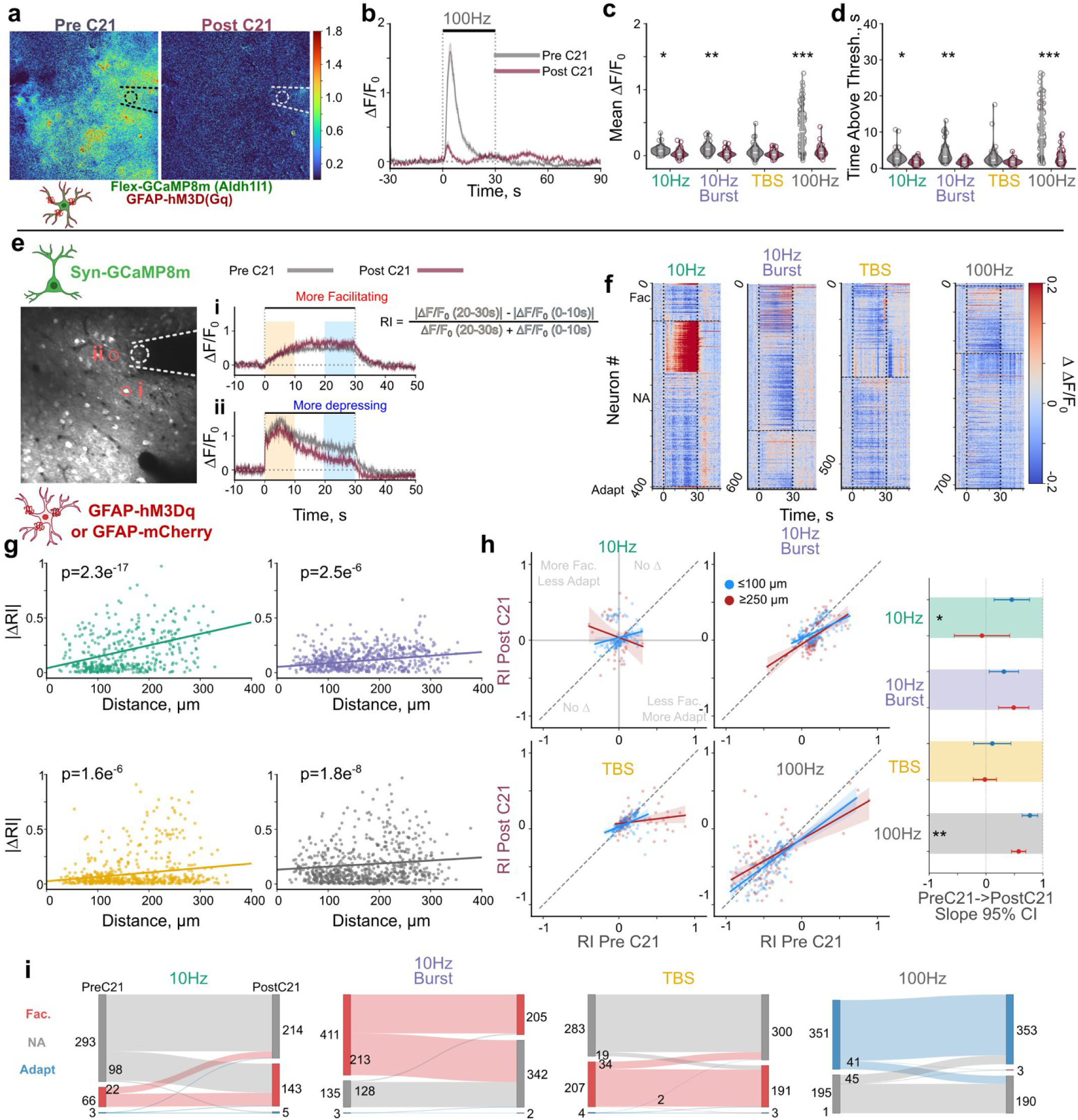
Attenuation of stimulation induced astrocyte calcium activity through DREADD activation may promote heterogeneous changes in neuronal calcium activity throughout the stimulation duration. **A.** Representative 100Hz stimulation averaged ΔF/F0 projection image of astrocytes before (left) and after (right) C21 administration in an animal model expressing GCaMP8m and hM3Dq in astrocytes. **B.** Stimulation induced calcium elevations during 100 Hz stimulation are substantially reduced after C21 administration. **C-D**. Activation of hM3Dq in astrocytes by C21 reduces stimulation induced mean calcium activity (C, LMM; Stim: p=2.8e-108, C21: p= 9.2e-6, interaction: p=1.1e-79) and time above threshold (D, LMM; Stim: p=1.4e-113, C21: p= 1.2e-5, interaction: p= 1.6e-75). N=3, n=80-86. **E**. Representative image of neurons expressing GCaMP8m (left). Following DREADD activation, some neurons displayed greater stimulation-induced calcium activity that tended to increase over time (i; TBS stimulation), while others displayed reduced calcium activity that tended to decrease more over time (ii; 100Hz stimulation). Formula represents the calculation of the response index (RI). **F.** Heatmaps showing difference in trial averaged calcium activity (Post-C21 – PreC21) highlights heterogeneous effect of attenuating astrocyte calcium responses on different neurons. Neurons grouped by response profile classification and sorted by difference in response index. **G**. The magnitude of the change in response index increases farther from the electrode (LMM: Distance p= 1.0e-32, Stim p = 0.08, interaction p= 8.3e-5; 10Hz: N=4, n=403; 10Hz-Burst: N=4, n=621; TBS: N=4, n=568; 100Hz: N=4, n = 713). **H.** left: Retention of response profile depends on stimulation condition and distance from the electrode (10Hz-prox/dist: N=4/3, n=107/39; 10Hz-Burst-prox/dist: N=4/4, n=139/105; TBS-prox/dist: N=4/4, n=125/93; 100Hz-prox/dist: N=4/4, n = 149/117). Right: 95% confidence intervals of slopes in left. **I.** Subsets of neurons display changes in response profile classification after attenuation of astrocyte calcium activity.

#### 2.2.1 Astrocyte Calcium Suppression Drives Bidirectional, Distance-Dependent Changes in Neuronal Responses

Having established that activation of astrocytic hM3Dq with C21 attenuates astrocyte calcium responses during ICMS, we asked how this astrocyte-specific suppression affects neuronal calcium dynamics. Astrocytes could provide a uniform gain on neuronal activity, such that reducing their calcium responses would lead to broad increases or decreases in neuronal activity. Alternatively, their influence may depend on how neurons are recruited within the local circuit. To address this, we expressed GCaMP8m in all neurons (syn-GCaMP8m) while maintaining hM3Dq expression in astrocytes (GFAP-hM3Dq-mCherry) and compared neuronal calcium signals before and after C21 administration within the same animals (Fig. 3e-i).

If astrocytes simply exert a stabilizing tone on neurons during ICMS, attenuating astrocyte responses would be expected to yield a global increase in the mean activity of neurons during ICMS potentially through more facilitating responses. However, this was not the case (Fig. S8) and visual inspection of the responses suggested that some neurons increase their activity while others decreased (Fig. 3e-f). To quantify this, we computed a response index (*RI*), comparing early versus late calcium activity within each 30-s ICMS train (Fig. 3e). In general, the *RI* values tend to distribute into 3 response categories that separates the neuronal dynamics into facilitating, where activity increases over time (Fac.: RI>0.10), non-adapting where activity tends to remain the same (NA: -0.32<RI<0.10), and adapting where activity decreases over time (Adapt.: RI<-0.32), consistent with previous work^42^ (Fig. S9). Grouping neurons by their initial response profile and viewing the difference in stimulation-evoked activity before and after C21 administration revealed heterogeneous effects across individual neurons (Fig. 3f). Importantly, these changes appeared to depend on each neuron’s initial response regime, where some neurons that were strongly facilitating before C21 shifted towards less facilitation, whereas strongly adapting neurons shifted toward less adaptation. Moreover, these effects depended on stimulation condition. Thus, suppression of astrocyte calcium responses to ICMS altered neuronal dynamics in a manner that appeared to depend on the manner in which a neuron was recruited within the circuit.

Electrical stimulation can recruit neurons through different mechanisms where neurons farther from the electrode are more likely to be recruited through synaptic pathways, while neurons near the electrode are more likely to be activated directly, we reasoned that astrocyte suppression would have a greater effect on distal neurons (≤ 100: ≥250 indirect)^42^. Therefore, we next asked *where* these changes were strongest by testing if the magnitude of the change in the response index depended on distance. Indeed, neurons farther from the electrode showed a greater change in the magnitude of trial-averaged response index across all stimulation conditions, with the strongest effect during 10 Hz stimulation (Fig. 3g). Thus, the increasing magnitude of modulation with distance is consistent with a greater astrocytic influence on neurons whose recruitment depends more heavily on local synaptic circuitry rather than direct recruitment. Neurons near the electrode, which are more likely directly activated by the electrical pulse, showed the smallest changes, as expected if their responses are dominated by the imposed drive rather than shaped by astrocyte signaling.

A neuron’s RI likely reflects the combined influence of direct recruitment, recurrent excitation, inhibition, intrinsic response properties and other circuit interactions that determine how its activity evolves during sustained stimulation. If astrocytes interact with these circuit processes, attenuation of astrocyte calcium should therefore alter the relationship between a neuron’s initial response regime and its subsequent response rather than producing a uniform displacement of all neurons. To resolve this, we regressed each neurons post-suppression RI against its pre-suppression RI within each stimulation condition and distance bin (Fig. 3h). Should astrocyte attenuation leave each neuron’s temporal response profile unchanged, post-C21 RI should remain close to pre-C21 RI, and the population should follow the unity line with a slope near 1. Deviation from this slope indicates that the effect of astrocyte attenuation depends on the initial response profile, and a difference in slope between local and distal populations indicates this effect is stronger on synaptically recruited neurons. Notably, these effects depended on both stimulation pattern and distance (Fig. S8, Fig. 3h). During 10 Hz and 100Hz, the relationship compressed toward zero with a significant shift in 10 Hz (LMM: PreC21: p=1.9e-16, Stim: p = 2.7e-16; Distance: p = 0.85, PreC21 x Stim: p=2.1e-14; PreC21 x Distance: p=0.02; Stim x Distance: p=0.04; PreC21 x Stim x Distance: p=0.02; 10 Hz proximal vs distal: p=0.01, 100 Hz proximal vs distal p = 0.003).

Although the slope of the distal 10 Hz neurons appears to be negative, which would indicate an inversion in the response of neurons, this was driven by only a few neurons and did not pass Cook’s test. Analysis across continuous distance further revealed a nonlinear spatial dependence, indicating that astrocyte modulation varies across the circuit rather than changing monotonically with distance. Finally, classification of neurons response profile before and after C21 administration showed that a subset underwent robust transitions between classes (Fig. 3i; see methods). Specifically, many neurons shift from non-adapting responses prior to astrocyte attenuation to facilitating during 10 Hz stimulation, while many neurons shift from facilitating to non-adapting during 10 Hz burst, and TBS stimulation.

Together, these data highlight that attenuating astrocyte calcium modulates individual neuronal responses to ICMS, particularly at greater distance from the electrode where neurons are more likely to be synaptically recruited, and during low frequency stimulation. Importantly, astrocyte calcium attenuation biased neuronal responses in opposite directions across the population, with some neurons becoming more facilitating and others more adapting, in a manner opposite to the adaptation produced by repeated stimulation. These bidirectional changes suggest that astrocytes may shape the temporal dynamics of population activity rather than simply increase or decrease neuronal responses.

#### 2.2.3 Mesoscale Imaging Reveals Network-Level Modulation

Because astrocyte-dependent effects were stronger at greater distances, we next asked how these heterogeneous cellular changes manifest when activity is integrated across a broader cortical network (Fig. 4). Mesoscale calcium imaging samples activity across a larger spatial field and multiple cortical layers^57^, providing a complementary measure of the network-level consequence of astrocyte attenuation. Consistent with the two-photon data, astrocyte activity was suppressed following activation of hM3D and not due to C21 administration in the absence of hM3D (Fig. 4a-d, Fig. S10). Correspondingly, broad neural activity increased during 10 Hz and TBS stimulation patterns (Fig. 4g-h) following astrocyte suppression, while mesoscale RI remained unchanged (Fig. 4i). Thus, although individual neurons local to the electrode displayed bidirectional changes, integration across the broader network revealed a net increase in neuronal activity.

**Figure 4.**
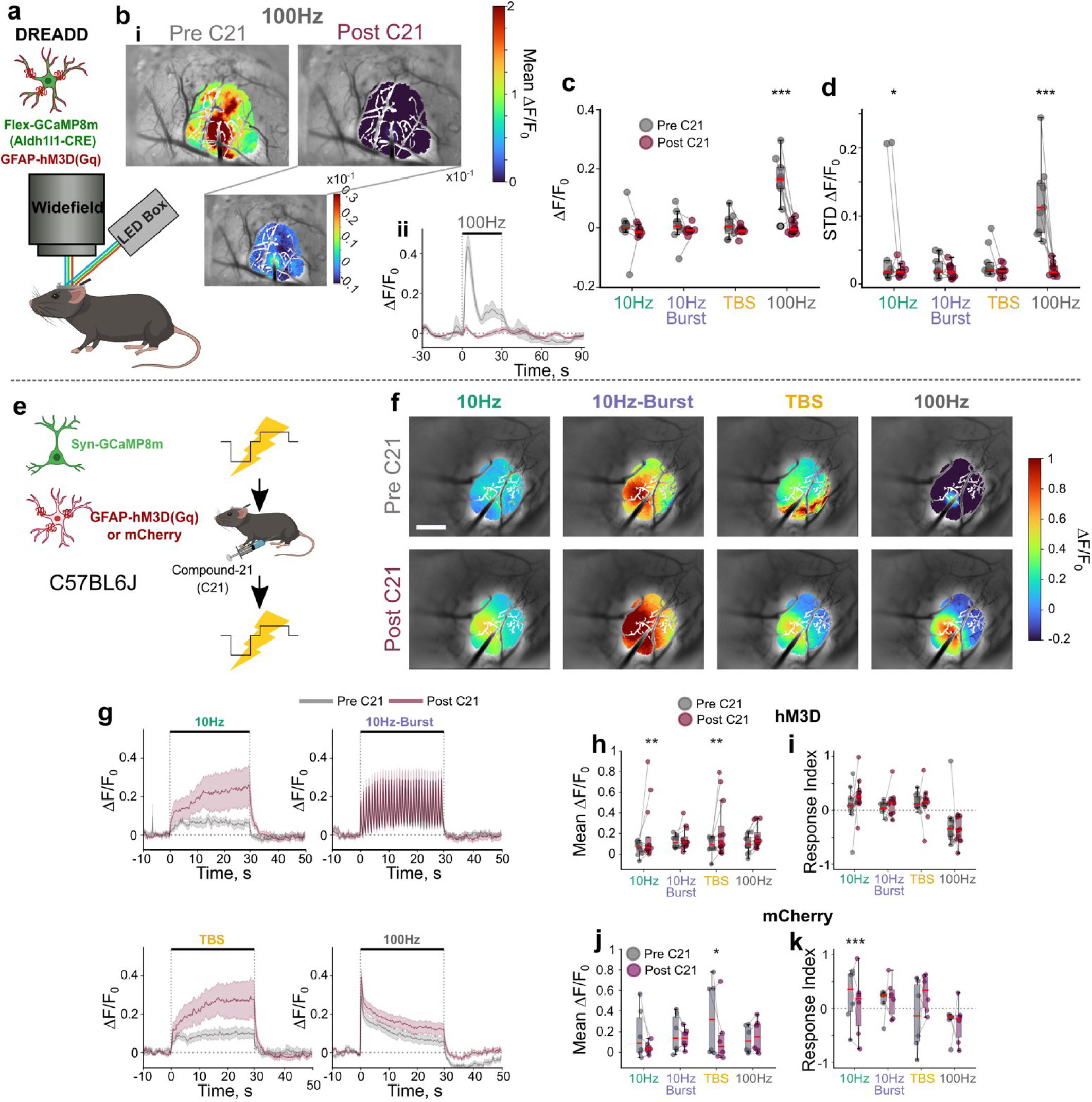
Mesoscale neuron calcium activity facilitates more and depresses less following astrocyte DREADD activation. **A.** Animal model and imaging set up. **B.** i: Representative average projection image of astrocyte calcium activity during 100 Hz stimulation before and after DREADD activation. Inset shows highly localized and smaller activation magnitude ii: 100 Hz stimulation induced astrocyte calcium activity trace before and after DREADD activation. **C.** Average mesoscale astrocyte calcium activity reduces DREADD activation (LMM; Stim: p=1.25e-6, C21: p=4.2e-5, interaction: p=1.5e-5; N=3, 3 trials). **D.** The standard deviation of astrocyte calcium activity during stimulation is significantly reduced following DREADD activation (lm Stim: p=4.8e-4, C21: p=2.8e-5, interaction: p=8.2e-4; N=3, 3 trials). **E**. Animal models and stimulation/DREADD activation experiment setup for neuron quantification. **F**. Representative average projection images highlight that stimulation induced neuronal calcium activity is higher after astrocyte DREADD activation. Scale bar = 500 µm. **G**. Trial averaged traces during different patterns of ICMS demonstrate that the magnitude of neuron activity is increased after DREADD activation, and the activity tends to increase more or depress less after DREADD activation. **H**. The mean neuronal calcium activity during stimulation increased following DREADD activation of astrocytes depending on stimulation pattern (LMM; Stim: p=0.70, C21: p=9.0e-5, interaction: p=0.18). **I**. The response index of neuron calcium activity does not significantly increase after astrocyte dreadd activation (LMM; Stim: p=6.1e-9, C21: p=0.51, interaction: p=0.29). **J**. The mean stimulation induced calcium activity following C21 administration is significantly reduced in validation animals (LMM; Stim: p=0.12, C21: p=0.036, interaction: p=0.16). **K**. The response index of neuron calcium activity significantly reduces after C21 administration in validation animals (LMM; Stim: p=0.08, C21: p=0.26, interaction: p=0.01). **L**. The change in response index is significantly reduced in validation animals compared to DREADD animals (LMM; Stim: p=0.24, C21: p=0.11, interaction: p=0.019). c-d N=3, 3 trials. g-I N= 4 3 trials. J-k N=2 3 trials *: p<0.05, **:p<0.01,***:p<0.0001.

Validation experiments distinguished this effect from changes caused by C21 or repeated stimulation alone. In validation animals, mean neuronal activity decreased following C21 administration (Fig. 4j), and responses shifted toward greater adaptation (lower RI; Fig. 4k). Critically, this validation trend is opposite to the DREADD effect. While repeated stimulation alone drove neuronal activity toward adaptation (lower response index), attenuating astrocyte calcium drove activity up, indicating that the astrocyte-dependent change is not an artifact of repeated stimulation (Fig. S6).

#### 2.2.4 Summary

Together, these results support a model in which astrocytes shape ICMS-evoked neuronal activity through circuit-dependent modulation rather than a uniform excitatory or inhibitory influence. At the single-neuron level, astrocyte attenuation drove bidirectional changes whose direction depended on the initial response regime and whose magnitude increased with distance, where recruitment is increasingly mediated through local synaptic circuitry. Across the mesoscale level, these heterogeneous effects resulted in a net increase in neuronal activity and opposed the trial-dependent attenuation of responses observed earlier. Astrocyte calcium therefore appears to shape the distribution and spatial organization of neuronal responses while constraining the overall network activation during ICMS. We next asked whether enhancing astrocyte calcium activity could produce the opposite effect, suppressing neuronal responses and constraining network activity.

### 2.3 Melanopsin Photostimulation Transiently Amplifies Astrocyte Calcium Activity and Attenuates Neuronal Calcium Activity During ICMS

#### 2.3.1 Amplifying Astrocyte Calcium with Melanopsin

Neuronal adaptation is highest during 100 Hz stimulation. We therefore reasoned that artificially increasing astrocyte calcium elevations like those elicited during 100 Hz stimulation could provide sufficient drive to modulate neuronal activity during ICMS. For this purpose, we selected melanopsin, a Gq-coupled blue light sensitive metabotropic opsin that engages endogenous astrocyte-calcium signaling pathways^58^. Unlike channelrhodopsin, melanopsin does not directly open an ion channel that can elevate extracellular potassium^59^ and alter neuronal excitability and responsiveness to repetitive stimulation^60^. A short 1s photostimulation (10Hz, 30ms pulse width, 1mW) of melanopsin-expressing astrocytes consistently elicited transient astrocyte calcium elevations across repeated trials with similar dynamics to those elicited by 100 Hz ICMS (Fig. 5a-b). Photostimulation in animals lacking melanopsin did not elicit such activity, confirming the response is opsin-dependent (Fig. S11). Astrocytes might have the strongest influence after the initial stabilization of neural activity, but they could also influence the initial onset of activity. Therefore, we tested two delivery timings: photostimulation 1.5s before ICMS (Pre-ICMS) or 5s after ICMS onset (post-ICMS). In both conditions, astrocyte activity was greater than ICMS alone, even during 100Hz stimulation indicating the response was not saturated (Fig.5c, Fig. S11). Validation animals showed little change with photostimulation (Fig. S11). These results demonstrate the feasibility of modulating the gain of astrocyte calcium activity with melanopsin to evaluate their influence on neuronal activity during ICMS.

**Figure 5.**
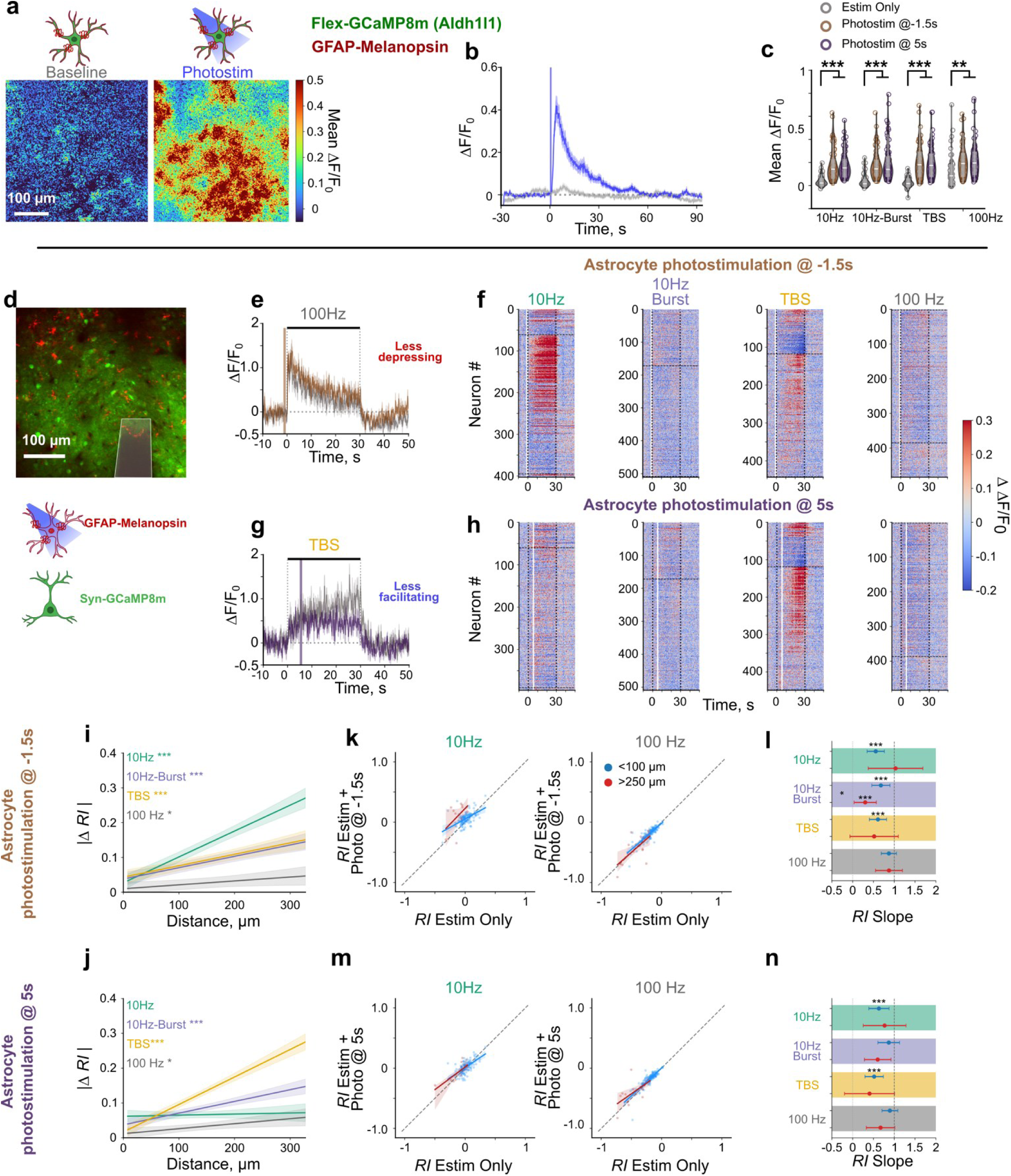
Amplification of astrocyte calcium activity via photostimulation may promote heterogeneous changes in neuronal calcium activity throughout the stimulation duration. **A.** Representative mean projection images of the ΔF/F0 in the 30s after photostimulation of melanopsin expressing astrocytes. **B.** Photostimulation with 1s of 470 nm light delivered at 10 Hz increases astrocyte calcium levels above baseline. **C.** Photostimulation of astrocytes (blue vertical line) increases astrocyte calcium activity during electrical stimulation across patterns (LMM: Stim: p= 1.7e-34, photostim: p= 4.6e-5, interaction p=2.4e-16, N=3 n=65). **D.** (top) Representative image of GCaMP8m expressing neurons (green) and melanopsin expressing astrocytes (red). (bottom) Representative traces of neuronal activity (trial average + standard error) highlights that some neurons will change their response profile over time with melanopsin photostimulation. **E-H.** Photostimulation of astrocytes before electrical stimulation may influence the temporal progression of neuronal activity, where some neurons depress less (E), while others facilitate less (G). Heatmaps of the difference between trial averaged activity of ICMS and ICMS+photostimulation delivered pre-ICMS (F) or post-ICMS (H) show heterogeneous increases and decreases in neuronal activity depending on stimulation pattern. White line indicates photostimulation. **I-J.** The magnitude of change of the response index correlates with distance depending on stimulation condition (LMM I: Dist: p = 3.9e-34, Stim: 0.03, interaction 4.0e-10, N=4, n = 403-511; J: Dist: p= 3.7e-28, Stim: p= 2.4e-5, interaction: p=4.9e-21, N=4, n=396-509). **K-N** Retention of response profile depends on stimulation condition and distance from the electrode. See Fig. S13 (K LMM: Estimonly p = 3.6e-39, Stim: p=8.2e-12, Dist: p = 9.5e-3, Estim only x Stim: p = 3.3e-4, Estim only x Dist p = 0.98, Stim x Dist p = 9.7e-8, Stim only x Stim x Dist: p= 0.01; 10 Hz-prox/dist n=133/11; 10Hz-Burst-prox/dist: n=148/26; TBS-prox/dist: n=135/17; 100Hz-prox/dist: n =148/18; M LMM: Estimonly p = 2.1e-39, Stim: p=3.2e-23, Dist: p = 0.95, Estim only x Stim: p = 0.12, Estim only x Dist p = 0.22, Stim x Dist p = 1.4e-11, Stim only x Stim x Dist: p= 0.42: 10 Hz-prox/dist n=128/11; 10Hz-Burst-prox/dist: n=146/25; TBS-prox/dist: n=141/18; 100Hz-prox/dist: n =146/18). L/N: 95% confidence intervals of slopes in K/M. F=N: N=4. *: p<0.05, **:p<0.01,***:p<0.0001.

#### 2.3.2 Heterogeneous Effects of Astrocyte Photostimulation on Neurons

To test whether transient astrocyte calcium amplification influences neuronal activity during ICMS, we expressed GCaMP8m in all neurons and melanopsin or mCherry in astrocytes (Fig. 5d-n, Fig. S12-13). Photostimulation of astrocytes alone did not increase neuronal calcium activity (Fig. S12). Astrocyte modulation did not globally modulate neuronal activity during ICMS, but there were selective increases during some stimulation parameters depending on when astrocytes were modulated (Fig. S12). These effects were not observed in validation animals, confirming that the changes were due to photostimulation of melanopsin rather than the laser or ICMS alone (Fig. S12). Visual inspection suggested heterogeneous effects in melanopsin-expressing animals, where some neurons showed reduced depression of activity and others showed reduced facilitation, regardless of photostimulation timing. We therefore asked whether astrocytes modulate neuronal responses differently depending on distance from the electrode and ICMS pattern.

To quantify this, we computed a response index across the stimulation train, comparing early versus late activity for each neuron. We then examined whether the magnitude of the change in response index depended on distance from the electrode. Consistent with the DREADD experiments, some patterns influenced the average response index in melanopsin animals (Fig. S12). More importantly, the magnitude of the change increased with distance depending on stimulation pattern and timing of astrocyte modulation (Fig. 5 i,l; Fig. S13). This distance dependence recapitulates the DREADD results and is consistent with the same recruitment framework, in which astrocyte modulation has greater opportunity to influence neurons as recruitment shifts from direct electrical activation near the electrode toward synaptic network recruitment at greater distances. Together, these results indicate that both the timing of astrocyte activity and the mode of neuronal recruitment influence the resulting neuronal response.

We next asked whether the response also depended on how the neurons were driven within the circuit, as it did when astrocyte responses were attenuated. Again, we regressed each neurons RI with astrocyte photostimulation, against its RI with ICMS alone and evaluated if this relationship changes with distance (Fig. 5k-n, Fig. S13). Deviations from the unity line indicated that the initial response profile of a neuron was associated with how astrocyte modulation influenced the timing of its response. Consistent with chemogenetic modulation (Fig.3), 100 Hz showed limited deviation when astrocytes were transiently modulated. Conversely, the effect depended on other stimulation parameters and timing of photostimulation delivery, with weaker influence on neurons farther from the electrode (Fig. 5 k-l, Fig. S13). This weaker response at greater distance may be due to the limited number of neurons ≥250 µm. Fitting this slope continuously over distance did show non-linear changes with distance from the electrode (Fig. S13). Furthermore, we did observe heterogeneous changes in the classification of the response profile based on response index depending on stimulation pattern and timing of astrocyte modulation (Fig. S13). These distance-dependent effects prompted us to next ask whether astrocyte modulation would produce a corresponding effect at the network level during mesoscale imaging.

#### 2.3.3 Mesoscale Imaging Confirms Transient Astrocyte Modulation of Neuronal Networks

Two-photon imaging captures neurons near the electrode, a mix of directly and synaptically recruited populations, making it difficult to separate photostimulation effects from plasticity. To examine network-level effects, we performed mesoscale imaging (Fig. 6a-j). One important limitation is that the single-photon excitation wavelength used to excite GCaMP also activates melanopsin. Despite using lower illumination power to image calcium responses, baseline imaging caused some increase in astrocyte calcium prior to ICMS (Fig. 6a-b, Fig. S14). Nevertheless, photostimulation was still able to transiently elevate astrocyte calcium above ICMS levels within 5s of photostimulation for some stimulation parameters (Fig. 6b-d). Validation animals showed no effect of photostimulation in the absence of melanopsin (Fig. S14).

**Figure 6.**
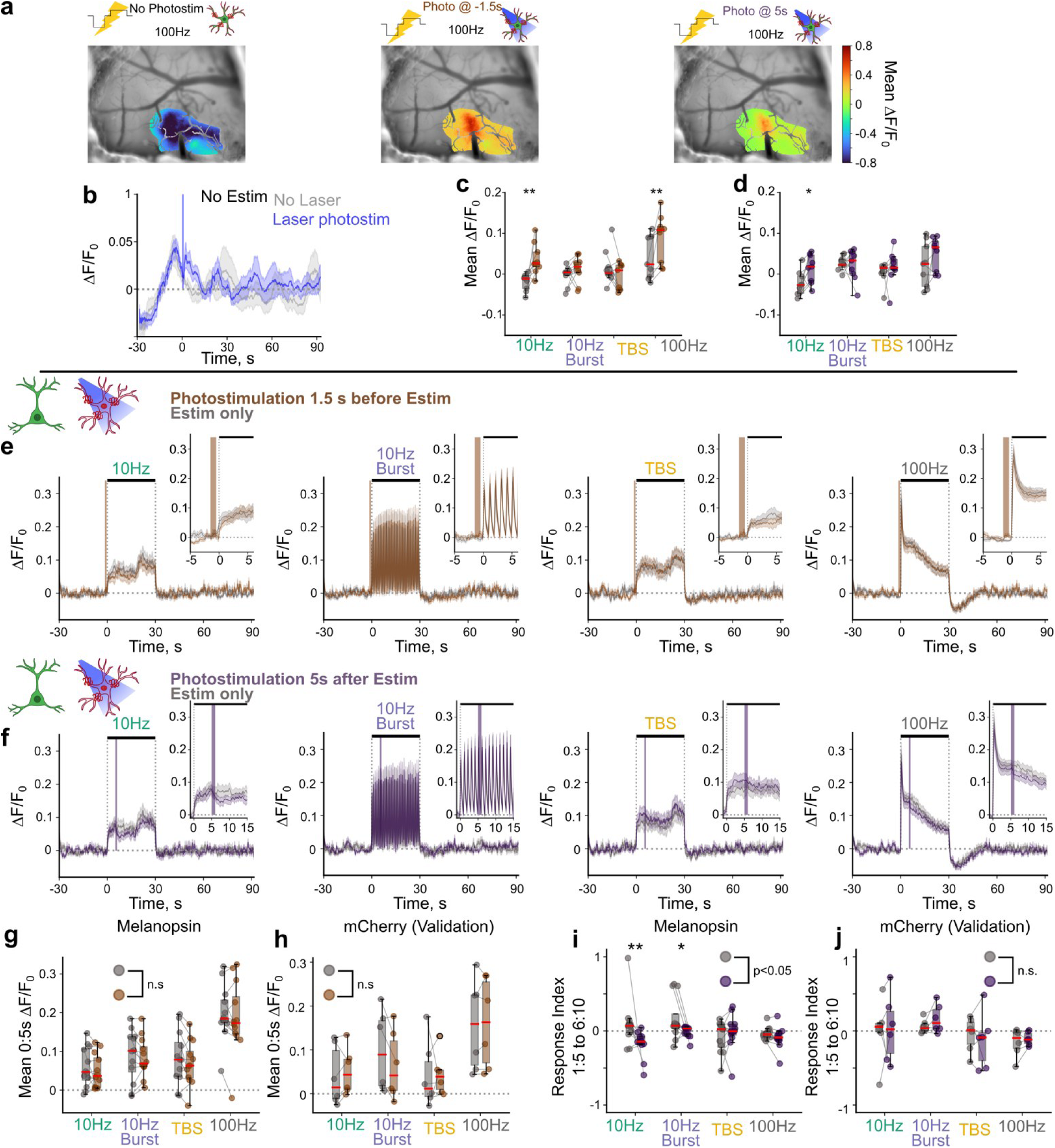
Photostimulation of astrocytes transiently decreases mesoscale neuronal activity within 400 µm. **A.** Representative images of the mean stimulation induced astrocyte calcium activity with and without laser photostimulation. **B**. Imaging of GCaMP excites melanopsin and increases astrocyte calcium during baseline, but laser stimulation (blue vertical line) still transiently increases astrocyte calcium. **C**. Laser photostimulation increases astrocyte calcium activity within the 5s of stimulation onset when delivered 1.5s before electrical stimulation onset (LMM: Stim: p=9.9e-7, Photo-pre: 4.9e-3, interaction: p= 6.1e-3, N=3, 3 trials). **D**. Laser photostimulation increases astrocyte calcium activity within 5s of photostimulation when delivered 5s after stimulation onset (LMM: Stim: p=3.9e-3, Photo-post: 0.019, interaction: p= 0.25, N=3, 3 trials). **E.** Average traces ± standard error show neuronal calcium activity is comparable when astrocyte photostimulation is delivered before stimulation onset. **F**. Neuronal calcium traces show that astrocyte photostimulation transiently decreases neuronal calcium activity when delivered 5s after stimulation onset. **G-H**. Astrocyte photostimulation before electrical stimulation does not affect the mean neuronal calcium activity during the first 5s of electrical stimulation for melanopsin (**G**; LMM: Stim: p=1.2e-25, Photo-pre: 0.10, interaction: p= 0.97) and validation (H; LMM: Stim: p=6.5e-9, Photo-pre: 0.72, interaction: p=0.44) animals. **I-J**. The response index between 1-5s of electrical stimulation onset and 6-10s after photostimulation is significantly decreased with photostimulation of astrocytes in melanopsin (I; LMM: Stim: p=0.09, Photo-post: 0.01, interaction: p= 0.04) animals, but not validation (J; LMM: Stim: p=0.17, Photo-post: 0.74, interaction: p= 0.90). N= 4, 3 trials for melanopsin; N=2, 3 trials for validation. *: p<0.05, **:p<0.01,***:p<0.0001.

We then examined how this transient astrocyte calcium elevation modulated neuronal activity during mesoscale imaging. When examining the overall traces of neuronal calcium activity with astrocyte photostimulation, we did not observe a broad change in the activity over the 30s ICMS train (Fig. 6e-f). However, detailed analysis revealed that photostimulation delivered 5 s after ICMS onset caused a transient decrease in neuronal activity, particularly for 10 Hz and 10Hz-burst stimulation (Fig. 6f insets). To quantify this effect, we computed the response index comparing the activity just after the onset of stimulation (1-5s) to the activity after the photostimulation (6-10s). Neurons in melanopsin animals showed a significant reduction in response index following photostimulation, while validation animals showed no effect (Fig. 6i-j). Thus, transient amplification of astrocyte calcium was sufficient to produce a short-lived reduction in neuronal activity at the network level.

#### 2.3.4 Summary

Together, these experiments show that astrocyte calcium can modulate neuronal activity during ICMS, with the direction and magnitude of this modulation depending on stimulation pattern, timing, neuronal response state, and distance from the electrode. Astrocyte calcium suppression preferentially altered responses of more distant neurons, consistent with greater modulation of synaptically recruited populations^61^, while transient astrocyte calcium amplification produced a short-lived reduction in neuronal activity at the network level. These effects were modest relative to the dominant influence of ICMS, indicating that astrocytes bias rather than determine neuronal responses during stimulation. The findings therefore support a modulatory role for astrocyte calcium in shaping the temporal dynamics of ICMS-evoked neuronal activity, while leaving the dominant mechanisms of perceptual adaptation unresolved.

## 3 Discussion

While electrical neuromodulation is a cornerstone of neurotherapeutics^2,9,62,63^, neural prostheses^4,14,64^, and circuit mapping^65^, therapeutic optimization remains limited by high response variability across stimulation paradigms^4,15^. One potential contributor to this variability is the neuron-centric view of electrical stimulation^19^, which focuses on the direct effects of stimulation on neurons while giving less consideration to glial responses that may shape neural activity over the same timescale^25^. Here, we show that astrocyte calcium activity modulates neuronal responses during ICMS in awake mice.

We first found that astrocytes showed heterogeneous calcium responses that depend on stimulation pattern and persist on a timescale compatible with modulation of ongoing neural activity (Fig. 1-2). Causal manipulation then showed that this modulation is neither simply excitatory nor inhibitory. Attenuating astrocyte calcium did not uniformly raise or lower neuronal activity. Instead, it compressed the population toward an intermediate response state, with initially facilitating and initially adapting neurons converging toward that state from opposite directions (Fig. 3-4). This shift was opposite to the adaptation observed in validation animals. Raising astrocyte calcium produced its own pattern- and timing-dependent changes in neuronal responses (Fig. 5-6). The direction and magnitude of these effects varied with stimulation pattern, timing, each neuron’s initial response state, and its distance from the electrode. Furthermore, the largest effects occurred in distal neurons, which are more likely to be recruited through synaptic activity than by direct electrical activation.

Together, these findings suggest that astrocyte calcium does not provide a fixed inhibitory or homeostatic tone. Instead, astrocyte calcium appears to gate how far neuronal responses are drawn toward an intermediate state. Astrocyte state may therefore contribute to the response variability that currently limits stimulation control, rather than simply representing unexplained noise.

### 3.1 Astrocytes reactively sense and actively shape evoked neural activity

Despite early speculation^66^, many studies show that astrocytes signal through calcium and, in doing so, release substances that act on neurons, a process called gliotransmission^32,41,52,67–71^. A single astrocyte can release different substances depending on the pattern and duration of neural activity^32^, as well as can act selectively on differentneuronal subtypes^12,70,72^. Therefore, if astrocytes shape ongoing nearby neural activity during stimulation, their calcium should rise and fall in step with the neural activity they are shaping.

We found that astrocytes showed large and widespread calcium elevations, with compartmental activity following neuronal activation, suggesting they are being recruited through synaptic activity rather than direct depolarization, which has not been observed directly *in vivo*^38,39^. Astrocytes therefore appear to respond after neurons are activated, rather than being directly activated by the electrical stimulus itself. The delay could also partly reflect slower calcium handling in astrocyte processes than in neurons^73–75^. Astrocyte calcium usually began near the electrode and spread outward as a wave, most clearly during 100 Hz stimulation (Fig. 1), matching our earlier finding of high glutamate within about 20 µm of the electrode during 100 Hz stimulation^76^. Earliest calcium activity appeared in the fine cloud-like processes rather than in the cell body (Fig. 1), and other work reports the strongest astrocyte responses in these process-rich regions^39^, together pointing to perisynaptic processes as the primary site of stimulation-evoked astrocyte calcium and the likely locus of synaptic modulation. These processes closely appose both presynaptic terminals, where neurotransmitters are released, and postsynaptic membranes, where those signals are received, forming the third component of the tripartite synapse. This anatomical arrangement places astrocytes at the site of neuronal communication, where they can detect synaptic activity and potentially modulate its transmission.

Astrocyte calcium also covered an area as large as, or larger than, the area of active neurons, rose and fell with neuronal activity, and was weighted toward the start of the train. Astrocytes also remained above threshold longer than neurons after stimulation ended (Fig. 2). During the first three trials, when astrocyte responses were strongest, the duration of this post-stimulation activity tracked the duration of neuronal suppression (Fig. 2i, Fig. S6). This association weakened over repeated trials as astrocyte responses desensitized (Fig. S6), suggesting it depends on the strength of the astrocyte response rather than being a fixed feature of the network. Because astrocyte activity outlasted the stimulus and neuronal activity, it may contribute to the short-term circuit changes that persist after stimulation ends. Taken together, this prolonged integration may allow astrocytes to shape the post-stimulation network state and contribute to post-stimulus depression^77^.

We further tested this with a second, independent manipulation. Raising astrocyte calcium with melanopsin also changed neuronal responses during ICMS, producing heterogeneous, distance-dependent shifts that recapitulated the pattern seen when astrocyte calcium was lowered (Fig. 6). These changes did not appear in validation animals expressing mCherry in place of melanopsin, confirming they depended on the astrocyte manipulation rather than on the laser or stimulation alone. Because two different tools that both change astrocyte calcium produced related changes in neuronal responses, while the inactive control did not, these effects are unlikely to reflect an off-target action of either tool. Together, the timing of astrocyte and neuronal activity shows that astrocytes respond to evoked neural activity on a timescale that could allow them to influence it. The causal manipulations then show that changing astrocyte calcium changes the resulting neuronal activity. Thus, astrocytes do not simply follow evoked neural activity, they can also shape it.

### 3.2 Astrocyte modulation of neural activity operates differentially across spatial scales

If astrocytes modulate neural activity during stimulation, where they act should depend on how neurons are recruited. Electrical stimulation elicits both direct (antidromic), and indirect (transsynaptic/orthodromic) recruitment of neurons^78^, with direct activation of axons presumed to occur within tens of microns of the electrode^79,80^. Consequently, activated neurons within ∼100 µm are presumed to have a higher probability of being recruited through direct engagement, while neurons ≥250 µm are more likely activated orthodromically^42^. While modeling suggests that synaptic input can influence how well a neuron’s cell body follows the stimulation^81^, it is unclear whether astrocytes exert sufficient drive to overcome the strength of direct electrical engagement, and therefore are expected to have a stronger influence on the neurons recruited through synaptic mechanisms.

This distinction provides a simple framework for interpreting the spatial pattern of our result. In this framework, neurons near the electrode are dominated by the imposed electrical (antidromic) drive, so astrocyte calcium there mostly reflects a response to neural activity rather than a way of controlling it. Astrocyte influence appears at greater distances, where neurons are recruited synaptically and synaptic transmission, the site where astrocytes act, dominates. We further speculate that the direction of this influence may itself shift with distance, with synaptically recruited neurons farther from the electrode responding differently from the directly driven neurons nearby, although testing the idea would require resolving the effect by distance and direction in future work. The recruitment difference also accounts for the divergence between the tightly coupled local two-photon signals near the electrode and the weaker coupling seen at mesoscale distances, since the two regimes sample directly driven versus synaptically recruited populations.

Under mesoscale imaging, which pools the signal from neurons across multiple cortical layers and can image large distances from the electrode^57^, astrocyte calcium shaped neuronal activity across the stimulation train in a direction determined by the manipulation. Lowering astrocyte calcium increased neuronal activity over time, opposite to the adaptation seen across repeated stimulation in validation animals receiving stimulation alone (Fig. 4; consistent with stimulation-driven desensitization,^39^). In contrast, raising astrocyte calcium transiently decreased neuronal activity (Fig. 6). Additionally, under these manipulations, individual neurons farther from the electrode showed greater changes in the temporal progression of their activity (Fig. 3,5). However, these changes did not simply reflect a global stabilizing effect across the population of local neurons as was observed in mesoscale imaging. This difference between local and mesoscale measurements is informative, because astrocyte calcium spread beyond the neuronal activation zone (Fig. 2), placing astrocytes in a position to influence surrounding synaptically recruited neurons, which is the distal region where their effects on neuronal response were strongest.

Near the electrode, responses look heterogeneous, yet across the network they sum to a consistent shift. These two observations reflect the same mechanism. Astrocytes act on synapses in a graded, cell-type-specific way, releasing different signaling molecules depending on the pattern and duration of neural activity and weakening nearby synapses that were not themselves active. This graded action does not mean astrocytes change every neuron in the same direction. Over a long stimulation train this signaling would push strongly facilitating and strongly adapting neurons toward the same middle ground, refining the balance of excitation and inhibition^61^. Cell-type-resolved recordings offer a candidate circuit basis for this distance dependence. Somatostatin (SOM) interneurons, which target distal dendrites, are preferentially recruited farther from the electrode during burst and theta-burst patterns, unlike other subtypes, and can disinhibit pyramidal neurons through SOM-to-PV interactions. Like astrocytes, these interneurons act as second-order modulators on the synaptically recruited distal zone rather than on directly driven neurons near the electrode. Astrocyte modulation of these distal, SOM-associated synapses would shift the balance of excitation and inhibition according to each neuron’s starting state.

In many cases the direction of the effect depended on a neuron’s initial response during baseline ICMS. That dependence is the core of the finding. Rather than uniformly raising or lowering activity, astrocyte calcium sets how far each neuron’s response is pulled toward a middle state, and the direction depends on whether the neuron started out excitation-dominated or adaptation-dominated. This regime-dependent compression, rather than a simple net increase or decrease, is what separates astrocyte modulation from a uniform tone. Acting selectively on individual neurons near the electrode, these local changes sum into a broad, directional shift across the wider network that may contribute to perceptual fading^15^.

### 3.3 Astrocyte influence depends on stimulation pattern

The efficacy of therapeutic neurostimulation^9^, and the quality of evoked percepts in cortical neuroprostheses^82^ depends on the pattern of stimulation. For example, a bursting paradigm was recently shown to elicit comparable improvement of ataxia in a mouse model of Parkinson’s disease as traditional continuous high-frequency deep brain stimulation, but with persistent relief after termination of stimulation^9^. This finding indicates that consideration of circuit level plastic effects could improve therapeutic outcomes of neuromodulation. Owing to their well-documented role in regulating synaptic plasticity, it is important to understand how astrocytes respond to different stimulation patterns, and whether their influence on neural activity is pattern-dependent.

Our results demonstrate that stimulation-induced astrocyte calcium dynamics are indeed pattern dependent, where the timing, magnitude, shape, and spatial scale are all modulated by stimulation pattern (Fig. 1-2). By lowering and raising astrocyte calcium during ICMS, we showed that the astrocyte contribution to the time course of neural activity critically depends on how stimulation is delivered (Fig. 4-6). Transiently raising astrocyte calcium either at the onset of stimulation or after neural activity began to stabilize (Fig. 5-6) changed different aspects of the neural response. The astrocyte effect was clearest during 10-Hz and TBS patterns and weakest during 100-Hz stimulation.

The weak influence during 100-Hz stimulation suggests that the strong decrease in activity at this pattern is driven mainly by the neurons themselves, through failure to keep pace with the pulses, increased inhibition, or depletion of neurotransmitters. Therefore, the 100-Hz condition serves as a built-in control for how specific the effect is. Because astrocyte manipulation cannot move a response that is already saturated by the neuron-intrinsic adaptation, the modulation we observe at lower frequencies is unlikely to be an off-target action of the manipulation itself. Instead, they act through the synaptic processes that dominate when intrinsic adaptation does not. When these neuron-intrinsic limits are absent, the longer-lasting astrocyte calcium seen during lower-frequency and bursting paradigms (Fig. 1-2) could provide sufficient drive to actively modulate ongoing neural activity^83^. Trial data point the same way. Astrocyte calcium was better sustained across repeated trials under bursting patterns, and neuronal responses adapted least under those same patterns. Here, astrocytes may be one reason bursting protocols preserve responses over repeated stimulation, offering a candidate glial contribution to the reduced perceptual fading reported for bursting paradigms^17,18^. Ultimately, these results suggest that optimizing the therapeutic efficacy of neuromodulation will require designing stimulation patterns that take these neuron-glia dynamics into account.

### 3.4 Limitations

Several methodological limitations must be considered to conceptualize our findings. First, spectral overlap between GCaMP emission and jRGECO excitation could confound our correlational analyses, though evoked activity profiles were consistent with other models and previous work. More broadly, calcium is an imperfect proxy for astrocyte function. Many of the pathways through which astrocytes shape synaptic transmission, including glutamate uptake, D-serine and ATP/adenosine release, and potassium buffering, are only partially reflected in, or can occur independently of, the somatic and microdomain calcium transients captured by GCaMP8m^84,85^. Consequently, the evoked calcium signal should be read as an indicator of astrocyte engagement rather than a complete readout of astrocyte influence, and attenuating this signal reduces, but may not fully abolish, calcium-independent astrocytic contributions to network activity. Combining calcium imaging with direct measurements of transmitter release, uptake, and extracellular potassium, would help distinguish the mechanisms underlying astrocytic modulation of stimulation-evoked activity. Stimulation of Gq DREADDs in astrocytes leads to a period of increased astrocyte calcium oscillations followed by a sustained elevation in baseline calcium lasting hours that limits further transient elevations to stimulation. Future experiments could determine how these distinct phases influence gliotransmission and other astrocyte functions. Because sustained intracellular calcium can decouple astrocyte gap junctions^86^, the increased excitation we observed (Fig. 4) may stem from impaired potassium buffering^87^, attenuated gliotransmission, or both. Either mechanism is consistent with our central finding that astrocyte calcium shapes stimulation-evoked neuronal responses. Resolving these alternatives would refine the mechanisms underlying that finding. Finally, because melanopsin activation can induce lasting changes in neural activity *in vitro*^58^, tracking baseline network excitability and recovery between trials would help distinguish effects of ongoing activity from modulation of subsequently ICMS responses.

## Conclusion

To our knowledge, this is the first study in awake animals demonstrating that astrocytes causally modulate the brain’s response to electrical stimulation. Instead of simply following neural activity, astrocytes shape it, pulling neuronal responses toward an intermediate state in a way that depends on stimulation pattern, timing, and each neuron’s starting state. These findings place astrocytes alongside neurons as active contributors to the cortical response to stimulation, and argue for expanding the neuron-centered view of clinical neuromodulation. Building these neuron-glia dynamics into how stimulation parameters are chosen may help design more effective neurotherapeutic stimulation.

### 4.0 Materials and Methods

#### 4.1 Animals

Animal procedures and care were conducted in accordance with federal guidelines and were approved by the Institutional Animal Care and Use Committee at the University of Pittsburgh. All procedures were performed in adult male or female mice (n=25, >8 weeks of age, 23-35g). Transgenic mouse strains were obtained from Jackson Laboratory (Bar Harbor, Me): Aldh1l1-CRE (023748, for measuring astrocyte calcium activity), C57BL/6J (000664, for measuring neuronal calcium activity). Specific animal numbers for each experimental group are listed in Sup Table 1 and figure legends. Mice were housed in 12h/12h light/dark cycle at constant room temperature and humidity with free access to food and water.

**Table 1.** AAVs used to generate animal models.

| Virus Name | Source |
| --- | --- |
| Syn-NES-jRGECO1a.WPRE.SV40 (AAV1) | Addgene |
| Syn-jGCaMP8m-WPRE (AAV1) | Addgene |
| CAG-FLEX-jGCaMP8m-WPRE (AAV1) | Addgene |
| GFAP104-Melanopsin-mCherry (AAV8) | UNC Vector Core |
| GFAP-hM3D(Gq)-mCherry | Addgene |
| GFAP104-mCherry | Addgene |

Experimental animals were divided into 3 groups. Characterization animals were Aldh1l1-Cre mice expressing Syn-jRGECO1a and Flex-GCaMP8m, permitting simultaneous imaging of neuronal and astrocyte calcium activity. DREADD animals expressed GFAP-hM3D(Gq)-mCherry for chemogenetic manipulation of astrocyte activity, with astrocyte calcium images in Aldh1l1-Cre mice (Flex-GCaMP8m), and neuronal calcium imaged in C57BL/6J mice (Syn-GCaMP8m). Melanopsin animals similarly expressed GFAP-Melanopsin-mCherry for optogenetic manipulation of astrocyte activity, with astrocyte and neuronal calcium images in separate Aldh1l1-Cre and C57BL/6J cohorts respectively. For DREADD and melanopsin experiments, validation animals expressed GFAP-mCherry instead of the active construct and underwent identical experimentation. Within-subject pre-manipulation measurements served as the primary control for effects of astrocyte manipulation on neuronal activity.

#### 2.2 Cranial Window Surgery and Virus Injection

Cranial window preparations are described in detail in previous publications^43,88,89^. Briefly, animals were initially anesthetized using an intraperitoneal (IP) injection of ketamine/xylazine (75 mg/kg; 7 mg/kg) and updated with ketamine (40 mg/kg) as needed throughout the procedure. Additionally, mice received a subcutaneous injection of 20% mannitol in PBS to limit brain swelling^90,91^. Pain was managed with an IP injection of ketoprofen (5 mg/kg) during surgery and for 2 days following the initial cranial window surgery. Following head-fixation, a custom stainless steel rectangular frame was cemented to the skull, two stainless steel bone screws were installed anterior to bregma, and 2 circular craniotomies (∼3 mm in diameter) were created over the left and right primary visual cortex (V1 1 mm anterior to lambda, ±3 mm lateral).

Viruses, and sources are listed in Table 1. All viruses were diluted to a final concentration of 5e12 in 20 % mannitol in PBS to aid effective spread^90^, with a small amount of sulforhodamine 101 to confirm injection location^20,79^. Viruses were delivered using pressure injection (Picospritzer, <10 psi) through a glass pipette (2-4µm in diameter) in 2-3 sites (<1 µl total) in both hemispheres. Following injection, the craniotomies were sealed with a glass coverslip cemented to the skull.

#### 2.3 Electrodes and Implantation Surgery

Custom, 4-channel (Platinum/Iridium) single-shank (50 µm spacing, 702 µm2 diameter) electrodes were obtained from NeuroNexus (NeuroNexus Technologies, Ann Arbor, MI). Electrodes were ‘activated’ to increase charge storage capacity and site impedance was confirmed to be below 500 kOhm^79^. Electrode implantation procedures were performed following habituation to head fixation and ≥ 4 weeks following the initial surgery. Mice were initially anesthetized with 1.5-2% isoflurane and maintained at 1% isoflurane throughout the procedure. After removing the glass coverslip of one hemisphere, an electrode was implanted into the virus expressing area at a 30° angle at a speed of 220 µm/s to a final depth of 180-260 µm below the cortical surface using an oil hydraulic Microdrive (MO-81 Narishige, Japan). Ground and reference wires were wrapped around the bone screws, and the window was sealed with Kwik-Sil (World Precision Instruments, Sarasota, FL) and a glass coverslip. Animals recovered for at least 1h with access to food and water before stimulation experiments.

#### 2.4 Electrical Stimulation

Electrical stimulation was performed in current-control mode using a TDT IZ2 stimulator controlled by an RZ5D system (Tucker-Davis Technologies, Alachua, FL). Stimulation consisted of 30s trains of cathodic-leading, charge-balanced pulses matching those used clinically (200 µs cathodic, 100µs interphase, 400 µs return, 10 µA, 2nC/phase) and within established safety limits for microstimulation^92^. Four stimulation paradigms were delivered: 100 Hz-uniform which acted as a control case with high neuronal adaptation, 10 Hz-uniform stimulation, which produces limited adaptation, and two bursting paradigms with an average frequency of 10 Hz matching the pulse count with 10Hz-uniform. The 10 Hz-burst paradigm consisted of a 100 ms burst of 10 pulses at 100 Hz followed by a 900 ms delay, while the theta-burst stimulation (TBS) paradigm consisted of bursts of 2 pulses delivered at 100 Hz with a 200 ms inter-burst interval. TBS was designed to approximate theta-burst paradigms used in repetitive transcranial magnetic stimulation, which is thought to promote plasticity^93^. 10 Hz burst stimulation was designed to evoke brief periods of high neuronal activity separated by recovery intervals. Paradigms were delivered in random order for 2-6 trials during two-photon and mesoscale imaging. Trials were separated by at least 2 minutes to limit short-term plasticity^22^, and consisted of a 10-30s pre-stimulation baseline, a 30s stimulation, and a 20-60 post-stimulation period, for a total image duration of 80-120s.

#### 2.5 Two-Photon Calcium Imaging

For all imaging experiments, animals were head-fixed on a custom treadmill during imaging. A two-photon laser scanning microscope (Bruker, Madison, WI) equipped with a 16x water immersion objective (0.8 NA, Nikon Inc, Tokyo, Japan) was used for population-level imaging of neuronal and/or astrocyte calcium activity. An OPO laser (insight Ds+; Spectra-Physics, Menlo Park, Ca) tuned to 920 nm was used to image GCaMP and an ultrafast laser at 1040 nm was used to image jRGECO1a. Total laser power was kept at or below 25 mW to limit photobleaching and heating of the electrode. Imaging was performed in resonant mode over a field of view of 413 x 413 µm at either 15 Hz (1024 x 1024 pixels, 0.403 µm/pixel) for high resolution imaging of astrocytes or at 30 Hz (512 x 512 pixels,0.806 µm/pixel) for imaging of neuronal activity.

#### 2.6 Mesoscale Epifluorescence and Hemodynamic Imaging

Mesoscopic widefield imaging of signals was conducted with a fluorescence microscope (MVX-10, Olympus Inc, Japan) over a field of view (256 x 344 pixels) of either 2.1 x 2.75 mm (5x zoom) or 3.2 x 4.3 mm (3.2x zoom) window, which permitted observation of the entire cranial window and regions outside the window. Three sequentially interleaved, bandpass filtered LEDs enabled pseudo-simultaneous acquisition of calcium and hemodynamic signals. Blue light (482.5 ± 15.5 nm #67-082 Edmund Optics Inc, NJ, US) was used to excite GCaMP, Lime light was used to excite jRGECO1a, Green light (532 ± 2 nm FLH532-4, Thorlabs Inc, NJ, US) was used for hemoglobin-sensitive imaging, and Red light (633 ± 2.5 nm; FLH633-5, Thorlabs, NJ, US) for deoxy-hemoglobin sensitive imaging. These two hemodynamic readouts were then used to extract oxy- and deoxy-hemoglobin changes evoked by ICMS. Acquisition of calcium and hemodynamic activity in animals expressing GCaMP8m in astrocytes and jRGECO1a was conducted using Blue, Lime, and Green light with a notch filter placed in the imaging path before the camera (QImaging Retiga R1; Cairn Research Ltd, Kent, UK) to block the excitation of GCaMP and jRGECO1a. In all other animals, Cyan, Green, and Red light were used with a 500 nm long-pass filter to block GCaMP excitation. Imaging was controlled using Micro-Manager software for a 30 Hz frame rate, resulting in an effective sample rate of 10 Hz for each signal.

#### 2.7 Optogenetically-combined Electrical Stimulation

Optogenetic stimulation of melanopsin was performed using a TTL-controlled 471 nm DPSS laser (LaserGlow, Canada) through an optical fiber (S405XP, ThorLabs Inc, NJ USA) with a core diameter of 125 µm placed on the coverglass targeting the electrode site being electrically stimulated^94^. High speed shutters covering the photomultiplier tubes (PMT) in the two-photon system were shut during photostimulation. Photostimulation consisted of a pulse width of 30 ms pulse trains delivered at 10 Hz for 1s at 1mW for all experiments combining optical and electrical stimulation. Light was delivered either 1.5s before or 5s after electrical stimulation began. These two paradigms let us determine the effect of astrocyte activity to the onset of electrical stimulation induced neural activity, or the effect on neural activity after the initial stabilization of neural activity.

#### 2.8 DREADD Activation

The DREADD agonist compound-21 (C21) was used to activate the Gq DREADD HM3d expressed on astrocytes due to its longer half-life and to reduce the potential of off target effects from CNO administration^95^. Following stimulation trials under the MVX and two-photon, mice received an IP injection of 1 mg/kg C21. Post-administration stimulation began 30 minutes after delivery of the agonist, which is consistent with the timing for astrocyte calcium signaling to reduce, potentially by exhausting astrocyte calcium stores.

### 2.9 Data Processing

All data processing and analysis were performed using custom written scripts in Python (Ver. 3.10, Python Software Foundation DE, US) with existing modules, with the exception of two-photon image registration being performed using Suite2p^96^.

#### Motion Correction and Region of Interest Identification

Two-photon image registration was performed using phase correlation methods in Suite2p with non-rigid registration. Mesoscale image registration was performed using custom scripts that also used phase correlation for rigid registration. Images were registered to a common image across all stimulation trials within a session. ROI identification of neurons was performed manually by inspecting temporally binned images and standard deviation projected images in FIJI (ImageJ, NIH^97^), using the oval ROI feature to create an ROI map. The ellipse parameters were imported into Python to determine the corresponding pixel locations to each neuron. Pixels corresponding to multiple ROIs were excluded from analysis.

#### Astrocyte ROI Identification

GCaMP8m expression in astrocytes provided a strong baseline fluorescence within the soma, main branches, and surrounding cloudy regions (Fig. S1). We developed a semiautomatic segmentation algorithm to classify these compartments, (Fig. S1) similar to CalSee^33^. Astrocytes with clearly identifiable somata and main branches were selected manually in FIJI, and the soma centroid was defined for subsequent segmentation. Mean projection images across trials were processed with a white-tophat filter to reduce background fluorescence and multiscale Gaussian filtering followed by Hessian feature extraction to enhance astrocyte morphology. A weighted image combining the white-tophat output (60%) and Hessian feature map was then locally thresholded using a Sauvola filter to segment the soma and main branches. This segmentation was combined with a manually defined astrocyte region mask to distinguish main branches from the surrounding cloudy region. To identify the soma, the astrocyte mask was applied to the weighted image, and a Sobel filter was used to detect its boundary. The soma boundary was defined as the point where radial bin fluorescence intensity fell below 50% of the maximum, which reliably delineated the soma across astrocytes. Branch and cloudy regions were further subdivided into ∼3×3 µm domains for analysis

#### Neuron ROI Identification

Neuronal soma regions were manually outlined using the oval tool in FIJI as previously described^43,79,88,89^. Potential neurons were identified by cross referencing mean and standard deviation projection images with movies downsampled to 3 Hz. The positions of the somata and fitted ellipse parameters for each ROI were extracted to create a ROI map within python.

#### Two-Photon Image Processing

Mean fluorescence was extracted from each neuronal or astrocyte ROI for every frame and converted to ΔF/F0, where F0 was the mean fluorescence during the pre-stimulation baseline. Traces were median filtered (7-frame kernel) and then low-pass filtered using a 3^rd^ order 5 Hz FIR filter. Photostimulation shutter artifacts were masked during filtering to prevent contamination of adjacent time points. ROIs without stimulation-dependent calcium activity in any manipulation were excluded using an ROI-specific activation threshold. Baseline values >2SD above the mean were first removed, after which the baseline mean and SD were recalculated. The activation threshold was calculated as the new mean + 3SD, as previously described (Fig. S2). An ROI was considered active when its ΔF/F0 exceeded its threshold for at least 500 ms for neurons, or 1s in astrocyte domains during stimulation. Activated astrocyte domains within each domain class (e.g. branch, cloudy region) were averaged to generate a fluorescent trace for that compartment (Fig. S2).

#### Mesoscale Image Processing

Mesoscale calcium activity was quantified in 20 µm radial bins centered on the peak activation site, which was defined manually from the mean 100 Hz stimulation image as the pixel with the highest intensity^43,88,89^. Calcium emission and green-light reflectance signals were extracted from each bin, and low-pass filtered using FIR filters with a 4.5 Hz and 4 Hz cutoffs, respectively. Signals were normalized to the pre-stimulation baseline as F/F0 and R/R0, and the GCaMP fluorescence was corrected for hemodynamic effects by dividing F/F0 by R/R0, as previously described^98^, before conversion to ΔF/F0.

Spatial and temporal activation were determined using the thresholding approach described above. A bin was considered active when its fluorescence exceeded threshold and all bins closer to the electrode were simultaneously active, as previously done^89^. Analyses were restricted to the virus-expression region while excluding blood vessels using a computationally generated mask (Fig. S3).

### 2.10 Quantification and Statistical Analysis

#### Quantification metrics

To compare temporal activation dynamics between astrocytes and neurons, we quantified activation latency, “activation time”, and time above threshold. Activation latency was defined as the first time an ROI, or radial bin exceeded its activation threshold for at least 300 ms. The activation time was defined as the time required for a signal to reach 50% of its cumulative activity during the 30s stimulation period, as previously described (Fig. S2) with values < 15s indicating activity was weighted toward stimulation onset, and values > 15s indicating activity weighted toward the end of stimulation. Changes in neuronal activity over time were also quantified using a response index (*RI*), calculated as the normalized difference in mean activity magnitude between two time windows, where (a) and (c) represent the first period of the earlier, and later analysis time windows, respectively. *RI* ranges from -1 to 1, with negative values indicating greater activity earlier (adapting responses) and positive values indicating greater activity later (facilitating responses). Unless otherwise stated *RI* was calculated between the first and last 10s of stimulation.

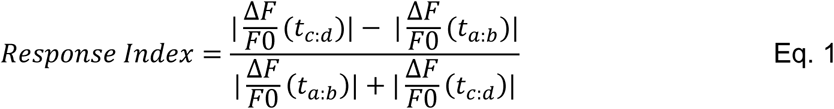

*RI* was used to classify neurons into different response classifications (e.g. adapting, non-adapting, facilitating). A gaussian mixture model of 3 components was fitted to all neuronal responses from animals expressing syn-GCaMP8m to predict the response classification (Fig. S7). We set two criteria to quantify whether a neuron switched its response class following astrocyte manipulation. First, neurons who did not maintain consistent response class throughout the trials were excluded. Second, candidate class switches were considered only when the change in *RI* was greater than two standard deviations of the respective class distribution, thus preventing false positive switches when the *RI* was close to the classification threshold.

The time above threshold was defined as the cumulative duration that an ROI remained above threshold from stimulation onset through 30s after stimulation. For mesoscale imaging, activation area was defined by the most distal active bin for which all intermediate bins were also active. To compare activity dynamics between cell types or experimental conditions, signals were averaged across a common radial range defined separately for each stimulation paradigm as the mean activated distance across animals plus one standard deviation (Fig. S5). In DREADD and photostimulation experiments, this range was defined from the pre-C21 and no-photostimulation conditions, respectively. Post-stimulation activity below baseline was classified as depressed when fluorescence fell >2SD below the baseline mean. Astrocyte calcium propagation speed was quantified from radial activity profiles under two-photon imaging by dividing the distance between the first 100 µm and the 200-300 µm regions by the difference in the peak activity time between these resions for trials showing above threshold activity in both regions.

## Statistical Analysis

All statistical analyses were performed in R(lme4 and emmeans packages^99^). Unless otherwise noted, differences between groups were assessed with linear mixed-effects models (LMMs) followed by type III F-tests, chosen for their robustness to unbalanced, complex designs and explicit accounting for within-animal variability. Models took the form analysis ∼ stimulation condition * manipulation + (1|animalid / trial / ROI), where manipulation denodes the DREADD or optogenetic stimulation group or ROI type (e.g. neuron vs astrocyte). Trial and ROI were included as nested random effects to incorporate repeated-measures and cell-level variability. Denominator degrees of freedom were approximated using the Satterthwaite method. Random effects that did not contribute to model variance (singular fits) were removed. Because mesoscale imaging did not include individual cellular ROIs, the ROI random effect was omitted for those analyses.

Three analyses differed from this primary model. To test how responses changes across repeated trials (Fig. 2, Fig. S6), trial was included as a mean-centered continuous predictor with stimulation condition and compartment, with a ROI random slope for trial. Second, to test whether the proportion of active ROIs changed across repeated trials, we used a binomial generalized linear mixed model on bounded counts (GLMM; logit link) of the form cbind(active,inactive) ∼ stimulation condition * manipulation + (1|animalid). This model was evaluated by Type III analysis of deviance. Third, to test associations between two measured signals (e.g. astrocyte vs. neuron mean activity, compression analysis) we model one signal as a function of the other with a random intercept for animal to account for repeated measures.

Following a significant factor or interaction effect (p<0.05), pairwise comparisons were performed on the estimated marginal means using the emmeans package. This approach incorporates the model’s covariance structure and applies the Holm correction for multiple comparisons. Data from male and female mice were pooled. Violin plots show the distribution of trial-averaged activity per ROI, where the overlayed box plot shows the median (horizontal line) and interquartile ranges. Because mesoscale data have no individual ROIs, Box plots show all trials from all animals, with the median highlighted by a horizontal line, as do the scatter plots for association analyses.

## Author Contributions: Conceptualization

K.S., T.D.K.Z., and A.V.; **Methodology:** K.S.; **Investigation:** K.S.; **Formal analysis:** K.S.; **Data curation:** K.S.; **Visualization:** K.S.; **Writing – original draft:** K.S. and T.D.K.Z.; **Writing – review & editing:** A.V.; **Supervision:** T.D.K.Z.; **Funding acquisition:** T.D.K.Z. All authors reviewed and approved the final manuscript.

## Supplementary Information

**Figure S1.**
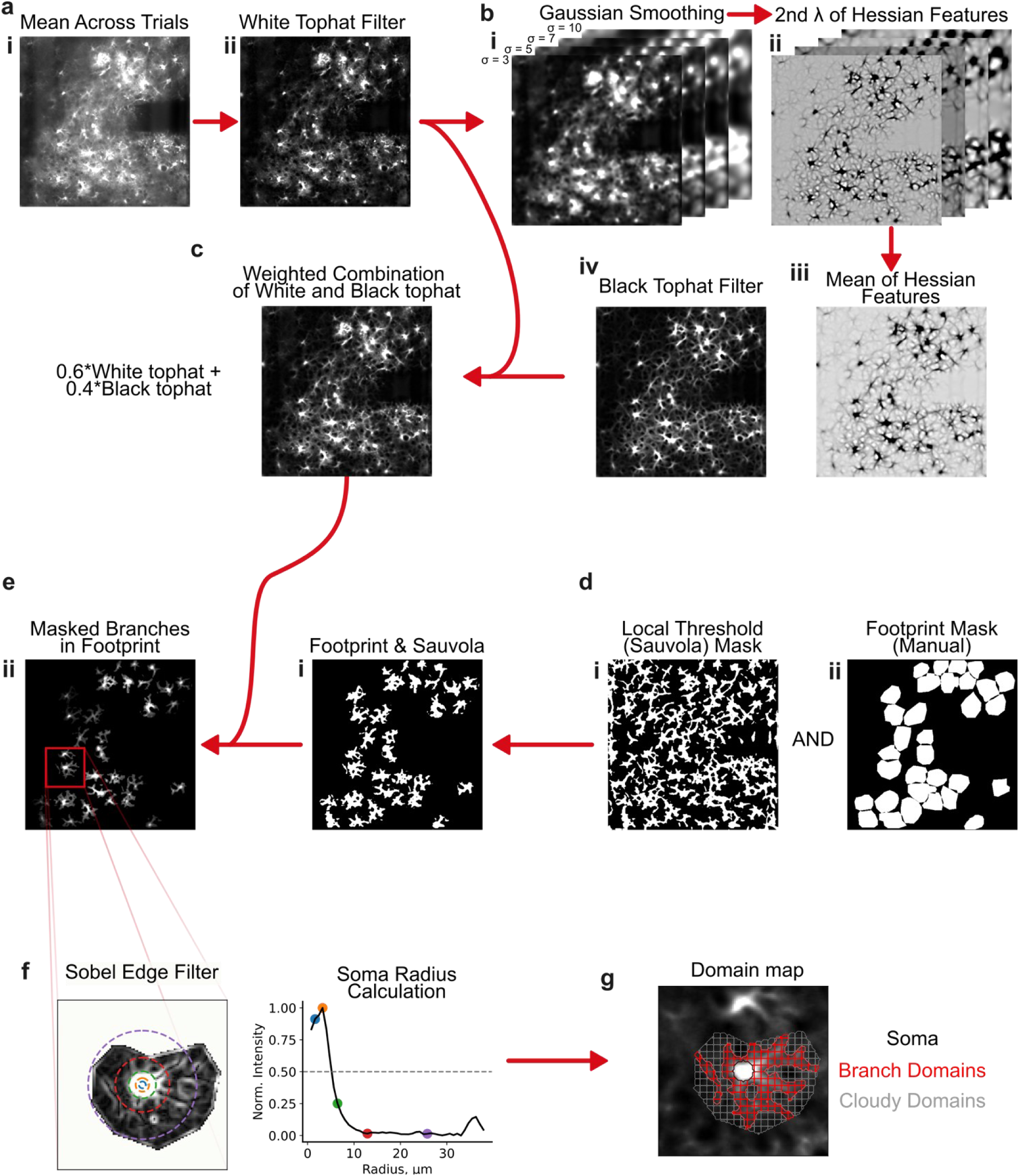
Preprocessing of images for semiautomatic segmentation of astrocyte soma from their branches and cloudy regions. **A**. A white top hat filter was applied to the trial averaged mean image for a given animal (ai) which helped to highlight astrocyte morphology from the background. **B**. The white tophat filtered image was smoothed with 4 different gaussian filters (i) and the second eigen value of the hessian feature matrix of those smoothed images (ii) were averaged together (iii). The sigmas for the gaussian smoothing were chosen to highlight astrocyte morphology without over-enhancing fine branching. A black tophat filter was used to highlight dark features of the hessian output. **C**. A weighted average of the hessian output and the white tophat filter was used to highlight original astrocyte morphology without overemphasizing hessian features. **D**. A Sauvola local threshold was used to identify bright regions representing astrocyte morphology. This threshold was logically combined with a region mask that was manually defined around identified astrocytes that contained clear soma and main branches (dii). **E.** The astrocyte footprint mask was merged with the weighted image. **F**. A Sobel edge filter was applied to a masked astrocyte region to highlight edges of the soma. The mean intensity of radial bins extending from the manually defined soma center was used to identify the edge of the soma, where the soma radius was defined as the distance where the mean intensity of a radial bin dropped below half of its maximal value. **G**. A circular ROI with radius defined in f, represented the soma, the thresholded region in (ei) excluding the soma was defined as the main branch region, and the remaining area in the manually defined astrocyte region was defined as the cloud region. The branches and cloud regions were then broken into domains of maximal size of ∼3×3 µm.

**Figure S2.**
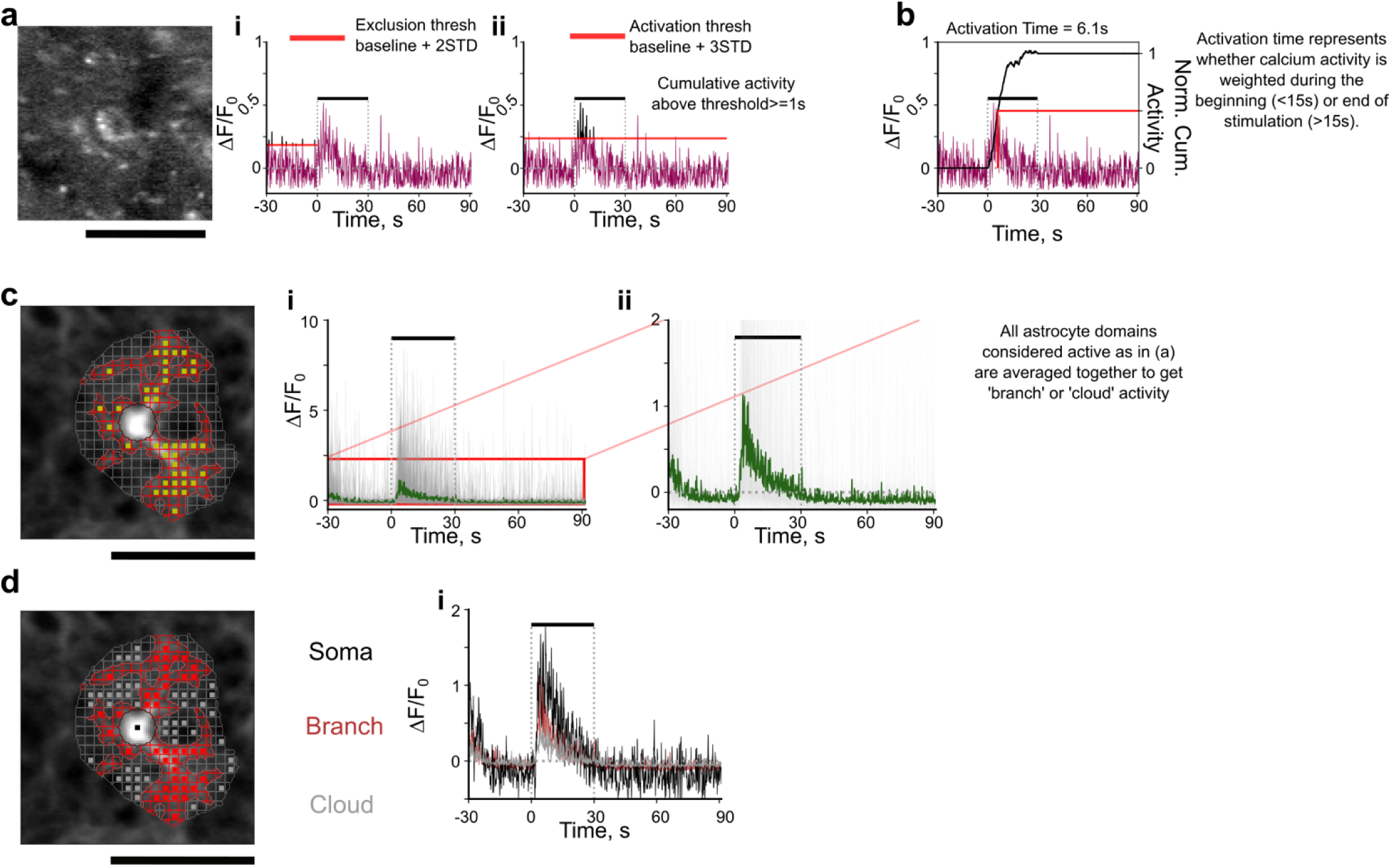
Demonstration of determination of activation for astrocytes or neurons. **A.** example neuron (left) and corresponding trace during stimulation for a given 100 Hz stimulation. To exclude certain potential activity from computation of noise floor baseline, activity above 2 standard deviations during the baseline period (threshold in red, activity above initial threshold represented in black) were excluded to recompute a more representative baseline and standard deviation of the noise level (i). The threshold was computed as the mean + 3 standard deviations above the recomputed baseline activity (red horizontal line). If neuronal or astrocyte calcium was elevated above this threshold for a total of 0.5, or 1s for neurons or astrocytes, respectively that neuron, or astrocyte domain was concluded to be activated for the given trial. Activity above threshold is represented in black. **B**. computation of activation time for the given neuron. Activation time represents the amount of time it takes to reach half the total cumulative calcium activity during stimulation. Black line (right axis) represents the normalized cumulative sum of activity during stimulation. Activation time is the time that the cumulative line reached half of total cumulative activity. **C**. Representative astrocyte and corresponding segmented domains (left). Yellow boxes represent branch domain regions determined to be activated during a given 100 Hz trial using method in (a). Their corresponding activity is represented by gray lines in (i). Because activity in these small domains were variable, they were averaged together to get activity for the branch domain (ii) showing clear elevation in calcium activity during stimulation. **D**. Representative image of the same astrocyte and corresponding activated soma, branch or cloud domains deemed activated (left), and their corresponding averaged activity for the 100 Hz stimulation trial.

**Figure S3.**
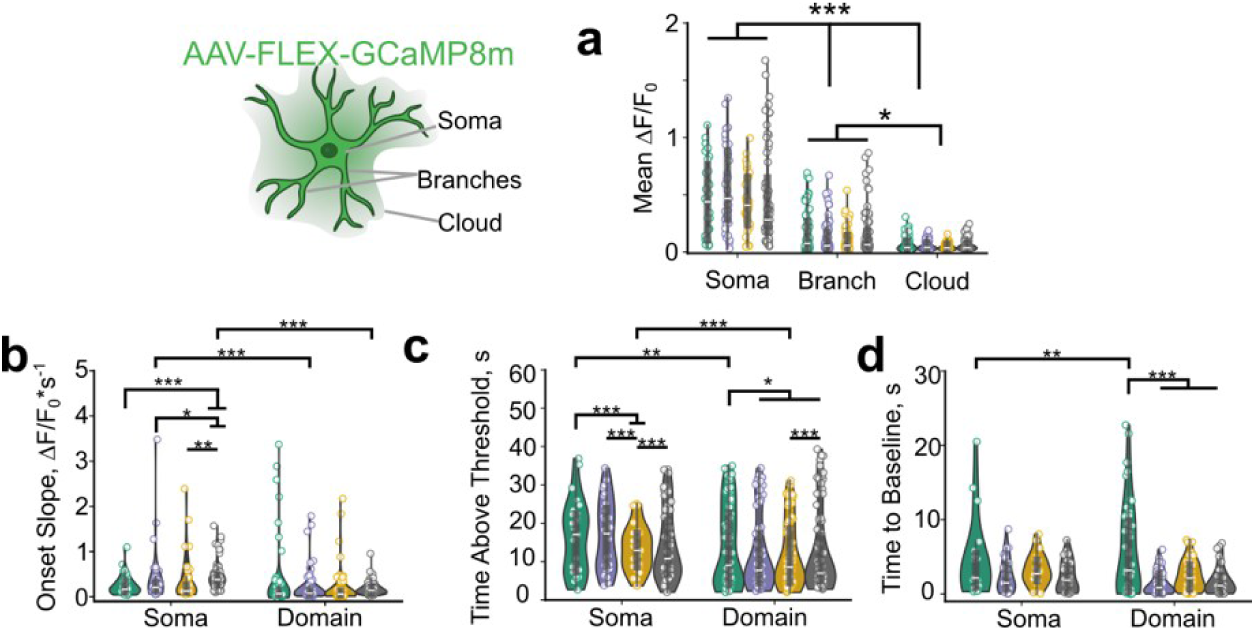
ICMS influences astrocyte calcium activity depending on stimulation pattern and compartment. **A)** (Left) Graphic representing different astrocyte compartments. ‘Domain’ represents the average of branch and cloud regions. (Right) Astrocyte soma display the significantly larger mean calcium activity in response to ICMS compared to branch and cloud domains (LMM; Stim: 2.8e-85, ROI type: p = 8.0e-16, interaction: p=8.7e-57). Activated branch and cloud domains were averaged together for future analyses. **B)** The activation slope of calcium activity in the somata is influenced by stimulation pattern (LMM; Stim: 3.8e-3, ROI type: p=2.0e-4, interaction: p=6.1e-3, trial: p=0.53). **C)** Stimulation parameters significantly influence the time of activity above threshold of astrocyte somata and domains (LMM; Stim: 3.6e-26, ROI type: p=0.06, interaction: p=5.7e-8, trial: p=0.04). **D)** Astrocytes display sustained calcium elevations after stimulation ends that is influenced by stimulation pattern in the domains (LMM; Stim: 4.1e-140, ROI type: p=9.5e-4, interaction: p=0.03, trial: p=0.22). N=3, n=31-82, 6 trials.

**Figure S4.**
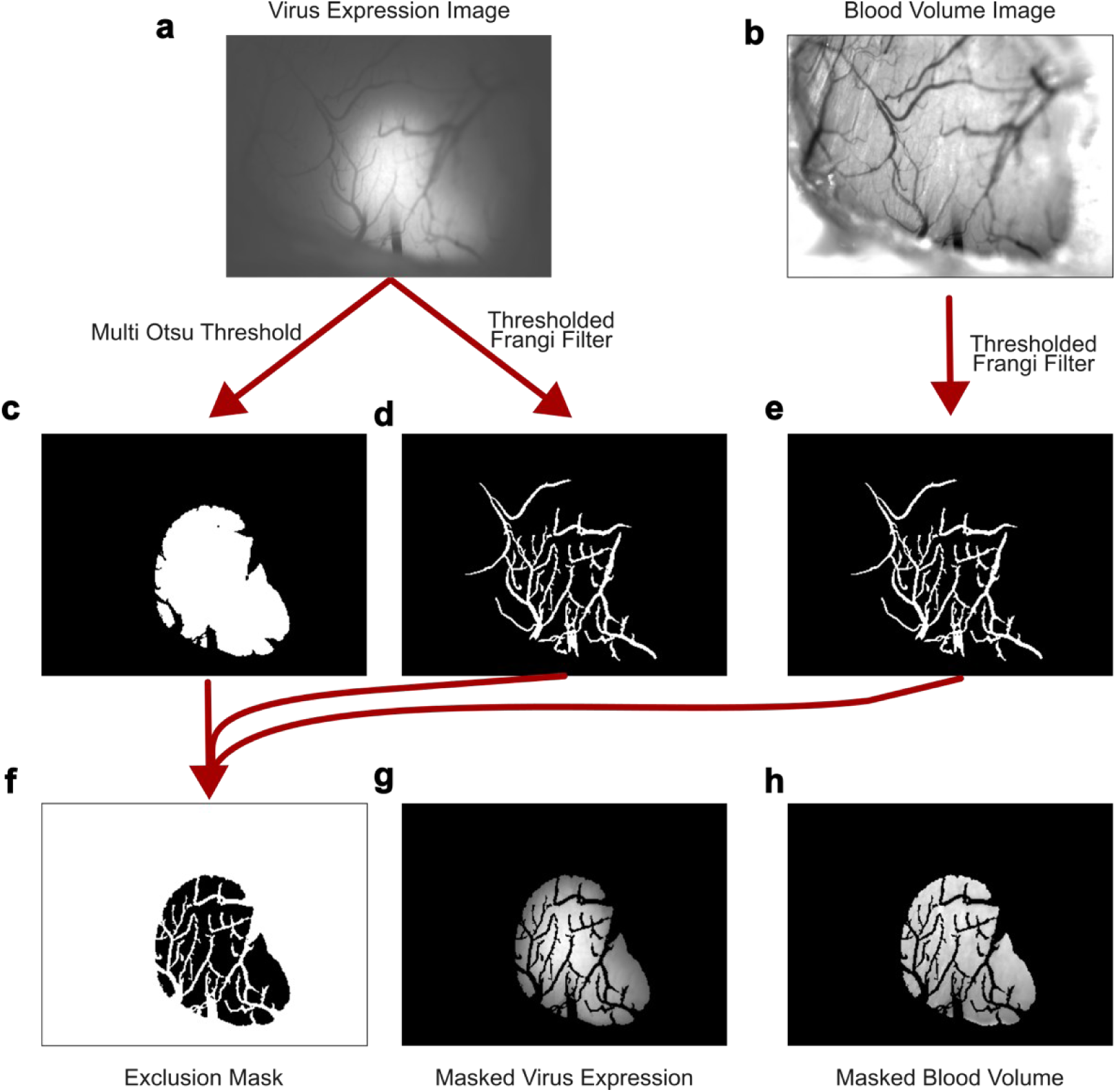
Method for mesoscale region and vessel masking. **A.** Representative average projection of virus expressing region. **B**. Average projection of raw green LED reflectance (for blood volume calculation). **C**. Area including virus expression was identified by selecting the connected pixels above the second threshold of a multi-Otsu threshold with 3 regions. **D-E**. Thresholding a Frangi filter applied to the virus expressing image and the blood volume image highlighted the large blood vessels on the brain surface. **F**. A mask to exclude non-expressing regions and blood vessels was created through a logical combinations of the masks in C-E. **G-H**. Representative virus expression and blood volume image masked by the exclusion mask.

**Figure S5.**
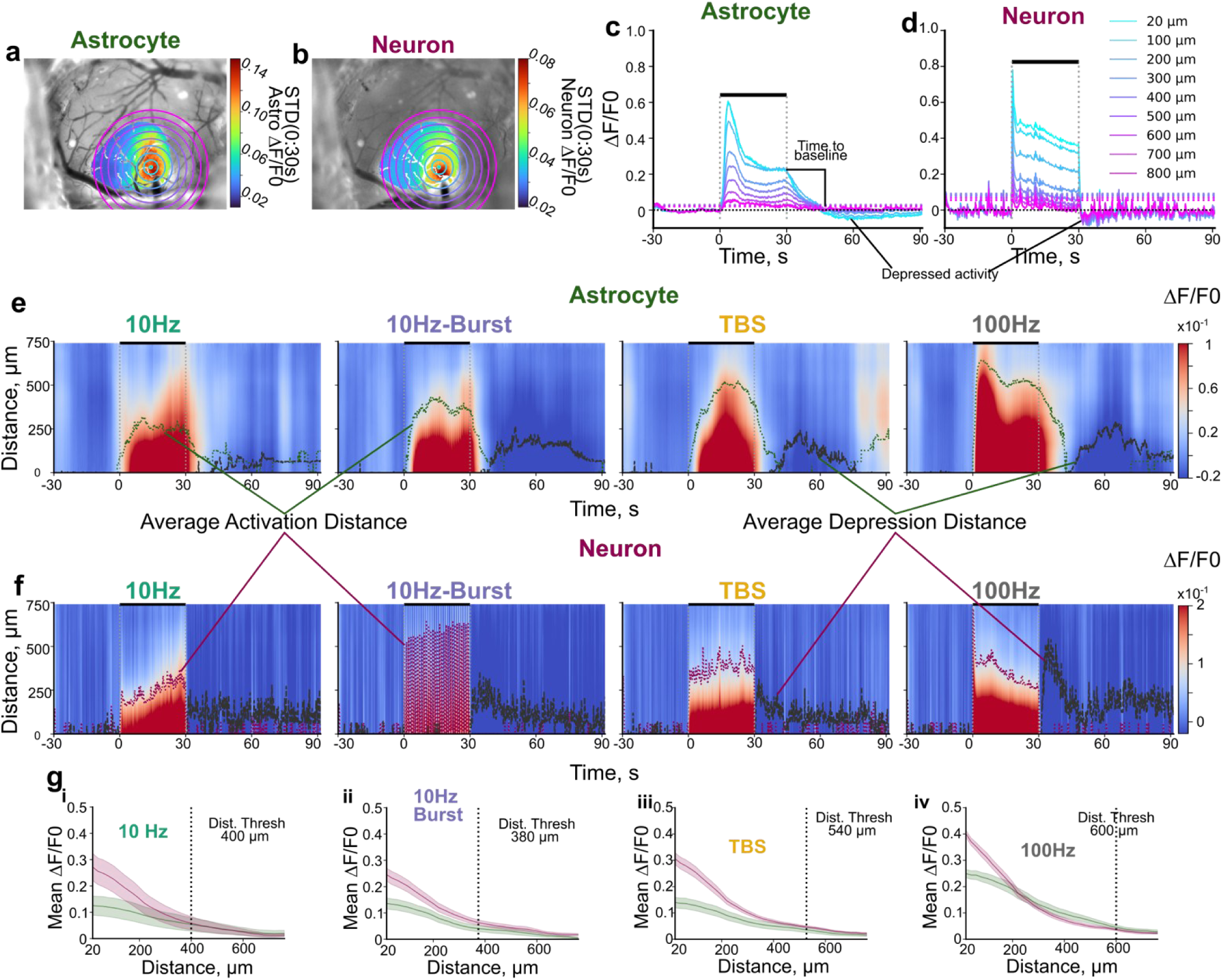
Identification of ‘activated’ regions in mesoscale imaging. A-b. Standard deviation projections within the masked regions for astrocyte (a) or neuronal (b) activation during a 100 Hz stimulation trial with radial bins drawn relative to the center of neuronal activation. C-d. Corresponding traces and activation threshold (dotted horizontal lines of corresponding color) for radial bins drawn in a-b for astrocytes (c) or neurons (d). Activity traces often depress below baseline (black horizontal line) after stimulation cessation. E-f Average heatmap projections across animals and trials over time and distance highlighting periods of high activity (red) during stimulation and depressed activity (blue) after stimulation ends for astrocytes (e) and neurons (f). Green and red traces highlight the average activation distance across animals over time for astrocytes and neurons. Dark gray traces highlight the average distance for depressed activity, which elevates after stimulation ends. g. Average ± standard error of activity over distance for different stimulation paradigms. The average distance to compare astrocyte and neuronal activity patterns represents the average activation area + 1 STD across animals and cell types for each stimulation condition.

**Figure S6.**
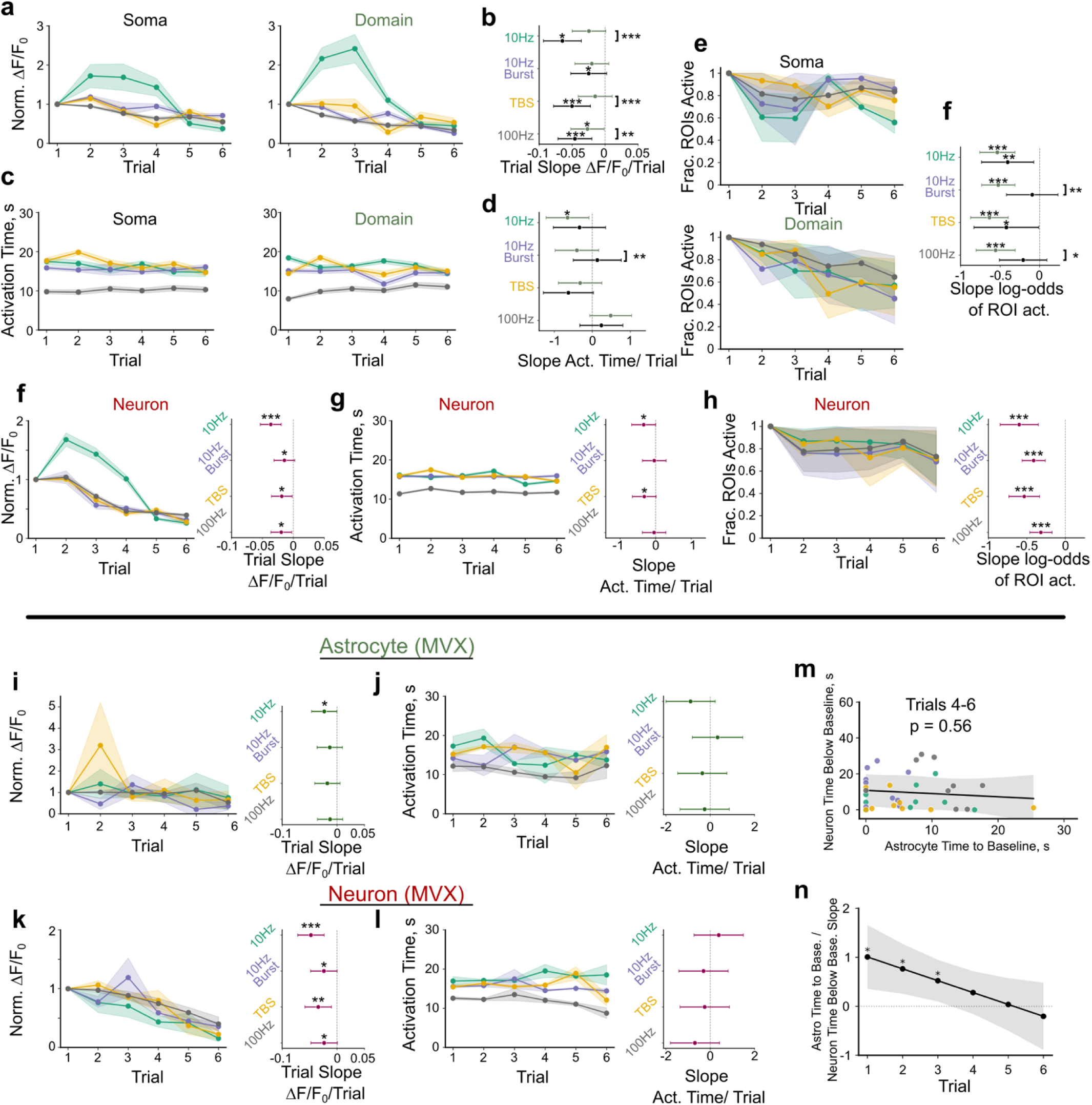
Magnitude of activation decays over trials, with temporal structure largely intact. A-b. Magnitude of astrocyte soma and domains reduces over trials (LMM Stim: p = 9.8e-87, compartment: p = 1.4e-9, trial: p = 5.6e-4, Stim x compartment: p = 1.9e-50, Stim x trial: p = 7.5e-3, domain x trial: p = 2.4e-10, N=3, n=31 - 82). Fitted slope ± 95% CI of magnitude of activation shows dependence on stimulation parameter and astrocyte compartment (b). C-d. Activation time of astrocyte compartments is largely consistent over trials LMM Stim: p = 4.1e-140, compartment: p = 9.5e-4, trial: p = 0.23, Stim x compartment: p = 0.03, Stim x trial: p = 1.0e-12, domain x trial: p = 0.73 N=3, n=31 - 82). D Fitted slope ± 95% CI of trial effect shows dependence on stimulation pattern and astrocyte compartment. E-f. Fraction of activated astrocyte compartment ROIs decays with trial. Fitted slope of the log-odds of an astrocyte ROI being active over trials shows decay with trial (Binomial GLMM: Stim: p= 1.7e-4, Compartment: p = 0.14, trial: p = 6.4e-16, stim x compartment: p=0.14, stim x trial: p = 0.35, compartment x trial: p = 1.1e-4). F. Magnitude of neuronal ROIs decays with trial (LMM: Stim 9.6e-137, Trial: p = 2.0e-3, Stim x trial1.6e-43; N = 3, n = 215-280). G. temporal structure of neuronal activity changes in some stimulation parameters (See Fig 2 LMM Stim: p = 1.5e-259, ROI type 0.61, trial p=0.06, Stim x roi type: p = 2.1e-22, Stim x trial: p= 6.0e-11, roi type x trial: p = 0.79). H. Fraction of active neuron ROIs decays with trial (GLMM Stim: p = 1.4e-3, Trial: p = 1.7e-9, Stim x trial: p = 0.037). *Right:* Slope of the log-odds of an ROI being active. I Magnitude of mesoscale astrocyte activity decays over trials (LMM Stim: p = 2.9e-3, Trial: p = 0.012, stim x trial: 0.76). j. Temporal structure of mesoscale astrocyte activity may change over trial (See Fig 2m stats: no significant posthoc decay with trial). K Magnitude of mesoscale neuronal activity reduces over trials (LMM Stim: p = 0.86, Trial p = 3.7e-5, Stim x trial p = 0.19). L. Temporal structure of neuronal activity is largely consistent over trials (see Fig 2m for stats, no significant posthoc decay with trial). M-N. The association of astrocyte time to baseline with neuron suppression depends on trial (See Fig 2o for stats). The relationship is not present for last three trials (M).

**Figure S7.**
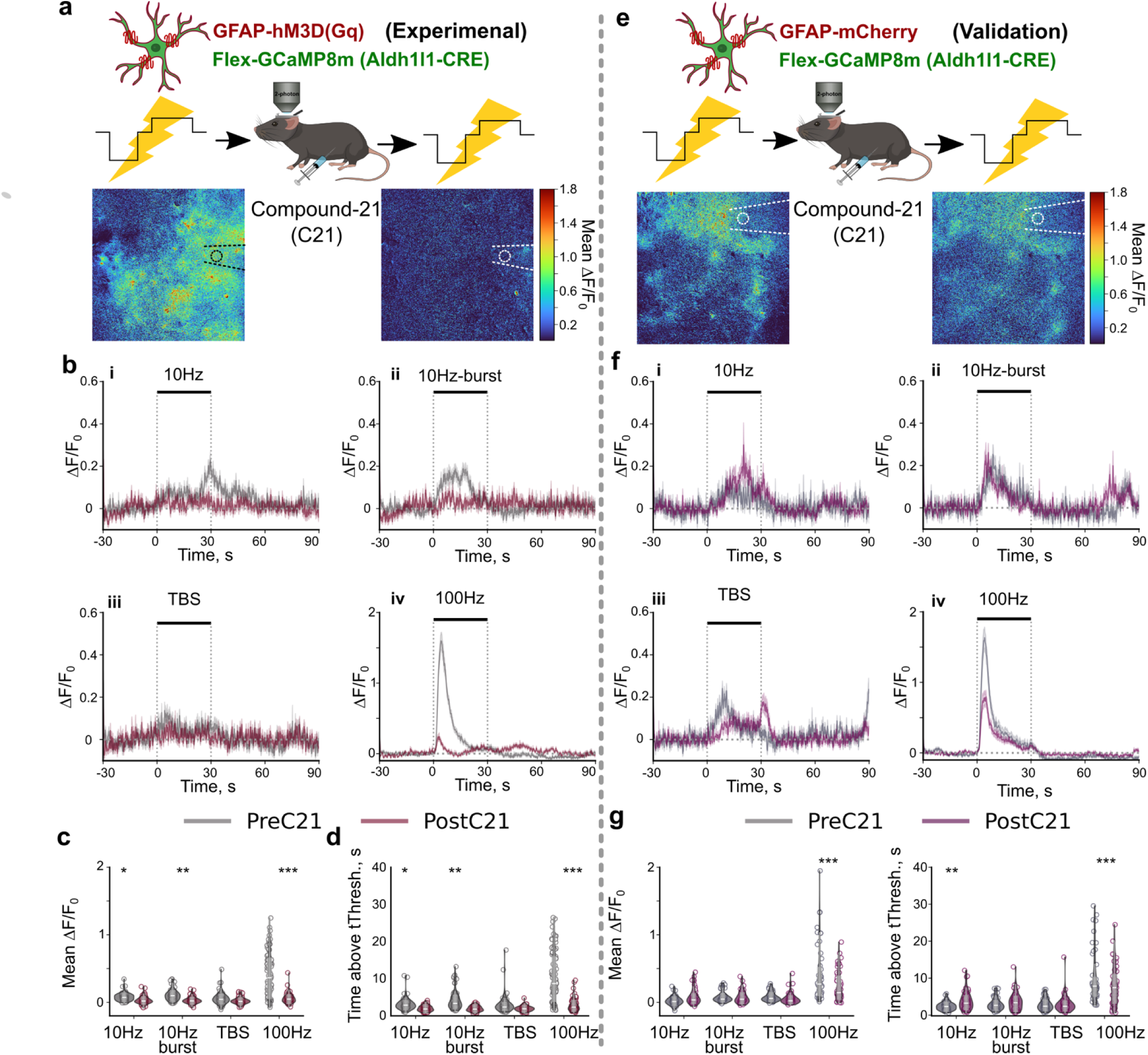
Gq DREADD activation in astrocytes reduces stimulation induced calcium activity under two-photon imaging. **A**. Aldh1l1-mice expressing GCaMP8m and hM3Dq in astrocytes received intracortical microstimulation before and after administration of 1 mg/kg of the DREADD agonist C21. Images show the reduced average magnitude of stimulation induced calcium activity during 100 Hz stimulation following administration of C21. **B**. Trial averaged traces ± standard error across cells across stimulation paradigms demonstrate that stimulation induced elevations in astrocyte calcium activity are substantially reduced following administration of C21. **C-D**. Mean stimulation induced astrocyte calcium activity (C, LMM; Stim: p=2.8e-108, C21: p= 9.2e-6, interaction: p=1.1e-79) and time above threshold (D, LMM; Stim: p=1.4e-113, C21: p= 1.2e-5, interaction: p= 1.6e-75) were significantly reduced following administration of C21. E-G similar analyses in validation animals expressing mCherry in place of hM3Dq did not display similar magnitudes of reduced stimulation-induced astrocyte calcium activity following administration of C21. **E-G.** Validation animals lacking hM3D. **E**. Aldh1l1-mice expressing GCaMP8m and mCherry in astrocytes received intracortical microstimulation before and after administration of 1 mg/kg of the DREADD agonist C21. Images show relatively comparable average magnitude of stimulation induced calcium activity during 100 Hz stimulation following administration of C21. **F**. Trial averaged traces of astrocyte calcium activity ± standard error across cells demonstrate that C21 administration without DREADD expression do not eliminate stimulation induced astrocyte calcium activity. **G-H**. Calcium activity was significantly reduced after C21 administration only during 100Hz stimulation (G; LMM; Stim: p=1.3e-82, C21: p= 4.4e-3, interaction: p= 3.4e-13), but time above threshold was either increased or decreased depending on stimulation pattern (H; LMM; Stim: p=1.5e-102, C21: p= 0.97, interaction: p=4.8e-9). N=2 animals across 3 trials, n = 48-50. *:p<0.05, **:p<0.01, ***:p<0.001.

**Figure S8.**
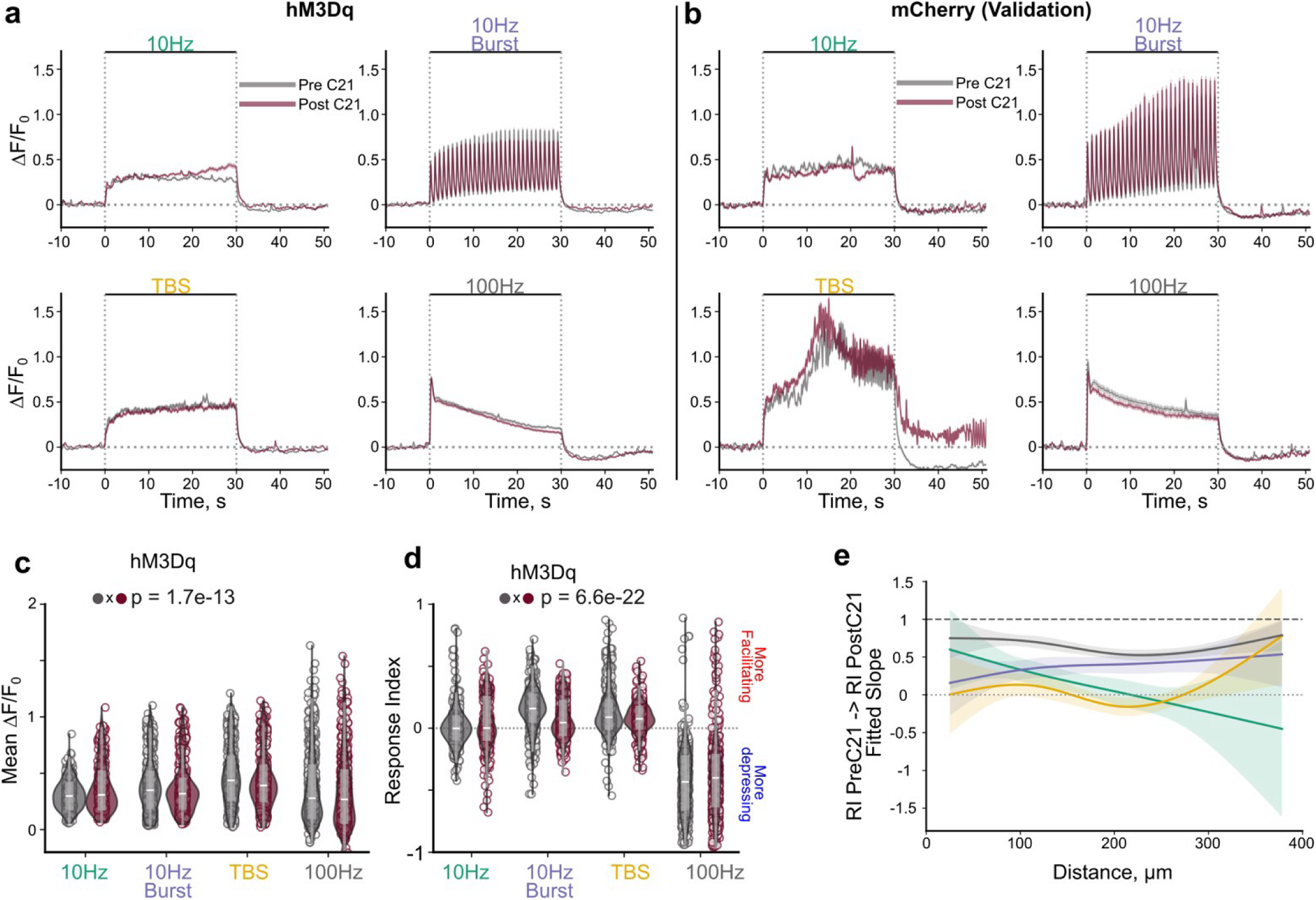
Astrocyte calcium attenuation does not cause global shifts in stimulation induced neural activity. **A.** Traces of neural activity before and after astrocyte attenuation. **B.** Traces of stimulation-induced activity in animals lacking hM3D in astrocytes show no major effects of C21 administration itself. **C**. Mean stimulation-induced activity before and after astrocyte attenuation (LMM: Stim: p<2.2e-16, C21: p=0.97, interaction: p=1.7e-13 N=4, n=403-713). **D.** Response index of neurons before and after attenuating astrocytes (LMM; Stim: p<2.2e-16, C21: p=0.99, interaction: p=6.6e-22, N=4, n= 403-713). **E**. Visualization of the relationship between the response index PreC21 to PostC21 of the form RI PostC21 ∼ RI PreC21 * distance * stimcondition + (1|animalid). Plotting was computed by fitting a natural cubic spline to the centered distance to allow visualization of non-linear variance over distance. Shading is 95% confidence interval (LMM; PreC21 p=3.8e-11; distance: p=2.5e-3; Stim: p=8.5e-3; PreC21 x Distance: p= 1.8e-5, PreC21 x Stim: p = 0.05, Distance x Stim: 1.3e-3; Distance x PreC21 x Stim: p= 1.7e-12; N=4, n= 344-674).

**Figure S9.**
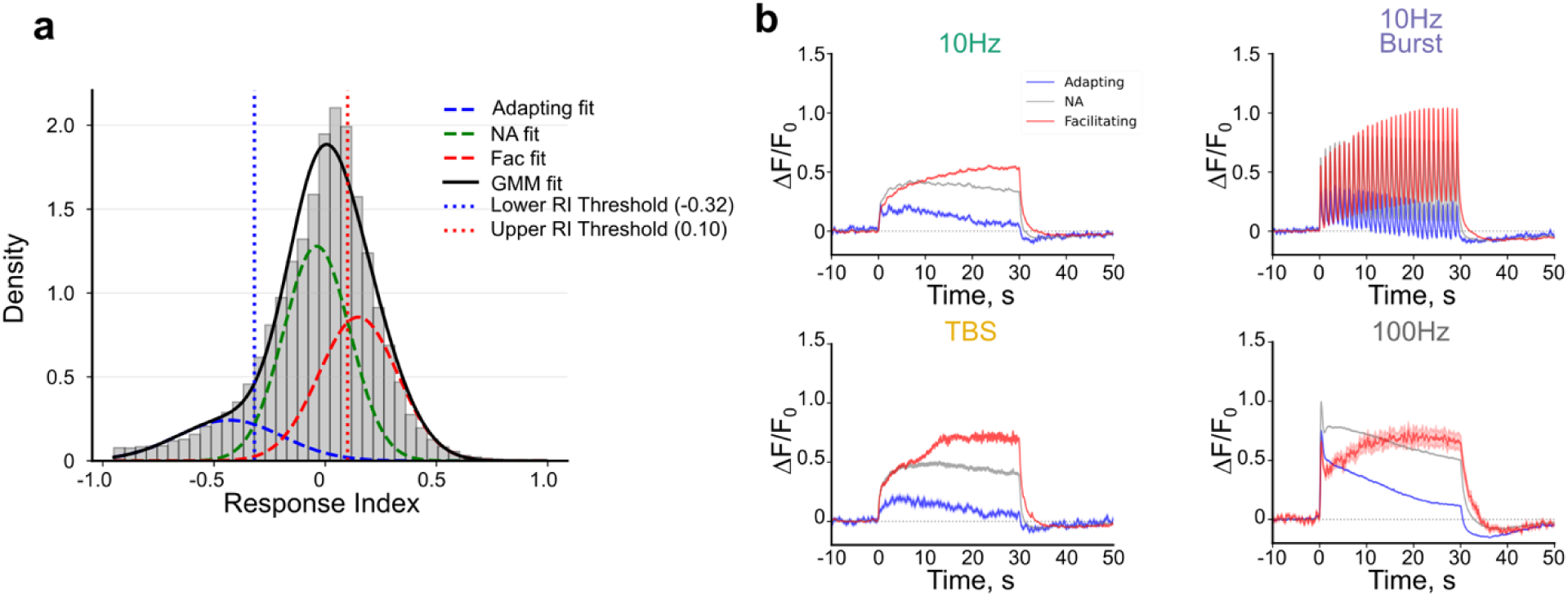
Neuron responses to ICMS display three general profiles facilitating, non-adapting, and adapting. A. A gaussian mixture model (GMM) of 3 distributions was fit to the response index of all neuron responses with GCaMP8m from this study (43,422 total responses across 1799 neurons from 12 animals). Data includes responses with hM3D and melanopsin manipulations and validation animals. B. Neuron responses (mean ± stderr shade) to each stimulation condition grouped by the respective GMM threshold highlights different classes of responses (facilitation: increase over time; non-adapting: small change over time, adapting: decrease over time).

**Figure S10.**
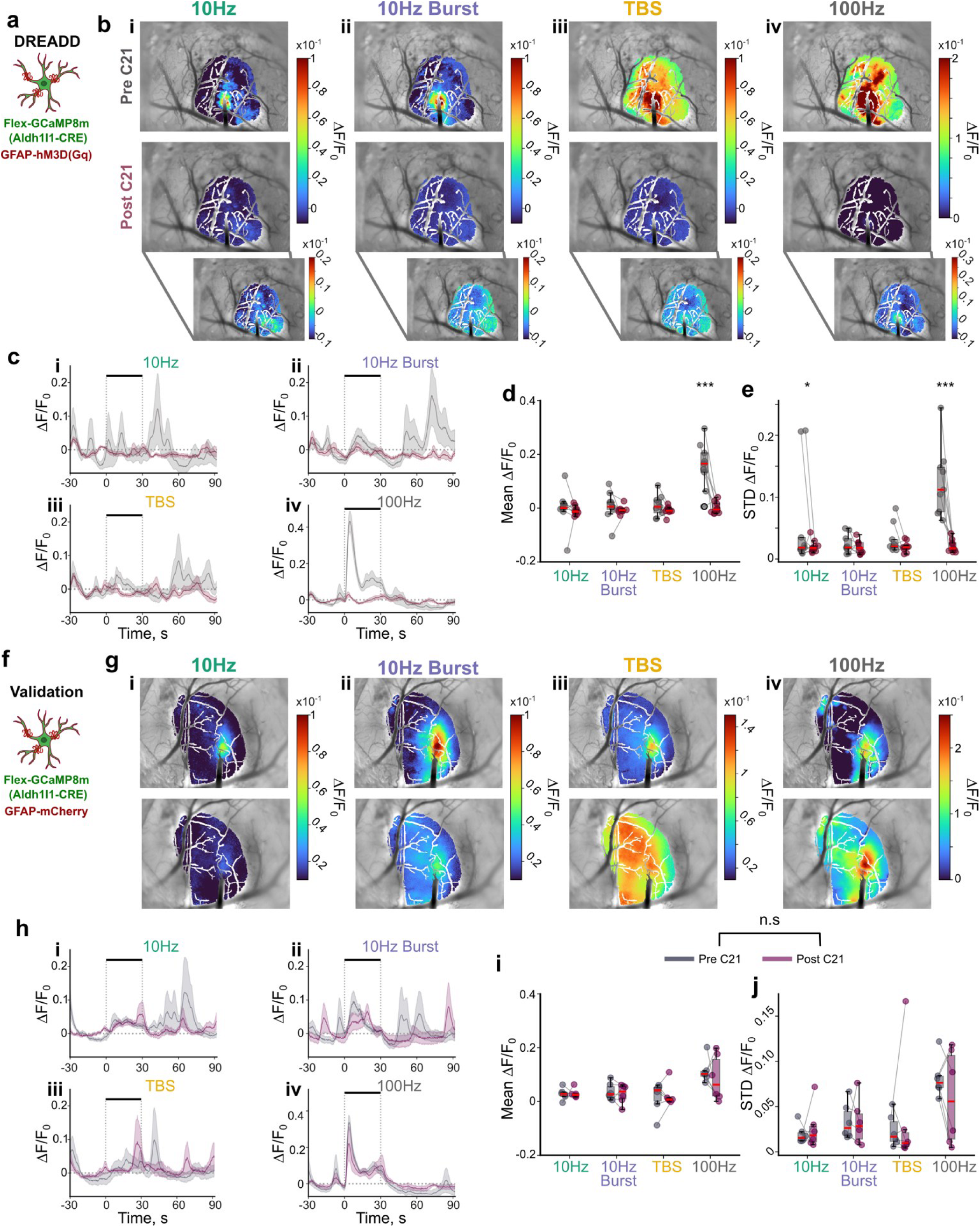
GqDREADD activation in astrocytes by 1 mg/kg C21 attenuates stimulation induced mesoscale astrocyte calcium activity. **A.** An Alhd1l1-CRE animal was injected with FLEX-GCaMP8m and GFAP-hM3DGq-mCherry >4weeks before implantation of an electrode to express GCaMP and a Gq DREADD in astrocytes. **B**. Representative images highlighting the mean stimulation induced astrocyte calcium activity within the virus expressing region is virtually non-existent for all stimulation patterns, except 100 Hz. **C**. Trial averaged mesoscale astrocyte calcium activity ± standard error across animals is reduced following astrocyte DREADD activation (Pre-C21 in Gray, Post-C21 in red). (**D-E**) Mean (D; LMM; Stim: p=1.25e-6, C21: p=4.19e-5, interaction: p=1.51e-6) and standard deviation of calcium activity (E; LMM; Stim: p=4.8e-4, C21: p= 2.81e-5, interaction: p= 8.2e-4) during stimulation is significantly reduced following DREADD activation. **F**. Validation animals were created in which Aldh1l1-CRE animals express GCaMP8m in astrocytes and mCherry (GFAP-mCherry) in place of the Gq DREADD. **G**. Representative images of mean stimulation-induced astrocyte calcium activity within the virus expressing region are slightly reduced in some stimulation trials. **H**. Mean trial averaged mesoscale astrocyte calcium activity ± standard error across animals appears relatively unchanged following administration of C21. I-j) mean (I; LMM; Stim: p=2.4e-5, C21: p= 0.54, interaction: p=0.71) and standard deviation (j; LMM; Stim: p=2.7e-3, C21: p=0.94, interaction: p=0.55) of mesoscale astrocyte calcium activity is not significantly attenuated following administration of C21. N=2-3 animals across 3 trials *:p<0.05, **:p<0.01, ***:p<0.001.

**Figure S11.**
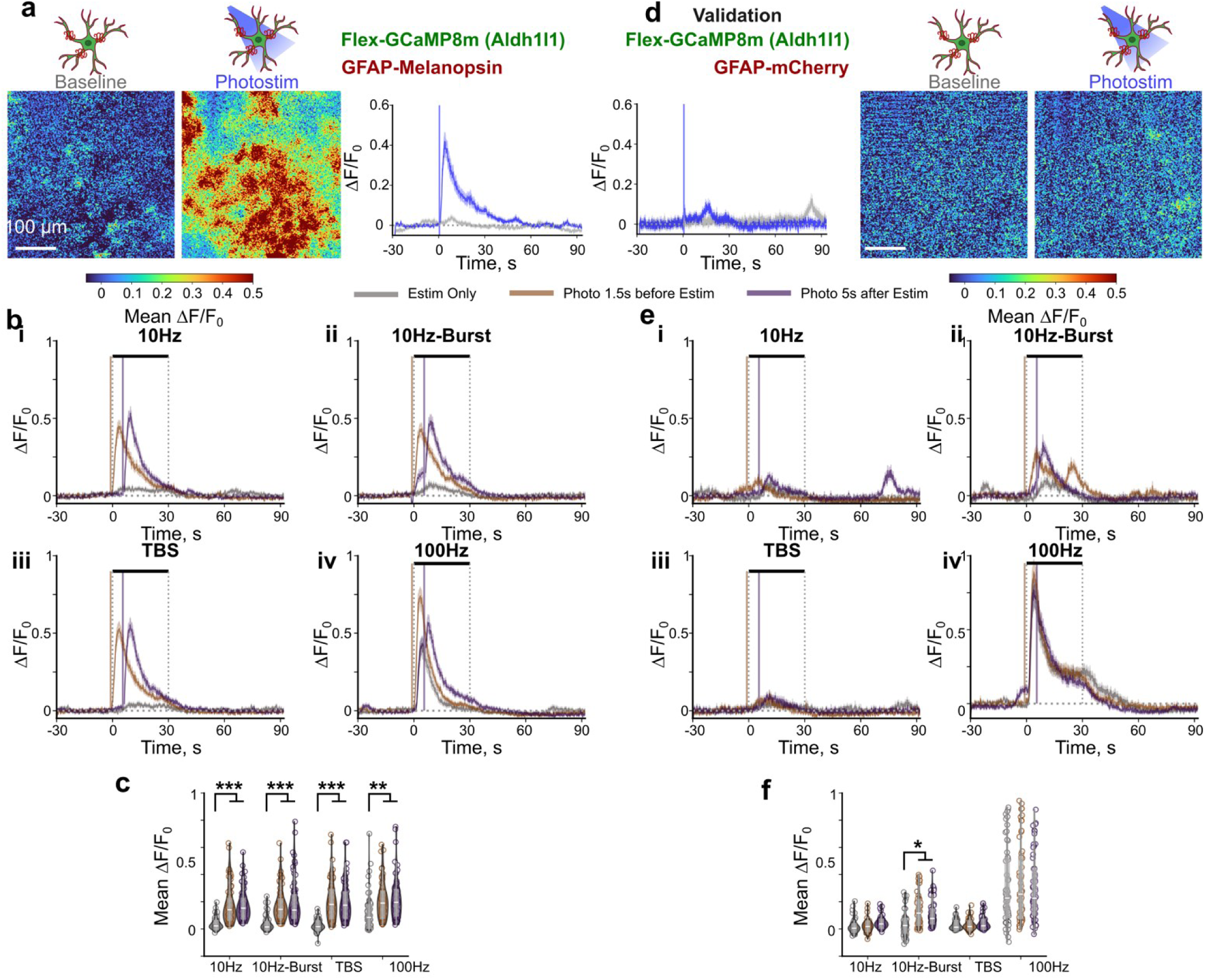
Photostimulation of melanopsin expressing astrocytes significantly increases astrocyte calcium activity during ICMS. **A**. Representative mean images demonstrating that photostimulation increases calcium activity within astrocytes (left) in mice expressing melanopsin and GCaMP8m in astrocytes. The mean calcium activity of all identified astrocytes was elevated following 1s of 1mW photostimulation at 10 Hz with a 30ms pulse width. **B**. Average traces of astrocyte calcium activity during different ICMS patterns showing that photostimulation elevates calcium activity whether photostimulation is delivered 1.5s before (orange traces) or 5s after ICMS (purple traces) compared to ICMS alone (gray traces). Notably, despite high levels of astrocyte calcium activity during 100 Hz stimulation alone, photostimulation still could amplify it suggesting it is not being saturated. **C**. the mean stimulation induced calcium activity is significantly elevated with both photostimulation conditions (LMM; Stim: p=1.7e-34, photostim: p=4.6e-5, interaction: p=2.4e-16) N=3, n=65. **D-F**. similar analyses in validation animals show that photostimulation in animals lacking the photosensitive opsin does not amplify astrocyte calcium activity (LMM; Stim: p=1.43e-215, photostim: p= 0.34, interaction: p=2.5e-6, N=2, n=58)*:p<0.05, **:p<0.01, ***:p<0.001.

**Figure S12.**
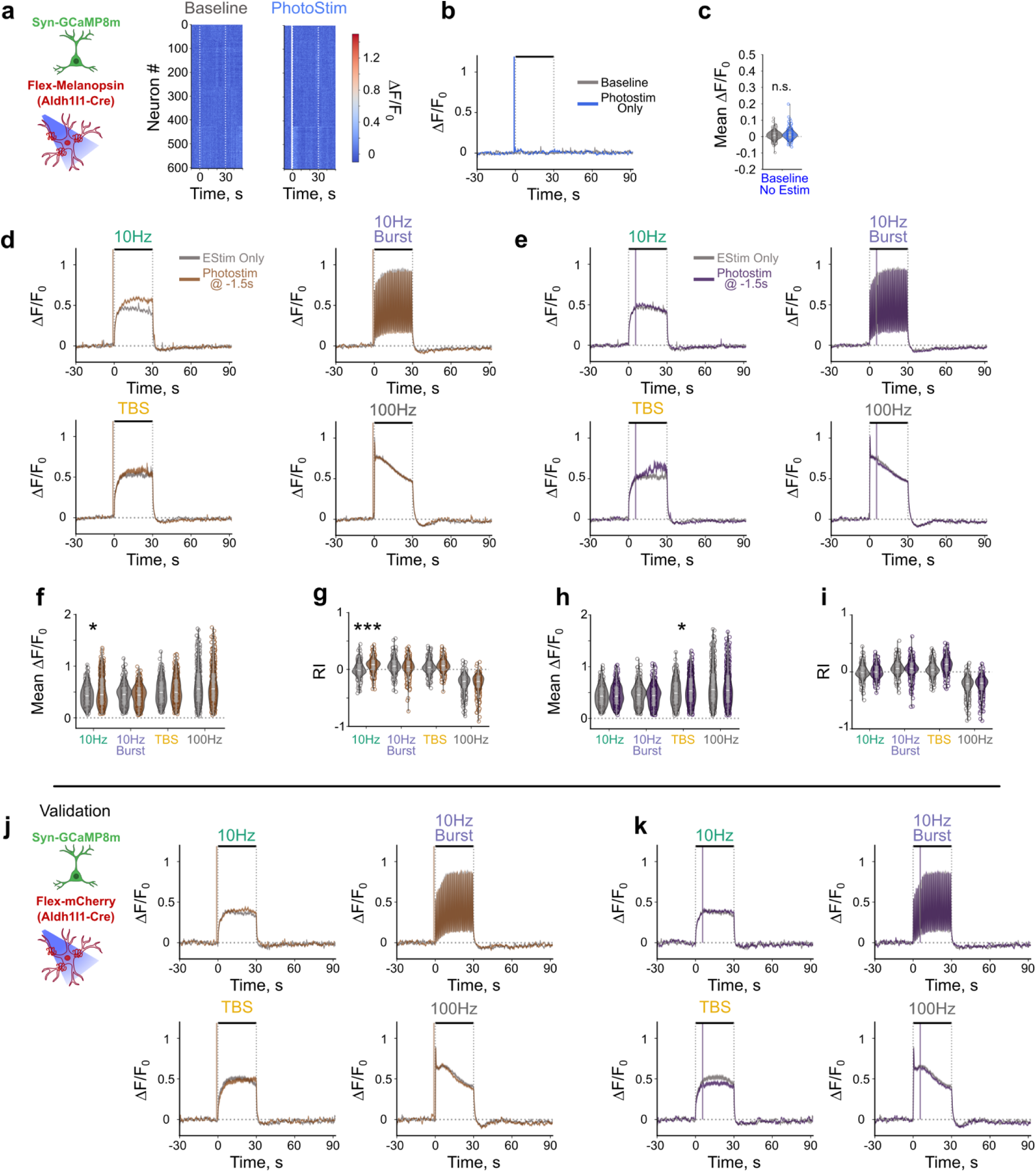
Photostimulation of melanopsin expressing astrocytes does not globally affect neuronal responses to ICMS. **A-b**. Heatmap of trial averaged neuronal responses (a) and traces of neuronal responses (b) shows that photostimulation of melanopsin in astrocytes does not increase neuronal calcium activity. C. Mean neuronal activity within 30 s of astrocyte photostimulation is comparable to baseline with no photostimulation. d-e. Traces of neuronal activity elicited by ICMS alone (gray) and astrocyte photostimulation pre-ICMS (d, orange) or post-ICMS (e, purple). F. Mean calcium activity of neurons during ICMS is increased with photostimulation of astrocytes pre-ICMS during 10 Hz stimulation (LMM Stim p<2e-16, Photostim: p = 0.76; Estim x Photo: p = 5.9e-24). G. Response index of neuronal activity during ICMS is increased with astrocyte photostimulation pre-ICMS during 10 Hz stimulation (LMM: Estim p <2e-16, Photostim: p = 0.46, interaction p = 1.3e-37). H. Mean neuronal activity of neuronal calcium activity during ICMS is increased with astrocyte photostimulation post-ICMS during TBS stimulation (LMM Estim: p <2e-16, Photostim: p = 0.92, interaction: p=3.6e-18). I Response index of neuronal activity is increased with photostimulation of astrocytes post-ICMS during TBS stimulation (LMM EStim: p<2e-16, Photostim: p = 0.31, interaction: p = 8.4e-38). j-k. Traces of neuronal calcium activity with photostimulation of animals lacking melanopsin pre-ICMS (j) or post-ICMS (k) does not show substantial change in neuronal response to ICMS. N=2-3 animals across 3 trials. *:p<0.05, **:p<0.01, ***:p<0.001.

**Figure S13.**
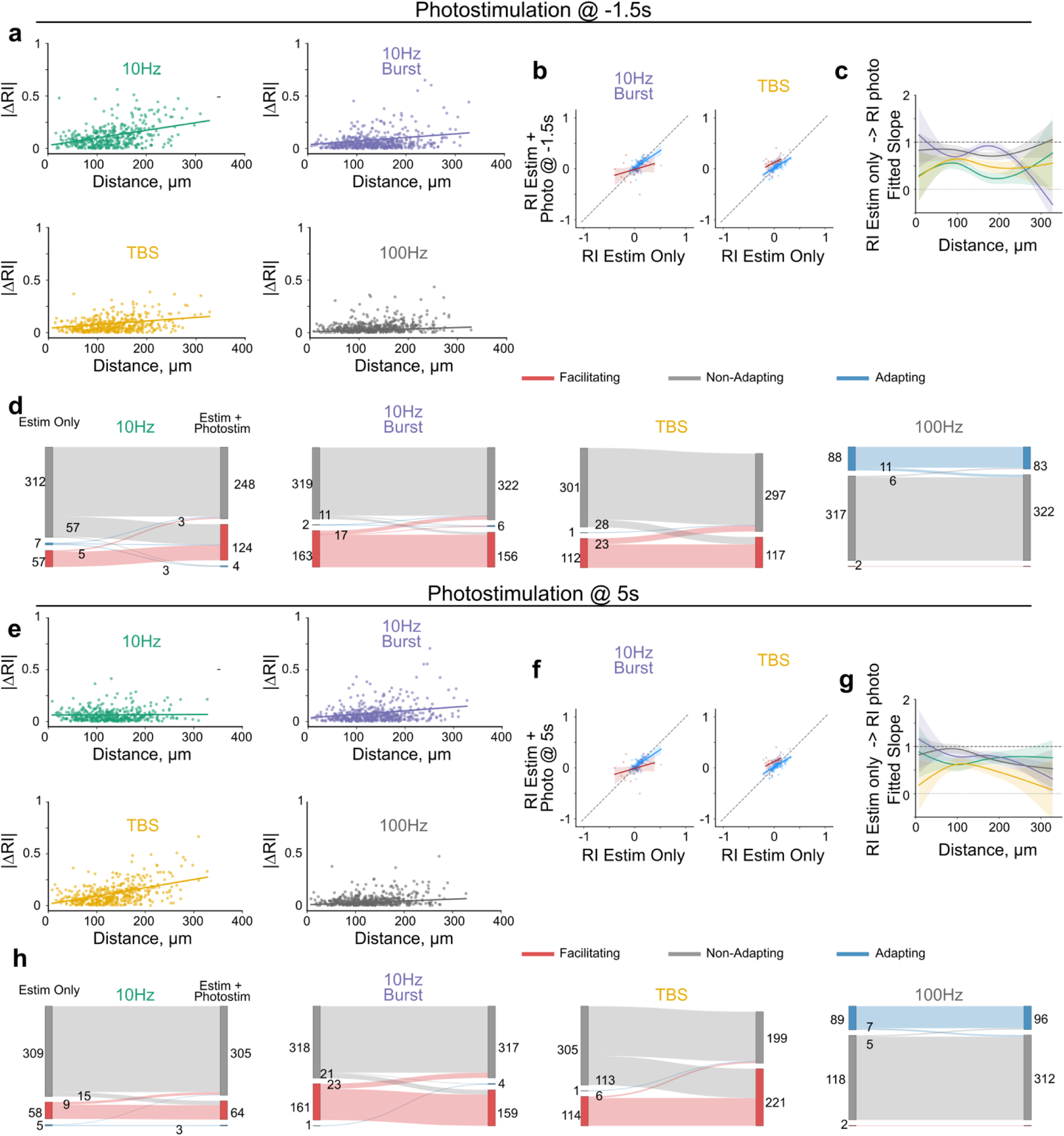
Photostimulation of astrocytes elicits heterogeneous effects on temporal profile of ICMS-induced neuronal activity. A. Magnitude of change of the response index with astrocyte photostimulation pre-ICMS increases with distance from the electrode depending on stimulation parameter. Scatter plots of fitted slopes in Fig. 5i (see Fig. 5i for stats). B-C. Fitted slope over distance of RI of neurons with photostimulation of astrocytes pre-ICMS over RI of neurons with ICMS only (see Fig. 5k-l for stats). D. Small populations of neurons change their response profile with photostimulation of astrocytes pre-ICMS. E. Magnitude of change of the response index with astrocyte photostimulation post-ICMS increases with distance from the electrode depending on stimulation parameter. Scatter plots of fitted slopes in Fig. 5j (See Fig. 5j for stats). f. Fitted slope over distance of RI of neurons with photostimulation of astrocytes post-ICMS over RI of neurons with ICMS only (See Fig. 5m-n for stats). f. Small populations of neurons change their response profile with photostimulation of astrocytes post-ICMS.

**Figure S14.**
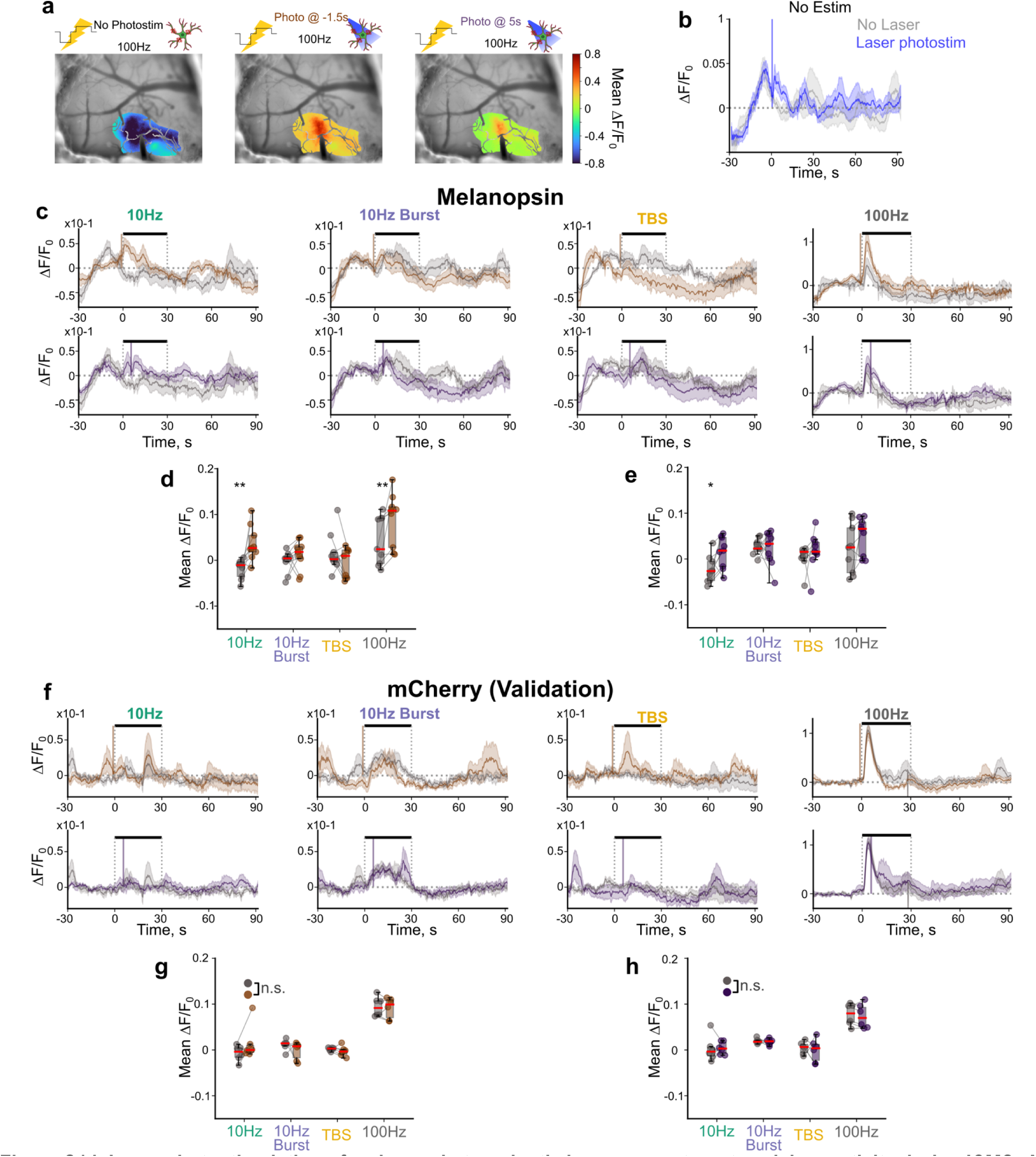
Laser photostimulation of melanopsin transiently increases astrocyte calcium activity during ICMS. **A**. Representative mean projection images during 100 Hz stimulation shows that astrocyte calcium activity is higher with laser photostimulation than without. **B**. The imaging modality is sufficient to increase astrocyte calcium activity during the baseline period before laser photostimulation. Laser photostimulation can still increase astrocyte calcium activity (blue trace) compared to without (gray) **C**. Laser photostimulation 1.5s before (orange) or 5s after (purple) transiently amplifies astrocyte calcium activity during stimulation more than electrical stimulation alone (gray). **D-E**. Astrocyte calcium activity is significantly higher 5s after photostimulation compared to no stimulation (gray) whether laser photostimulation is delivered 1.5s before electrical stimulation onset (d; LMM; Stim: p=9.9e-7, photostim: p= 4.9e-3, interaction: p=3.9e-3) or 5s after electrical stimulation onset (e; LMM; Stim: p=3.9e-3, photostim: p=0.019, interaction: p=0.25). **F**. Astrocyte calcium activity is comparable between trials without photostimulation (gray) and with laser photostimulation 1.5s before (orange) or 5s after (purple) electrical stimulation onset in validation animals. **G-H**. Photostimulation does not significantly affect astrocyte calcium activity in the 5s after photostimulation delivery in validation animals (g; LMM; Stim: p=1.74e-14, photostim: p=0.22, interaction: p=0.97; H; LMM; Stim: p=1.7e-11, photostim: p= 0.68, interaction: p=0.97). N=2-3 animals across 3 trials. *:p<0.05, **:p<0.01, ***:p<0.001.

## References

1 Treue, S. & Maunsell, J. H. Attentional modulation of visual motion processing in cortical areas MT and MST. Nature 382, 539–541 (1996).

2 Byrd, D. L., Marks, W. J., Jr. & Starr, P. A. Deep brain stimulation for advanced Parkinson’s disease. AORN journal 72, 387–390, 393-408; quiz 409-314, 416-388 (2000).

3 Flesher, S. N., et al. A brain-computer interface that evokes tactile sensations improves robotic arm control. Science 372, 831–836 (2021). 10.1126/science.abd0380

4 Hughes, C. L., et al. Perception of microstimulation frequency in human somatosensory cortex. eLife 10, e65128 (2021). 10.7554/eLife.65128

5 Penfield, W. & Welch, K. Instability of response to stimulation of the sensorimotor cortex of man. The Journal of physiology 109, 358–365, illust (1949).

6 Fritsch, G. & Hitzig, E. On the electrical excitability of the cerebrum. Some papers on the cerebral cortex, 73–96 (1870).

7 Romo, R., Hernandez, A., Zainos, A. & Salinas, E. Somatosensory discrimination based on cortical microstimulation. Nature 392, 387–390 (1998). 10.1038/32891

8 Agnesi, F., Johnson, M. D. & Vitek, J. L. Deep brain stimulation: how does it work? Handb Clin Neurol 116, 39–54 (2013). 10.1016/b978-0-444-53497-2.00004-8

9 Spix, T. A., et al. Population-specific neuromodulation prolongs therapeutic benefits of deep brain stimulation. Science 374, 201–206 (2021). doi:10.1126/science.abi7852

10 Kozai, T. D. Y., et al. Solving the Problem of Inception: A cross-species perspective on strategies for a mechanistic refinement of intracortical microstimulation. Journal of Neural Engineering 23, 023001 (2026).

11 Wood, K. C., Blackwell, J. M. & Geffen, M. N. Cortical inhibitory interneurons control sensory processing. Current opinion in neurobiology 46, 200–207 (2017).

12 Perea, G., Yang, A., Boyden, E. S. & Sur, M. Optogenetic astrocyte activation modulates response selectivity of visual cortex neurons in vivo. Nature communications 5, 3262 (2014). 10.1038/ncomms4262

13 Ahmadpour, N., et al. Cortical astrocyte N-methyl-D-aspartate receptors influence whisker barrel activity and sensory discrimination in mice. Nature communications 15, 1571 (2024). 10.1038/s41467-024-45989-3

14 Schmidt, E. M., et al. Feasibility of a visual prosthesis for the blind based on intracortical microstimulation of the visual cortex. Brain : a journal of neurology 119 (Pt 2), 507–522 (1996).

15 Hughes, C. L., Flesher, S. N. & Gaunt, R. A. Effects of stimulus pulse rate on somatosensory adaptation in the human cortex. Brain stimulation 15, 987–995 (2022).

16 Michelson, N. J., Eles, J. R., Vazquez, A. L., Ludwig, K. A. & Kozai, T. D. Y. Calcium activation of cortical neurons by continuous electrical stimulation: Frequency dependence, temporal fidelity, and activation density. Journal of neuroscience research 97, 620–638 (2019). 10.1002/jnr.24370

17 Hughes, C. & Kozai, T. Dynamic amplitude modulation of microstimulation evokes biomimetic onset and offset transients and reduces depression of evoked calcium responses in sensory cortices. Brain Stimulation 16, 939–965 (2023).

18 Hobbs, T. G., et al. Biomimetic stimulation patterns drive natural artificial touch percepts using intracortical microstimulation in humans. Journal of Neural Engineering 22, 036014 (2025).

19 Hughes, C. L., Chen, X., Grill, W. M. & Kozai, T. D. Y. Neural mechanisms underlying intracortical microstimulation for sensory restoration. Nature BME 10, 197–213 (2026).

20 Sanzeni, A., et al. Inhibition stabilization is a widespread property of cortical networks. eLife 9, e54875 (2020). 10.7554/eLife.54875

21 Gottlieb, J. P. & Keller, A. Intrinsic circuitry and physiological properties of pyramidal neurons in rat barrel cortex. Experimental brain research 115, 47–60 (1997). 10.1007/PL00005684

22 Iremonger, K. J., Anderson, T. R., Hu, B. & Kiss, Z. H. T. Cellular Mechanisms Preventing Sustained Activation of Cortex During Subcortical High-Frequency Stimulation. Journal of neurophysiology 96, 613–621 (2006). 10.1152/jn.00105.2006

23 Bekar, L., et al. Adenosine is crucial for deep brain stimulation–mediated attenuation of tremor. Nature medicine 14, 75–80 (2008). 10.1038/nm1693

24 Preszler, C., Stieger, K., Chen, K., Zhang, G. & Kozai, T. Microglia surveillance is directed toward neuron activation during sustained intracortical microstimulation. Journal of Neural Engineering (2026).

25 Chen, K., Stieger, K. C. & Kozai, T. D. Challenges and opportunities of advanced gliomodulation technologies for excitation-inhibition balance of brain networks. Current Opinion in Biotechnology 72, 112–120 (2021).

26 Wellman, S. M., et al. Dynamic changes in the structure and function of brain mural cells around chronically implanted microelectrodes. Biomaterials 315, 122963 (2025).

27 Rakymzhan, A., et al. Parvalbumin interneurons drive depth-dependent vascular responses. iScience 28 (2025).

28 Krawchuk, M. B., Ruff, C. F., Yang, X., Ross, S. E. & Vazquez, A. L. Optogenetic assessment of VIP, PV, SOM and NOS inhibitory neuron activity and cerebral blood flow regulation in mouse somato-sensory cortex. Journal of Cerebral Blood Flow & Metabolism 0, 0271678X19870105 (2019). 10.1177/0271678x19870105

29 Fenoy, A. J., Goetz, L., Chabardes, S. & Xia, Y. Deep brain stimulation: are astrocytes a key driver behind the scene? CNS neuroscience & therapeutics 20, 191–201 (2014). 10.1111/cns.12223

30 Hess, C. W., Vaillancourt, D. E. & Okun, M. S. The temporal pattern of stimulation may be important to the mechanism of deep brain stimulation. Exp Neurol 247, 296–302 (2013). 10.1016/j.expneurol.2013.02.001

31 Vedam-Mai, V., et al. Deep brain stimulation and the role of astrocytes. Molecular psychiatry 17, 124–131, 115 (2012). 10.1038/mp.2011.61

32 Covelo, A. & Araque, A. Neuronal activity determines distinct gliotransmitter release from a single astrocyte. eLife 7, e32237 (2018). 10.7554/eLife.32237

33 Lines, J., Martin, E. D., Kofuji, P., Aguilar, J. & Araque, A. Astrocytes modulate sensory-evoked neuronal network activity. Nature communications 11, 3689 (2020). 10.1038/s41467-020-17536-3

34 Nam, Y., et al. Reversible Induction of Pain Hypersensitivity following Optogenetic Stimulation of Spinal Astrocytes. Cell reports 17, 3049–3061 (2016). 10.1016/j.celrep.2016.11.043

35 Xu, Q., et al. Astrocytes contribute to pain gating in the spinal cord. Science Advances 7, eabi6287 (2021).

36 Parker, J. L., Karantonis, D. M., Single, P. S., Obradovic, M. & Cousins, M. J. Compound action potentials recorded in the human spinal cord during neurostimulation for pain relief. Pain 153, 593–601 (2012).

37 Capogrosso, M., et al. A computational model for epidural electrical stimulation of spinal sensorimotor circuits. The Journal of Neuroscience 33, 19326–19340 (2013).

38 Ma, Z., et al. Two-photon calcium imaging of neuronal and astrocytic responses: the influence of electrical stimulus parameters and calcium signaling mechanisms. Journal of Neural Engineering (2021).

39 Grundfest, D., et al. Spatiotemporally distinctive astrocytic and neuronal responses to repetitive intracortical microstimulation. Journal of Neural Engineering 23, 046010 (2026).

40 Kofuji, P. & Araque, A. G-protein-coupled receptors in astrocyte–neuron communication. Neuroscience 456, 71–84 (2021).

41 Araque, A., et al. Gliotransmitters Travel in Time and Space. Neuron 81, 728–739 (2014). 10.1016/j.neuron.2014.02.007

42 Hughes, C. L., Stieger, K. C., Chen, K., Vazquez, A. L. & Kozai, T. D. Spatiotemporal properties of cortical excitatory and inhibitory neuron activation by sustained and bursting electrical microstimulation. iScience 28 (2025).

43 Stieger, K. C., Eles, J. R., Ludwig, K. A. & Kozai, T. D. Intracortical microstimulation pulse waveform and frequency recruits distinct spatiotemporal patterns of cortical neuron and neuropil activation. Journal of neural engineering 19, 026024 (2022).

44 Khakh, B. S. & McCarthy, K. D. Astrocyte calcium signaling: from observations to functions and the challenges therein. Cold Spring Harbor Perspectives in Biology 7, a020404 (2015).

45 Ghézali, G., Dallérac, G. & Rouach, N. Perisynaptic astroglial processes: dynamic processors of neuronal information. Brain Structure and Function 221, 2427–2442 (2016).

46 Lines, J., et al. A spatial threshold for astrocyte calcium surge. Elife 12, RP90046 (2024).

47 Semyanov, A. & Verkhratsky, A. Astrocytic processes: from tripartite synapses to the active milieu. Trends in Neurosciences 44, 781–792 (2021). 10.1016/j.tins.2021.07.006

48 Zhang, Y., et al. Fast and sensitive GCaMP calcium indicators for imaging neural populations. Nature 615, 884–891 (2023). 10.1038/s41586-023-05828-9

49 Dana, H., et al. High-performance calcium sensors for imaging activity in neuronal populations and microcompartments. Nature Methods 16, 649–657 (2019). 10.1038/s41592-019-0435-6

50 Yu, X., Moye, S. L. & Khakh, B. S. Local and CNS-Wide Astrocyte Intracellular Calcium Signaling Attenuation In Vivo with CalExflox Mice. Journal of Neuroscience 41, 4556–4574 (2021).

51 Yu, X., et al. Reducing Astrocyte Calcium Signaling In Vivo Alters Striatal Microcircuits and Causes Repetitive Behavior. Neuron 99, 1170–1187.e1179 (2018). 10.1016/j.neuron.2018.08.015

52 Durkee, C. A., et al. Gi/o protein-coupled receptors inhibit neurons but activate astrocytes and stimulate gliotransmission. Glia 67, 1076–1093 (2019).

53 Nagai, J., et al. Specific and behaviorally consequential astrocyte Gq GPCR signaling attenuation in vivo with iβARK. Neuron 109, 2256–2274. e2259 (2021).

54 Delepine, C., Shih, J., Li, K., Gaudeaux, P. & Sur, M. Differential effects of astrocyte manipulations on learned motor behavior and neuronal ensembles in the motor cortex. Journal of Neuroscience 43, 2696–2713 (2023).

55 Vaidyanathan, T. V., Collard, M., Yokoyama, S., Reitman, M. E. & Poskanzer, K. E. Cortical astrocytes independently regulate sleep depth and duration via separate GPCR pathways. Elife 10, e63329 (2021).

56 Lai, B., et al. Astrocytic Ca2+ prevents synaptic depotentiation by limiting repetitive activity in dendrites during motor learning. Nature Neuroscience 28, 2296–2309 (2025).

57 Waters, J. Sources of widefield fluorescence from the brain. Elife 9, e59841 (2020).

58 Mederos, S., et al. Melanopsin for precise optogenetic activation of astrocyte-neuron networks. Glia 67, 915–934 (2019).

59 Octeau, J. C. et al. Transient, Consequential Increases in Extracellular Potassium Ions Accompany Channelrhodopsin2 Excitation. Cell reports 27, 2249–2261.e2247 (2019). 10.1016/j.celrep.2019.04.078

60 Meeks, J. P. & Mennerick, S. Selective effects of potassium elevations on glutamate signaling and action potential conduction in hippocampus. The Journal of neuroscience : the official journal of the Society for Neuroscience 24, 197–206 (2004). 10.1523/jneurosci.4845-03.2004

61 Mederos, S. & Perea, G. GABAergic-astrocyte signaling: A refinement of inhibitory brain networks. Glia 67, 1842–1851 (2019).

62 Kuncel, A. M., et al. Clinical response to varying the stimulus parameters in deep brain stimulation for essential tremor. Movement disorders : official journal of the Movement Disorder Society 21, 1920–1928 (2006). 10.1002/mds.21087

63 Figee, M., et al. Deep brain stimulation for depression. Neurotherapeutics 19, 1229–1245 (2022).

64 Fernández, E., et al. Visual percepts evoked with an Intracortical 96-channel microelectrode array inserted in human occipital cortex. The Journal of Clinical Investigation (2021).

65 Kozai, T. D. Y., et al. Solving the problem of inception: A cross-species perspective on strategies for a mechanistic refinement of intracortical microstimulation. Journal of Neural Engineering 23, 0230001 (2026).

66 Bazargani, N. & Attwell, D. Astrocyte calcium signaling: the third wave. Nature Neuroscience 19, 182–189 (2016). 10.1038/nn.4201

67 Agulhon, C., et al. Calcium Signaling and Gliotransmission in Normal vs. Reactive Astrocytes. Front Pharmacol 3, 139 (2012). 10.3389/fphar.2012.00139

68 Cavaccini, A., Durkee, C., Kofuji, P., Tonini, R. & Araque, A. Astrocyte Signaling Gates Long-Term Depression at Corticostriatal Synapses of the Direct Pathway. The Journal of Neuroscience 40, 5757 (2020). 10.1523/JNEUROSCI.2369-19.2020

69 Ding, S., et al. Enhanced astrocytic Ca2+ signals contribute to neuronal excitotoxicity after status epilepticus. The Journal of neuroscience : the official journal of the Society for Neuroscience 27, 10674–10684 (2007). 10.1523/jneurosci.2001-07.2007

70 Tan, Z., et al. Glia-derived ATP inversely regulates excitability of pyramidal and CCK-positive neurons. Nature communications 8, 13772 (2017). 10.1038/ncomms13772

71 de Ceglia, R., et al. Specialized astrocytes mediate glutamatergic gliotransmission in the CNS. Nature, 1–10 (2023).

72 Perea, G., et al. Activity-dependent switch of GABAergic inhibition into glutamatergic excitation in astrocyte-neuron networks. eLife 5, e20362 (2016). 10.7554/eLife.20362

73 McMahon, S. M. & Jackson, M. B. An inconvenient truth: calcium sensors are calcium buffers. Trends in neurosciences 41, 880–884 (2018).

74 Semyanov, A., Henneberger, C. & Agarwal, A. Making sense of astrocytic calcium signals — from acquisition to interpretation. Nature Reviews Neuroscience 21, 551–564 (2020). 10.1038/s41583-020-0361-8

75 Lee, S. H., Rosenmund, C., Schwaller, B. & Neher, E. Differences in Ca2+ buffering properties between excitatory and inhibitory hippocampal neurons from the rat. The Journal of physiology 525, 405–418 (2000).

76. Eles, J. R. & Kozai, T. D. Y. in Biomaterials Vol. 234 (Elsevier Ltd, 2020).

77 Suematsu, N., Vazquez, A. L. & Kozai, T. D. Activation and depression of neural and hemodynamic responses induced by the intracortical microstimulation and visual stimulation in the mouse visual cortex. Journal of Neural Engineering 21, 026033 (2024).

78 Tehovnik, E. J., Tolias, A. S., Sultan, F., Slocum, W. M. & Logothetis, N. K. Direct and indirect activation of cortical neurons by electrical microstimulation. Journal of neurophysiology 96, 512–521 (2006). 10.1152/jn.00126.2006

79 Eles, J. R. & Kozai, T. D. Y. In vivo imaging of calcium and glutamate responses to intracortical microstimulation reveals distinct temporal responses of the neuropil and somatic compartments in layer II/III neurons. Biomaterials 234, 119767 (2020). 10.1016/j.biomaterials.2020.119767

80 Histed, M. H., Bonin, V. & Reid, R. C. Direct activation of sparse, distributed populations of cortical neurons by electrical microstimulation. Neuron 63, 508–522 (2009). 10.1016/j.neuron.2009.07.016

81 Yi, G. & Grill, W. M. Frequency-dependent antidromic activation in thalamocortical relay neurons: effects of synaptic inputs. J Neural Eng 15, 056001 (2018). 10.1088/1741-2552/aacbff

82 Hughes, C. L., et al. Perception of microstimulation frequency in human somatosensory cortex. Elife 10, e65128 (2021).

83 Savtchouk, I. & Volterra, A. Gliotransmission: beyond black-and-white. The Journal of Neuroscience 38, 14–25 (2018).

84 Verkhratsky, A. & Nedergaard, M. Physiology of astroglia. Physiological reviews 98, 239–389 (2018).

85 Goenaga, J., Araque, A., Kofuji, P. & Herrera Moro Chao, D. Calcium signaling in astrocytes and gliotransmitter release. Frontiers in Synaptic Neuroscience **Volume** 15 - 2023 (2023). 10.3389/fnsyn.2023.1138577

86 Giaume, C. & Venance, L. Intercellular calcium signaling and gap junctional communication in astrocytes. Glia 24, 50–64 (1998).

87 Onodera, M., et al. Exacerbation of Epilepsy by Astrocyte Alkalization and Gap Junction Uncoupling. The Journal of Neuroscience 41, 2106 (2021). 10.1523/JNEUROSCI.2365-20.2020

88 Eles, J. R., Stieger, K. C. & Kozai, T. D. Y. The temporal pattern of Intracortical Microstimulation pulses elicits distinct temporal and spatial recruitment of cortical neuropil and neurons. J Neural Eng (2020). 10.1088/1741-2552/abc29c

89 Stieger, K. C., Eles, J. R., Ludwig, K. A. & Kozai, T. D. Y. In vivo microstimulation with cathodic and anodic asymmetric waveforms modulates spatiotemporal calcium dynamics in cortical neuropil and pyramidal neurons of male mice. Journal of neuroscience research 98, 2072–2095 (2020). 10.1002/jnr.24676

90 Mastakov, M. Y., Baer, K., Xu, R., Fitzsimons, H. & During, M. J. Combined injection of rAAV with mannitol enhances gene expression in the rat brain. Molecular therapy 3, 225–232 (2001).

91 Bareyre, F., Wahl, F., McINTOSH, T. K. & Stutzmann, J.-M. Time course of cerebral edema after traumatic brain injury in rats: effects of riluzole and mannitol. Journal of neurotrauma 14, 839–849 (1997).

92 McCreery, D., Pikov, V. & Troyk, P. R. Neuronal loss due to prolonged controlled-current stimulation with chronically implanted microelectrodes in the cat cerebral cortex. J Neural Eng 7, 036005 (2010). 10.1088/1741-2560/7/3/036005

93 Huang, Y.-Z., Edwards, M. J., Rounis, E., Bhatia, K. P. & Rothwell, J. C. Theta burst stimulation of the human motor cortex. Neuron 45, 201–206 (2005).

94 Rakymzhan, A., et al. Parvalbumin interneurons drive depth-dependent vascular responses. iScience (2025).

95 Thompson, K. J., et al. DREADD agonist 21 is an effective agonist for muscarinic-based DREADDs in vitro and in vivo. ACS Pharmacology & Translational Science 1, 61–72 (2018).

96 Pachitariu, M., et al. Suite2p: beyond 10,000 neurons with standard two-photon microscopy. BioRxiv, 061507 (2016).

97 Schindelin, J., et al. Fiji: an open-source platform for biological-image analysis. Nature methods 9, 676–682 (2012).

98 Ma, Y., et al. Resting-state hemodynamics are spatiotemporally coupled to synchronized and symmetric neural activity in excitatory neurons. Proceedings of the National Academy of Sciences of the United States of America 113, E8463–e8471 (2016). 10.1073/pnas.1525369113

99 Bates, D., Mächler, M., Bolker, B. & Walker, S. Fitting Linear Mixed-Effects Models Using lme4. Journal of Statistical Software 67, 1–48 (2015). 10.18637/jss.v067.i01

