## Supplementary Materials for "Astrocyte modulation of neural activity during intracortical microstimulation"

Supplemental Information: Astrocyte modulation of neural activity during intracortical microstimulation

**Supplementary Information:**

Suppemental Table 1 Animal numbers

| **Animal model** | **Animal numbers** | **Relevant Figures** |
| --- | --- | --- |
| Aldh1l1-Cre: Syn-jRGeco1a + Flex-GCaMP8m | 1F, 2M = 3 | Fig. 1,2, Fig. S1-S6 |
| Aldh1l1-Cre: Flex-GCaMP8m + GFAP-hM3D(Gq)-mCherry | 1F, 2M = 3 | Fig. 3,4, Fig. S7, S10 |
| Aldh1l1-Cre: Flex-GCaMP8m + GFAP104-mCherry (Validation) | 1F,1M = 2 | Fig. S7, S10 |
| C57BL6J: Syn-GCaMP8m + GFAP-hM3D(Gq)-mCherry | 2F, 2M = 4 | Fig. 3,4, Fig. S8 |
| C57BL6J: Syn-GCaMP8m + GFAP104-mCherry (Validation) | 1M,1F = 2 | Fig. 4, Fig. S8 |
| Aldh1l1-Cre: Flex-GCaMP8m + GFAP104-Melanopsin-mCherry | 1F, 2M = 3 | Fig. 5,6, Fig. S11, S14 |
| Aldh1l1-Cre: Flex-GCaMP8m + GFAP104-mCherry (Validation) | 1F, 1M = 2 | Fig. S11, S14 |
| C57BL6J: Syn-GCaMP8m + GFAP104-Melanopsin-mCherry | 2F, 2M = 4 | Fig. 5,6, Fig. S12, S13 |
| C57BL6J: Syn-GCaMP8m + GFAP104-mCherry (Validation) | 1M, 1F = 2 | Fig. 6, Fig. S12 |

**Supplemental Figures:**

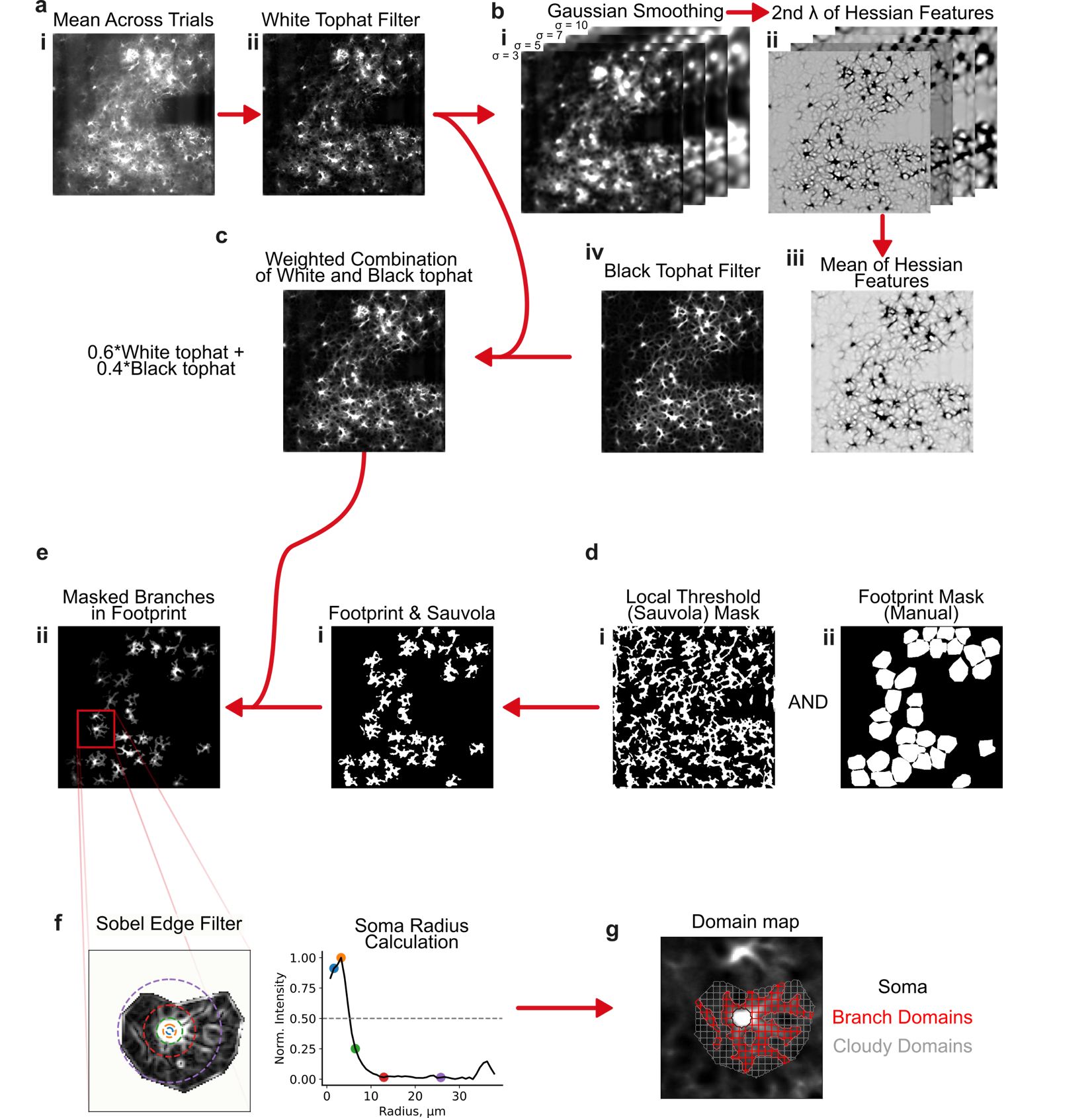

**Figure S1. Preprocessing of images for semiautomatic segmentation of astrocyte soma from their branches and cloudy regions. A**. A white top hat filter was applied to the trial averaged mean image for a given animal (ai) which helped to highlight astrocyte morphology from the background. **B**. The white tophat filtered image was smoothed with 4 different gaussian filters (i) and the second eigen value of the hessian feature matrix of those smoothed images (ii) were averaged together (iii). The sigmas for the gaussian smoothing were chosen to highlight astrocyte morphology without over-enhancing fine branching. A black tophat filter was used to highlight dark features of the hessian output. **C**. A weighted average of the hessian output and the white tophat filter was used to highlight original astrocyte morphology without overemphasizing hessian features. **D**. A Sauvola local threshold was used to identify bright regions representing astrocyte morphology. This threshold was logically combined with a region mask that was manually defined around identified astrocytes that contained clear soma and main branches (dii). **E.** The astrocyte footprint mask was merged with the weighted image. **F**. A Sobel edge filter was applied to a masked astrocyte region to highlight edges of the soma. The mean intensity of radial bins extending from the manually defined soma center was used to identify the edge of the soma, where the soma radius was defined as the distance where the mean intensity of a radial bin dropped below half of its maximal value. **G**. A circular ROI with radius defined in f, represented the soma, the thresholded region in (ei) excluding the soma was defined as the main branch region, and the remaining area in the manually defined astrocyte region was defined as the cloud region. The branches and cloud regions were then broken into domains of maximal size of ~3x3 µm.

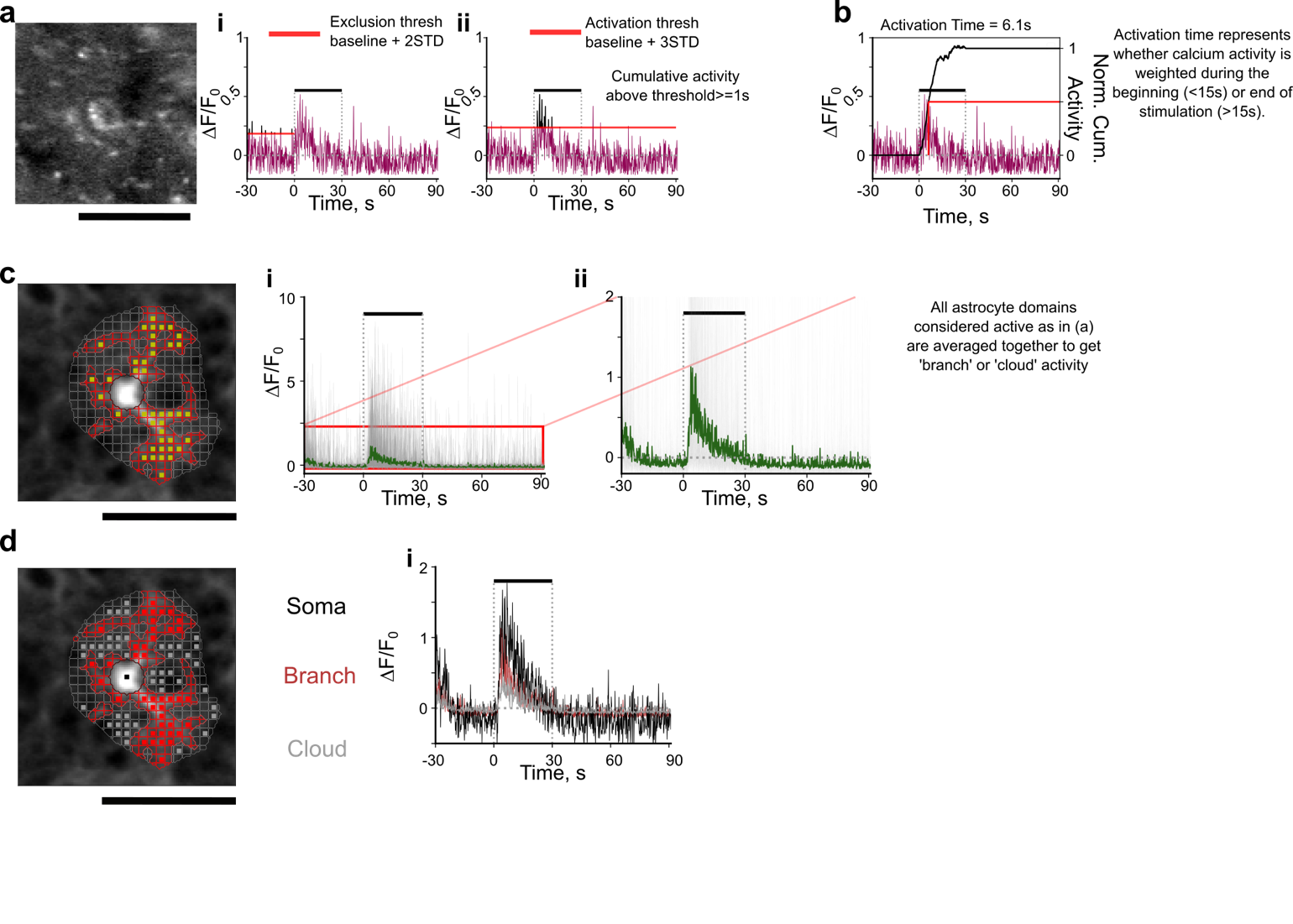

**Figure S2. Demonstration of determination of activation for astrocytes or neurons.**

**A.** example neuron (left) and corresponding trace during stimulation for a given 100 Hz stimulation. To exclude certain potential activity from computation of noise floor baseline, activity above 2 standard deviations during the baseline period (threshold in red, activity above initial threshold represented in black) were excluded to recompute a more representative baseline and standard deviation of the noise level (i). The threshold was computed as the mean + 3 standard deviations above the recomputed baseline activity (red horizontal line). If neuronal or astrocyte calcium was elevated above this threshold for a total of 0.5, or 1s for neurons or astrocytes, respectively that neuron, or astrocyte domain was concluded to be activated for the given trial. Activity above threshold is represented in black. **B**. computation of activation time for the given neuron. Activation time represents the amount of time it takes to reach half the total cumulative calcium activity during stimulation. Black line (right axis) represents the normalized cumulative sum of activity during stimulation. Activation time is the time that the cumulative line reached half of total cumulative activity. **C**. Representative astrocyte and corresponding segmented domains (left). Yellow boxes represent branch domain regions determined to be activated during a given 100 Hz trial using method in (a). Their corresponding activity is represented by gray lines in (i). Because activity in these small domains were variable, they were averaged together to get activity for the branch domain (ii) showing clear elevation in calcium activity during stimulation. **D**. Representative image of the same astrocyte and corresponding activated soma, branch or cloud domains deemed activated (left), and their corresponding averaged activity for the 100 Hz stimulation trial.

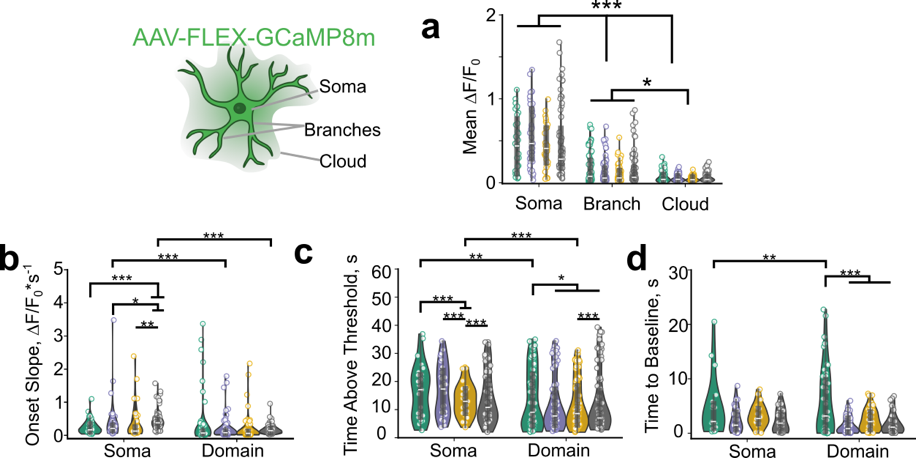

**Figure S3) ICMS influences astrocyte calcium activity depending on stimulation pattern and compartment. A)** (Left) Graphic representing different astrocyte compartments. ‘Domain’ represents the average of branch and cloud regions. (Right) Astrocyte soma display the significantly larger mean calcium activity in response to ICMS compared to branch and cloud domains (LMM; Stim: 2.8e-85, ROI type: p = 8.0e-16, interaction: p=8.7e-57). Activated branch and cloud domains were averaged together for future analyses. **B)** The activation slope of calcium activity in the somata is influenced by stimulation pattern (LMM; Stim: 3.8e-3, ROI type: p=2.0e-4, interaction: p=6.1e-3, trial: p=0.53). **C)** Stimulation parameters significantly influence the time of activity above threshold of astrocyte somata and domains (LMM; Stim: 3.6e-26, ROI type: p=0.06, interaction: p=5.7e-8, trial: p=0.04). **D)** Astrocytes display sustained calcium elevations after stimulation ends that is influenced by stimulation pattern in the domains (LMM; Stim: 4.1e-140, ROI type: p=9.5e-4, interaction: p=0.03, trial: p=0.22). N=3, n=31-82, 6 trials.

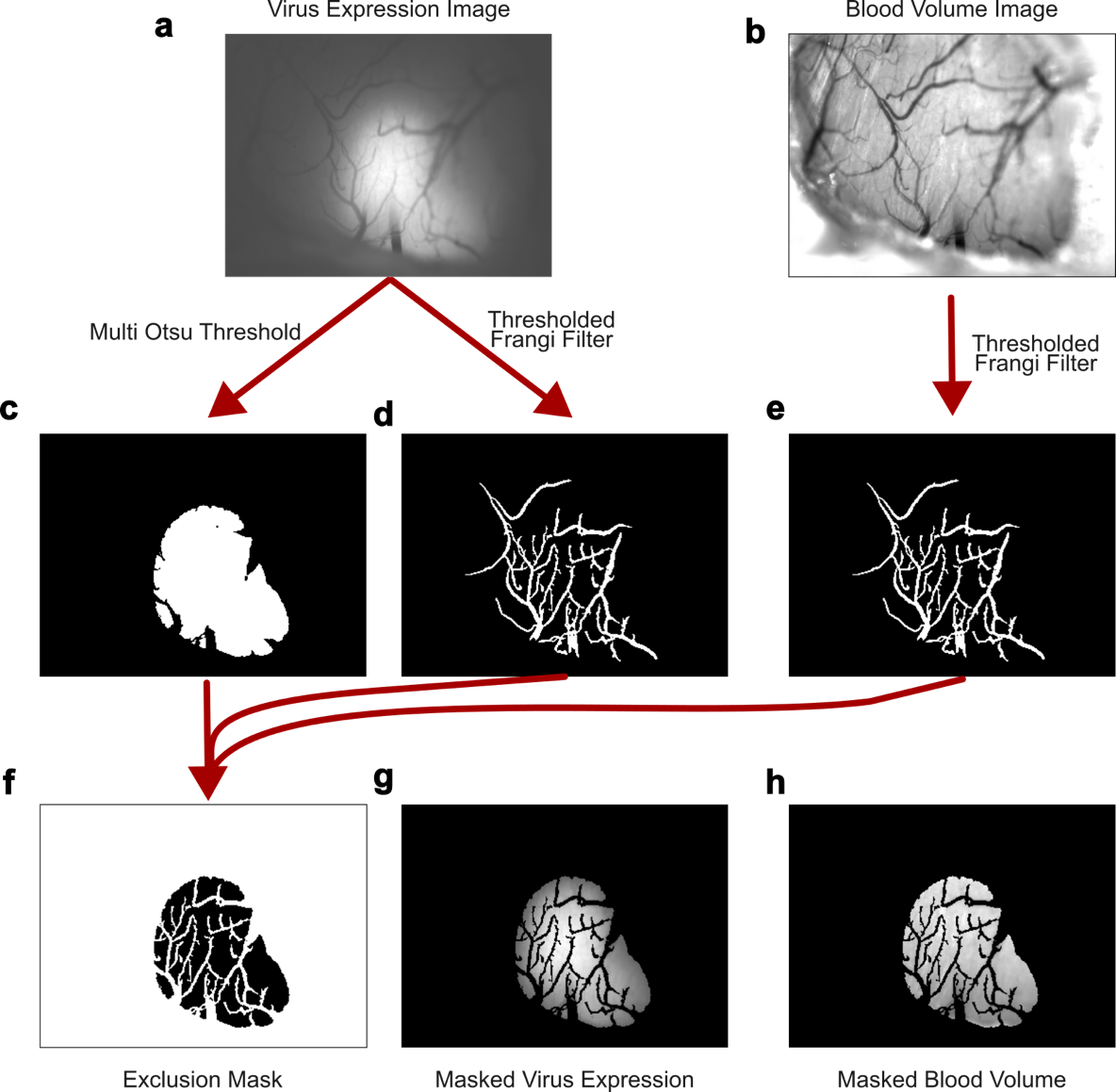

**Figure S4. Method for mesoscale region and vessel masking.** **A.** Representative average projection of virus expressing region. **B**. Average projection of raw green LED reflectance (for blood volume calculation). **C**. Area including virus expression was identified by selecting the connected pixels above the second threshold of a multi-Otsu threshold with 3 regions. **D-E**. Thresholding a Frangi filter applied to the virus expressing image and the blood volume image highlighted the large blood vessels on the brain surface. **F**. A mask to exclude non-expressing regions and blood vessels was created through a logical combinations of the masks in C-E. **G-H**. Representative virus expression and blood volume image masked by the exclusion mask.

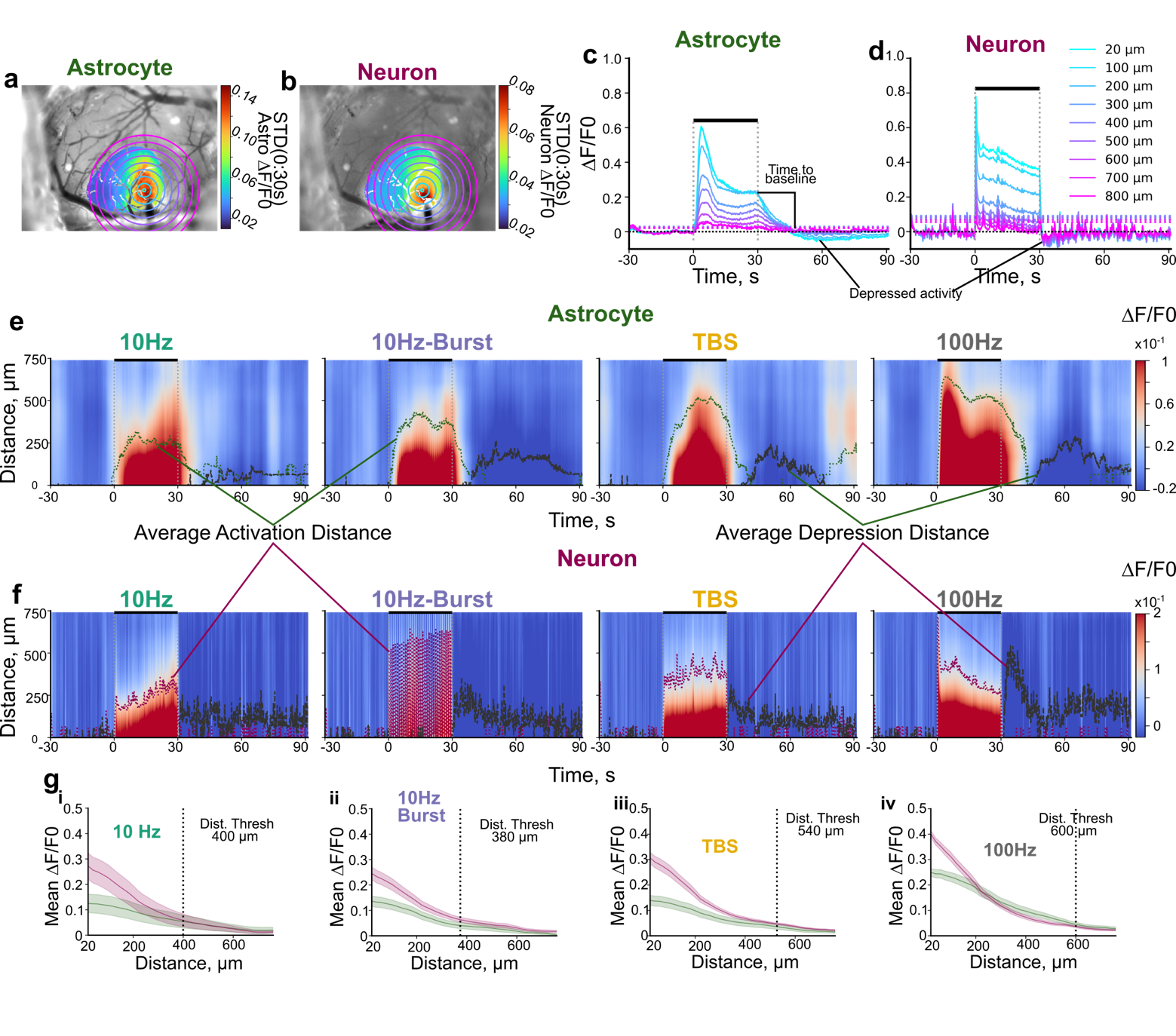

**Figure S5. Identification of ‘activated’ regions in mesoscale imaging.** A-b. Standard deviation projections within the masked regions for astrocyte (a) or neuronal (b) activation during a 100 Hz stimulation trial with radial bins drawn relative to the center of neuronal activation. C-d. Corresponding traces and activation threshold (dotted horizontal lines of corresponding color) for radial bins drawn in a-b for astrocytes (c) or neurons (d). Activity traces often depress below baseline (black horizontal line) after stimulation cessation. E-f Average heatmap projections across animals and trials over time and distance highlighting periods of high activity (red) during stimulation and depressed activity (blue) after stimulation ends for astrocytes (e) and neurons (f). Green and red traces highlight the average activation distance across animals over time for astrocytes and neurons. Dark gray traces highlight the average distance for depressed activity, which elevates after stimulation ends. g. Average ± standard error of activity over distance for different stimulation paradigms. The average distance to compare astrocyte and neuronal activity patterns represents the average activation area + 1 STD across animals and cell types for each stimulation condition.

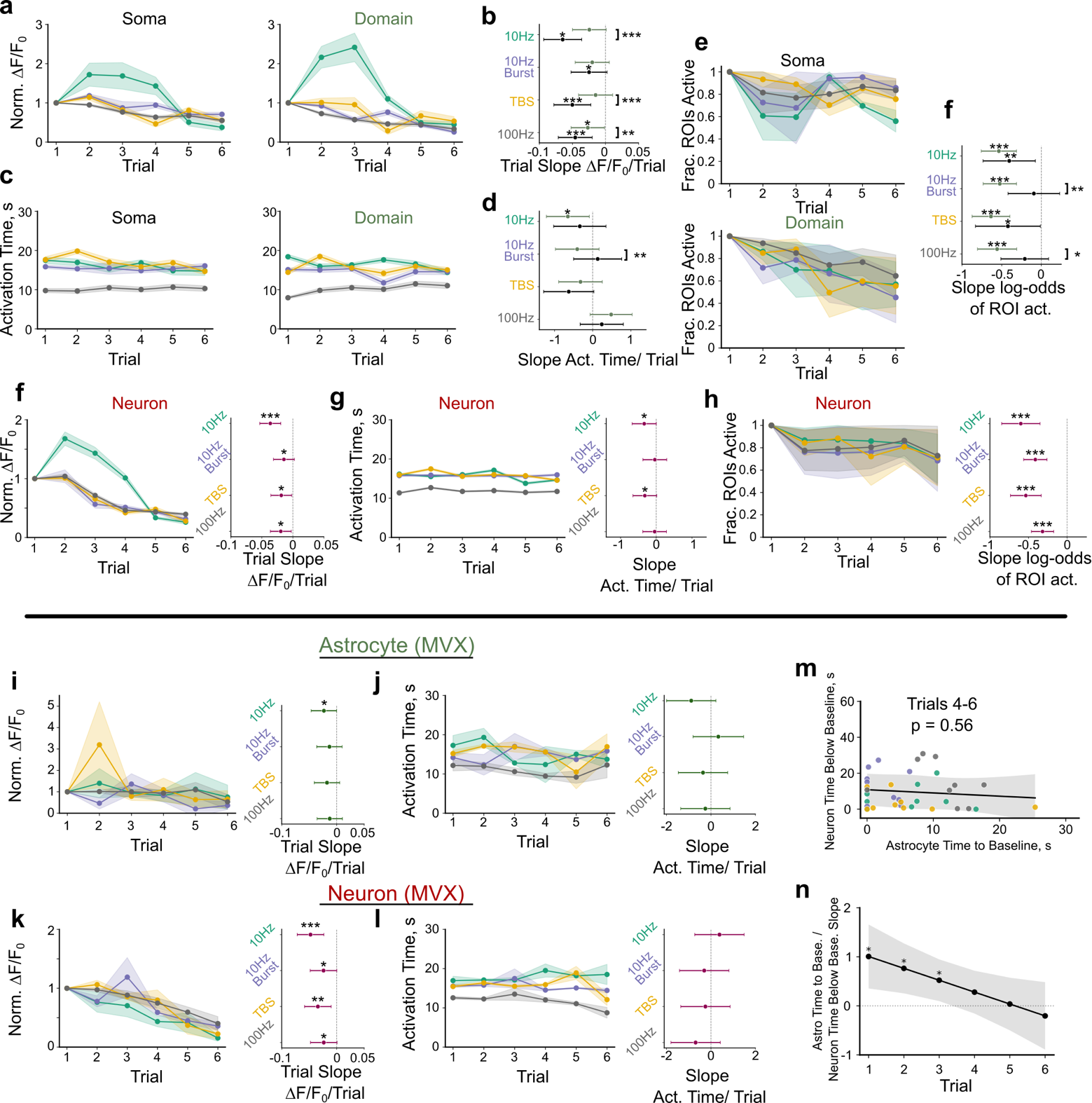

**Figure S6. Magnitude of activation decays over trials, with temporal structure largely intact.** A-b. Magnitude of astrocyte soma and domains reduces over trials (LMM Stim: p = 9.8e-87, compartment: p = 1.4e-9, trial: p = 5.6e-4, Stim x compartment: p = 1.9e-50, Stim x trial: p = 7.5e-3, domain x trial: p = 2.4e-10, N=3, n=31 - 82). Fitted slope ± 95% CI of magnitude of activation shows dependence on stimulation parameter and astrocyte compartment (b). C-d. Activation time of astrocyte compartments is largely consistent over trials LMM Stim: p = 4.1e-140, compartment: p = 9.5e-4, trial: p = 0.23, Stim x compartment: p = 0.03, Stim x trial: p = 1.0e-12, domain x trial: p = 0.73 N=3, n=31 - 82). D Fitted slope ± 95% CI of trial effect shows dependence on stimulation pattern and astrocyte compartment. E-f. Fraction of activated astrocyte compartment ROIs decays with trial. Fitted slope of the log-odds of an astrocyte ROI being active over trials shows decay with trial (Binomial GLMM: Stim: p= 1.7e-4, Compartment: p = 0.14, trial: p = 6.4e-16, stim x compartment: p=0.14, stim x trial: p = 0.35, compartment x trial: p = 1.1e-4). F. Magnitude of neuronal ROIs decays with trial (LMM: Stim 9.6e-137, Trial: p = 2.0e-3, Stim x trial1.6e-43; N = 3, n = 215-280). G. temporal structure of neuronal activity changes in some stimulation parameters (See Fig 2 LMM Stim: p = 1.5e-259, ROI type 0.61, trial p=0.06, Stim x roi type: p = 2.1e-22, Stim x trial: p= 6.0e-11, roi type x trial: p = 0.79). H. Fraction of active neuron ROIs decays with trial (GLMM Stim: p = 1.4e-3, Trial: p = 1.7e-9, Stim x trial: p = 0.037). *Right:* Slope of the log-odds of an ROI being active. I Magnitude of mesoscale astrocyte activity decays over trials (LMM Stim: p = 2.9e-3, Trial: p = 0.012, stim x trial: 0.76). j. Temporal structure of mesoscale astrocyte activity may change over trial (See Fig 2m stats: no significant posthoc decay with trial). K Magnitude of mesoscale neuronal activity reduces over trials (LMM Stim: p = 0.86, Trial p = 3.7e-5, Stim x trial p = 0.19). L. Temporal structure of neuronal activity is largely consistent over trials (see Fig 2m for stats, no significant posthoc decay with trial). M-N. The association of astrocyte time to baseline with neuron suppression depends on trial (See Fig 2o for stats). The relationship is not present for last three trials (M).

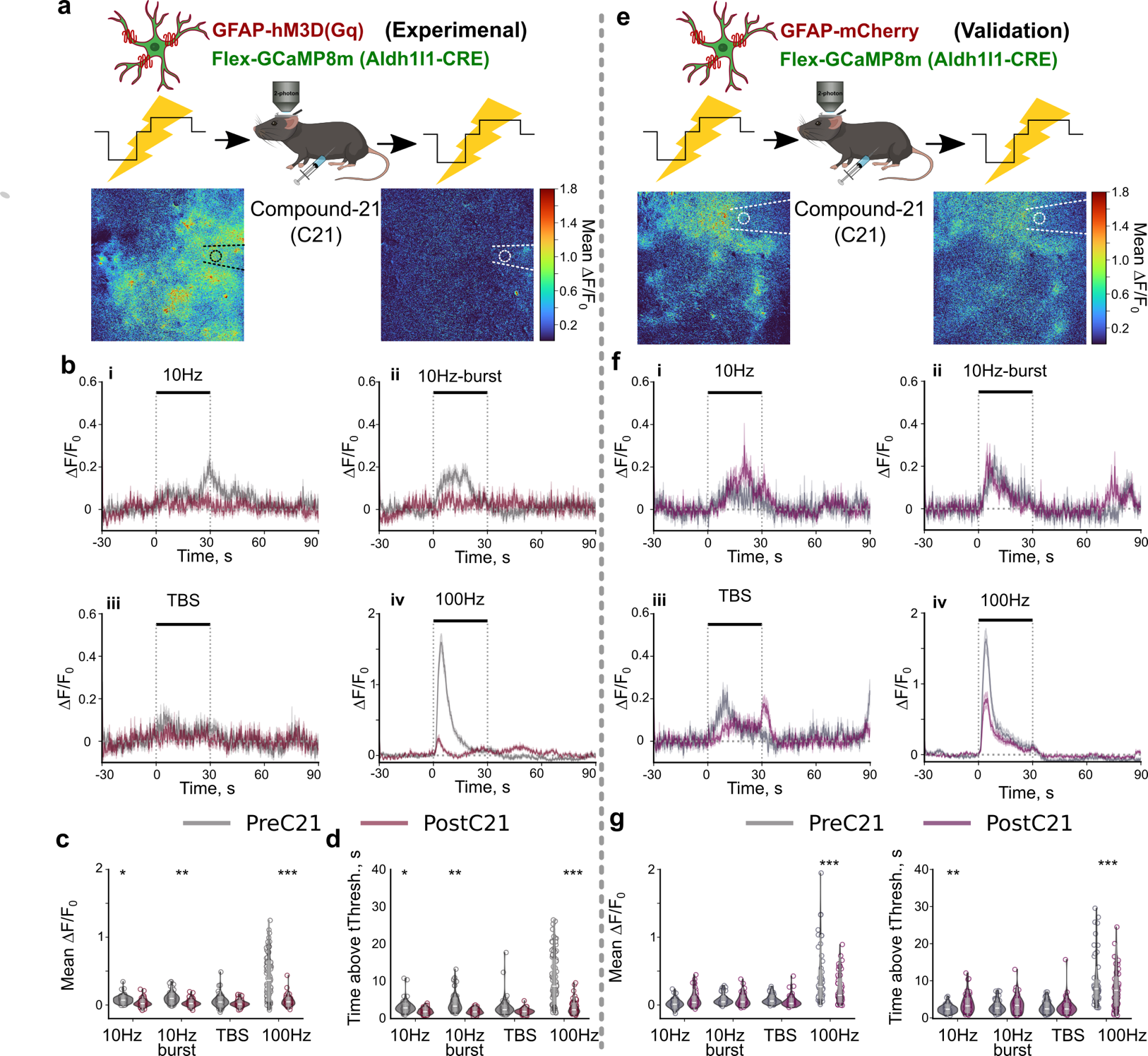

**Figure S7. Gq DREADD activation in astrocytes reduces stimulation induced calcium activity under two-photon imaging.** **A**. Aldh1l1-mice expressing GCaMP8m and hM3Dq in astrocytes received intracortical microstimulation before and after administration of 1mg/kg of the DREADD agonist C21. Images show the reduced average magnitude of stimulation induced calcium activity during 100 Hz stimulation following administration of C21. **B**. Trial averaged traces ± standard error across cells across stimulation paradigms demonstrate that stimulation induced elevations in astrocyte calcium activity are substantially reduced following administration of C21. **C-D**. Mean stimulation induced astrocyte calcium activity (C, LMM; Stim: p=2.8e-108, C21: p= 9.2e-6, interaction: p=1.1e-79) and time above threshold (D, LMM; Stim: p=1.4e-113, C21: p= 1.2e-5, interaction: p= 1.6e-75) were significantly reduced following administration of C21. E-G similar analyses in validation animals expressing mCherry in place of hM3Dq did not display similar magnitudes of reduced stimulation-induced astrocyte calcium activity following administration of C21. **E-G.**  Validation animals lacking hM3D. **E**. Aldh1l1-mice expressing GCaMP8m and mCherry in astrocytes received intracortical microstimulation before and after administration of 1mg/kg of the DREADD agonist C21. Images show relatively comparable average magnitude of stimulation induced calcium activity during 100 Hz stimulation following administration of C21. **F**. Trial averaged traces of astrocyte calcium activity ± standard error across cells demonstrate that C21 administration without DREADD expression do not eliminate stimulation induced astrocyte calcium activity. **G-H**. Calcium activity was significantly reduced after C21 administration only during 100Hz stimulation (G; LMM; Stim: p=1.3e-82, C21: p= 4.4e-3, interaction: p= 3.4e-13), but time above threshold was either increased or decreased depending on stimulation pattern (H; LMM; Stim: p=1.5e-102, C21: p= 0.97, interaction: p=4.8e-9). N=2 animals across 3 trials, n = 48-50. *:p<0.05, **:p<0.01, ***:p<0.001.

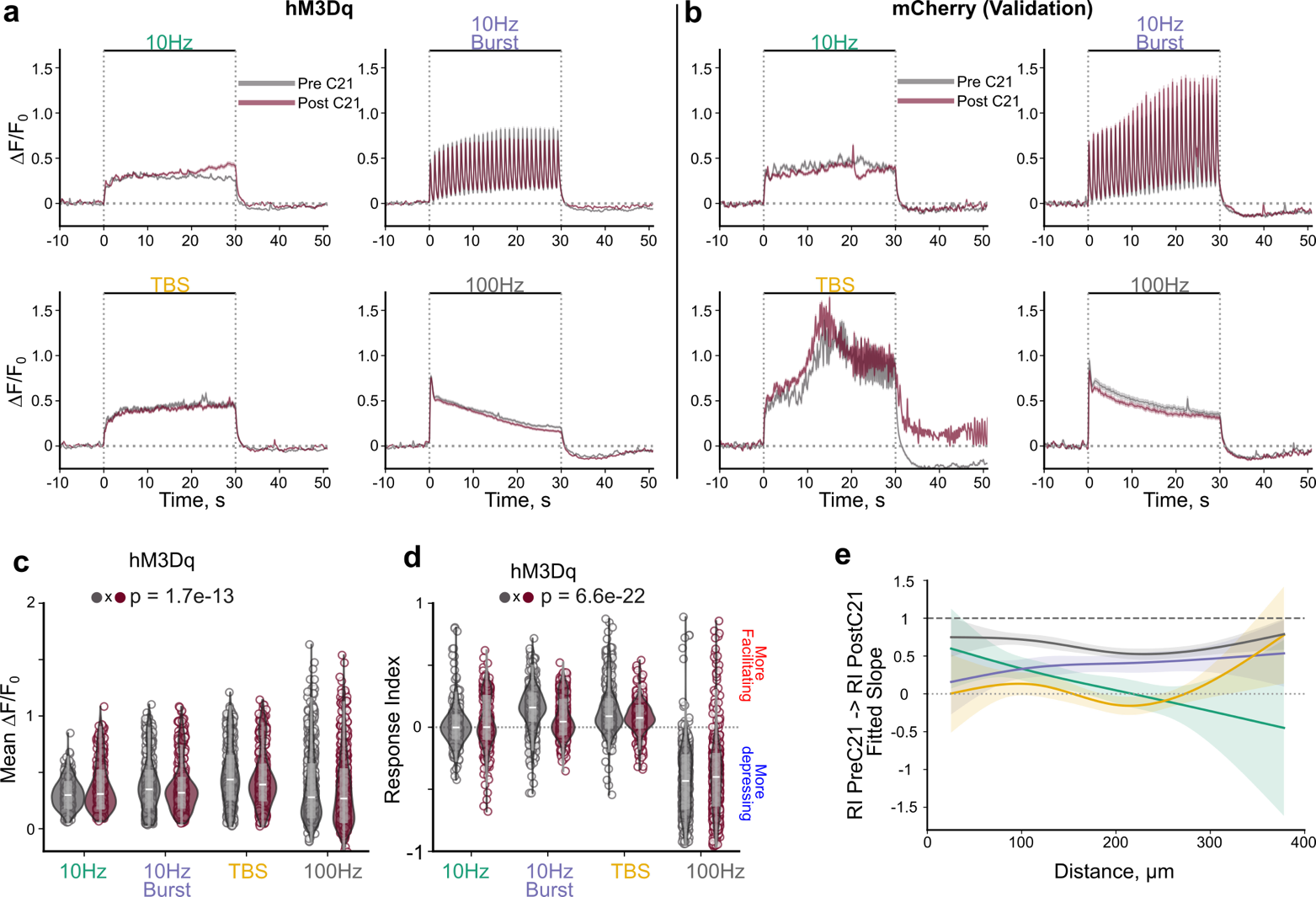

**Figure S8. Astrocyte calcium attenuation does not cause global shifts in stimulation induced neural activity.** **A.** Traces of neural activity before and after astrocyte attenuation. **B.** Traces of stimulation-induced activity in animals lacking hM3D in astrocytes show no major effects of C21 administration itself. **C**. Mean stimulation-induced activity before and after astrocyte attenuation (LMM: Stim: p<2.2e-16, C21: p=0.97, interaction: p=1.7e-13 N=4, n=403-713). **D.** Response index of neurons before and after attenuating astrocytes (LMM; Stim: p<2.2e-16, C21: p=0.99, interaction: p=6.6e-22, N=4, n= 403-713). **E**. Visualization of the relationship between the response index PreC21 to PostC21 of the form RI PostC21 ~ RI PreC21 * distance * stimcondition + (1|animalid). Plotting was computed by fitting a natural cubic spline to the centered distance to allow visualization of non-linear variance over distance. Shading is 95% confidence interval (LMM; PreC21 p=3.8e-11; distance: p=2.5e-3; Stim: p=8.5e-3; PreC21 x Distance: p= 1.8e-5, PreC21 x Stim: p = 0.05, Distance x Stim: 1.3e-3; Distance x PreC21 x Stim: p= 1.7e-12; N=4, n= 344-674).

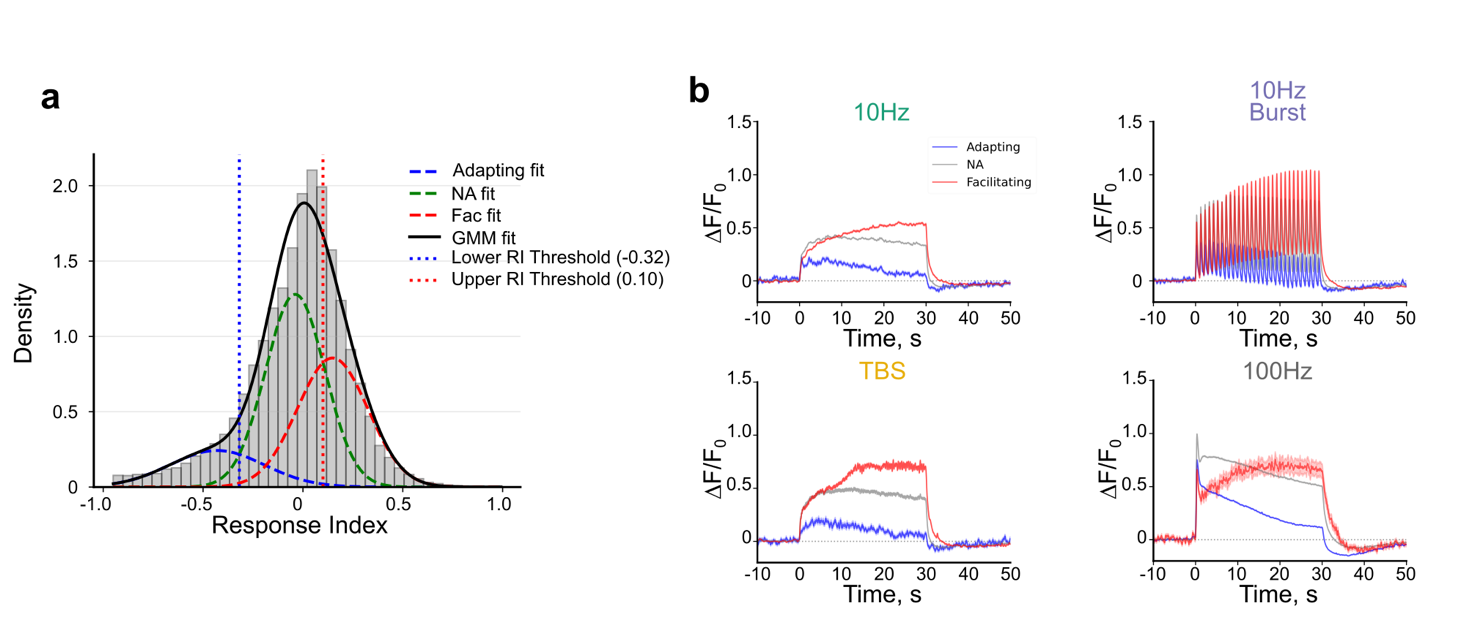

**Figure S9. Neuron responses to ICMS display three general profiles facilitating, non-adapting, and adapting.** A. A gaussian mixture model (GMM) of 3 distributions was fit to the response index of all neuron responses with GCaMP8m from this study (43,422 total responses across 1799 neurons from 12 animals). Data includes responses with hM3D and melanopsin manipulations and validation animals. B. Neuron responses (mean ± stderr shade) to each stimulation condition grouped by the respective GMM threshold highlights different classes of responses (facilitation: increase over time; non-adapting: small change over time, adapting: decrease over time).

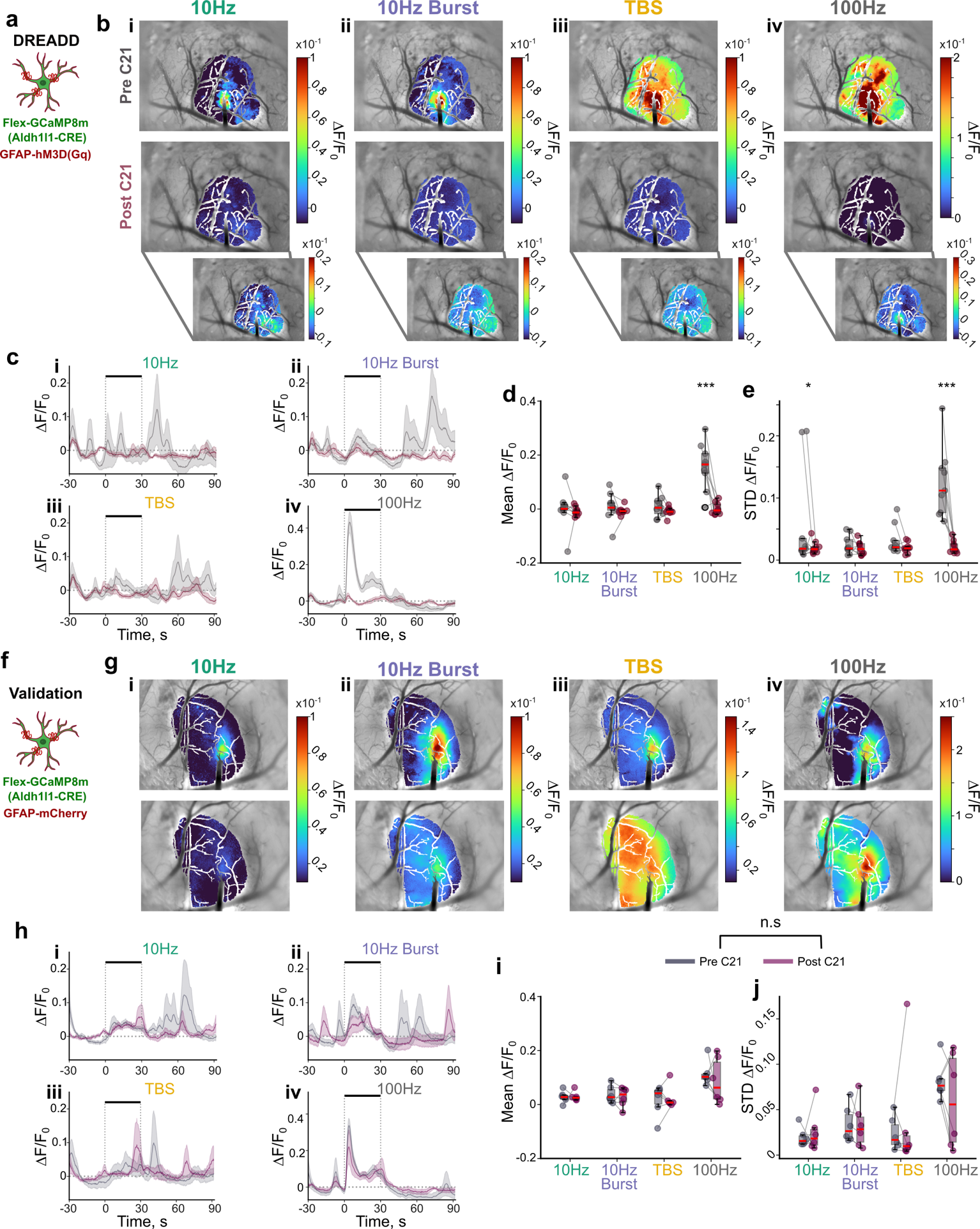

**Figure S10. GqDREADD activation in astrocytes by 1mg/kg C21 attenuates stimulation induced mesoscale astrocyte calcium activity. A**. An Aldh1l1-CRE animal was injected with FLEX-GCaMP8m and GFAP-hM3DGq-mCherry >4weeks before implantation of an electrode to express GCaMP and a Gq DREADD in astrocytes. **B**. Representative images highlighting the mean stimulation induced astrocyte calcium activity within the virus expressing region is virtually non-existent for all stimulation patterns, except 100 Hz. **C**. Trial averaged mesoscale astrocyte calcium activity ± standard error across animals is reduced following astrocyte DREADD activation (Pre-C21 in Gray, Post-C21 in red). (**D-E**) Mean (D; LMM; Stim: p=1.25e-6, C21: p=4.19e-5, interaction: p=1.51e-6) and standard deviation of calcium activity (E; LMM; Stim: p=4.8e-4, C21: p= 2.81e-5, interaction: p= 8.2e-4) during stimulation is significantly reduced following DREADD activation. **F**. Validation animals were created in which Aldl1l1-CRE animals express GCaMP8m in astrocytes and mCherry (GFAP-mCherry) in place of the Gq DREADD. **G**. Representative images of mean stimulation-induced astrocyte calcium activity within the virus expressing region are slightly reduced in some stimulation trials. **H**. Mean trial averaged mesoscale astrocyte calcium activity ± standard error across animals appears relatively unchanged following administration of C21. I-j) mean (I; LMM; Stim: p=2.4e-5, C21: p= 0.54, interaction: p=0.71) and standard deviation (j; LMM; Stim: p=2.7e-3, C21: p=0.94, interaction: p=0.55) of mesoscale astrocyte calcium activity is not significantly attenuated following administration of C21. N=2-3 animals across 3 trials *:p<0.05, **:p<0.01, ***:p<0.001.

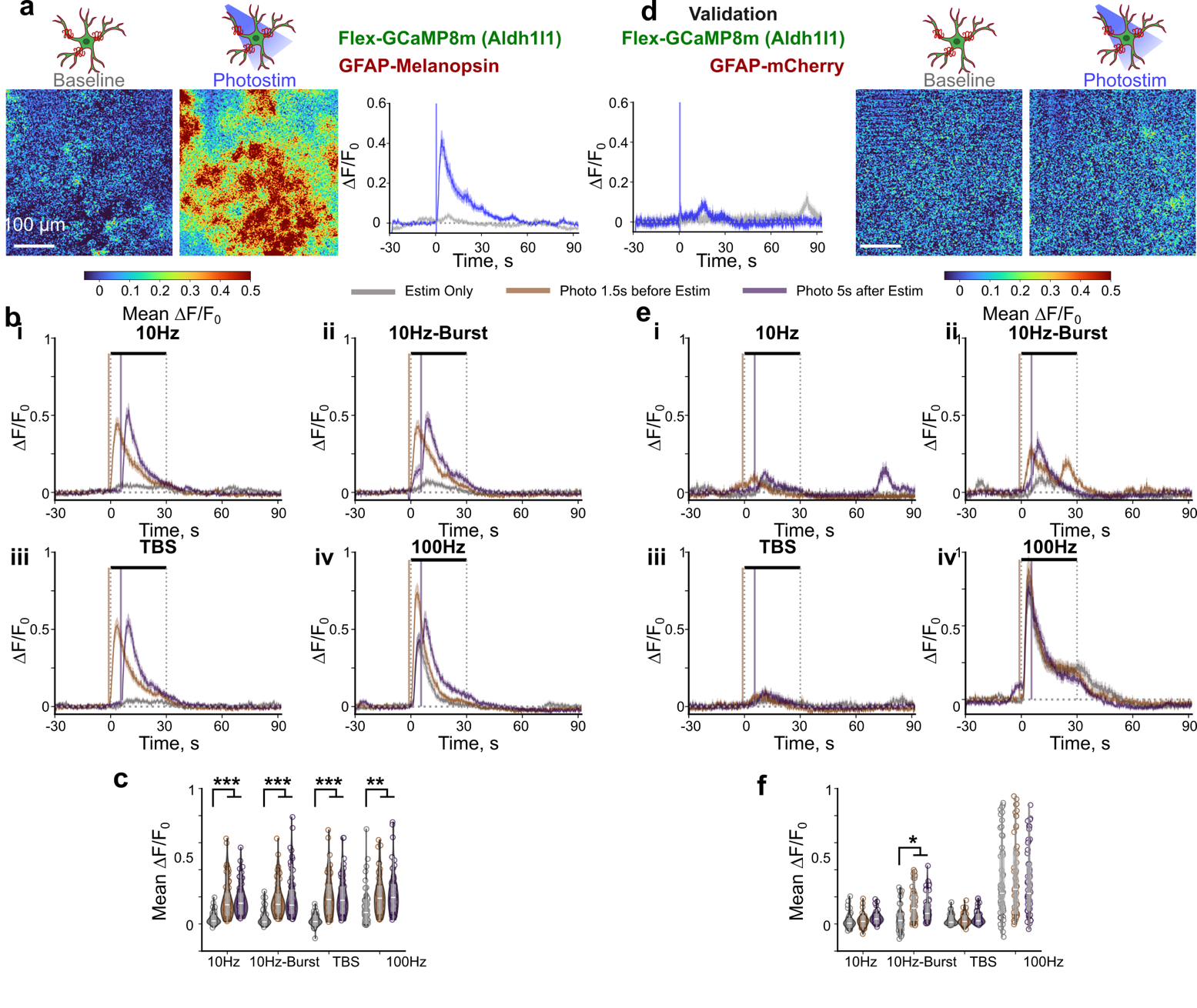

**Figure S11. Photostimulation of melanopsin expressing astrocytes significantly increases astrocyte calcium activity during ICMS. A**. Representative mean images demonstrating that photostimulation increases calcium activity within astrocytes (left) in mice expressing melanopsin and GCaMP8m in astrocytes. The mean calcium activity of all identified astrocytes was elevated following 1s of 1mW photostimulation at 10 Hz with a 30ms pulse width. **B**. Average traces of astrocyte calcium activity during different ICMS patterns showing that photostimulation elevates calcium activity whether photostimulation is delivered 1.5s before (orange traces) or 5s after ICMS (purple traces) compared to ICMS alone (gray traces). Notably, despite high levels of astrocyte calcium activity during 100 Hz stimulation alone, photostimulation still could amplify it suggesting it is not being saturated. **C**. the mean stimulation induced calcium activity is significantly elevated with both photostimulation conditions (LMM; Stim: p=1.7e-34, photostim: p=4.6e-5, interaction: p=2.4e-16) N=3, n=65. **D-F**. similar analyses in validation animals show that photostimulation in animals lacking the photosensitive opsin does not amplify astrocyte calcium activity (LMM; Stim: p=1.43e-215, photostim: p= 0.34, interaction: p=2.5e-6, N=2, n=58)*:p<0.05, **:p<0.01, ***:p<0.001.

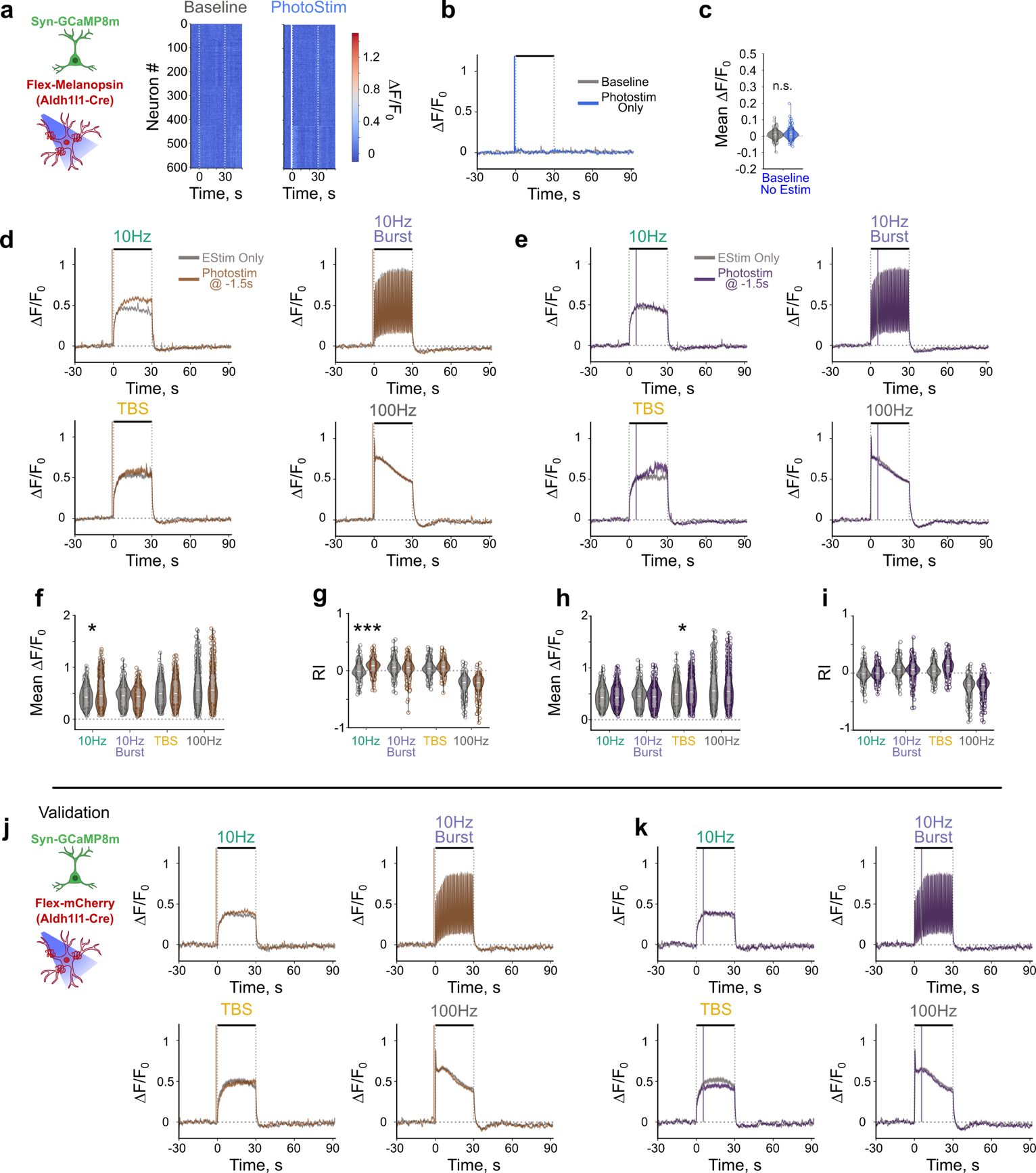

**Figure S12. Photostimulation of melanopsin expressing astrocytes does not globally affect neuronal responses to ICMS. A-b**. Heatmap of trial averaged neuronal responses (a) and traces of neuronal responses (b) shows that photostimulation of melanopsin in astrocytes does not increase neuronal calcium activity. C. Mean neuronal activity within 30 s of astrocyte photostimulation is comparable to baseline with no photostimulation. d-e. Traces of neuronal activity elicited by ICMS alone (gray) and astrocyte photostimulation pre-ICMS (d, orange) or post-ICMS (e, purple). F. Mean calcium activity of neurons during ICMS is increased with photostimulation of astrocytes pre-ICMS during 10 Hz stimulation (LMM Stim p<2e-16, Photostim: p = 0.76; Estim x Photo: p = 5.9e-24). G. Response index of neuronal activity during ICMS is increased with astrocyte photostimulation pre-ICMS during 10 Hz stimulation (LMM: Estim p <2e-16, Photostim: p = 0.46, interaction p = 1.3e-37). H. Mean neuronal activity of neuronal calcium activity during ICMS is increased with astrocyte photostimulation post-ICMS during TBS stimulation (LMM Estim: p <2e-16, Photostim: p = 0.92, interaction: p=3.6e-18). I Response index of neuronal activity is increased with photostimulation of astrocytes post-ICMS during TBS stimulation (LMM EStim: p<2e-16, Photostim: p = 0.31, interaction: p = 8.4e-38). j-k. Traces of neuronal calcium activity with photostimulation of animals lacking melanopsin pre-ICMS (j) or post-ICMS (k) does not show substantial change in neuronal response to ICMS. N=2-3 animals across 3 trials. *:p<0.05, **:p<0.01, ***:p<0.001.

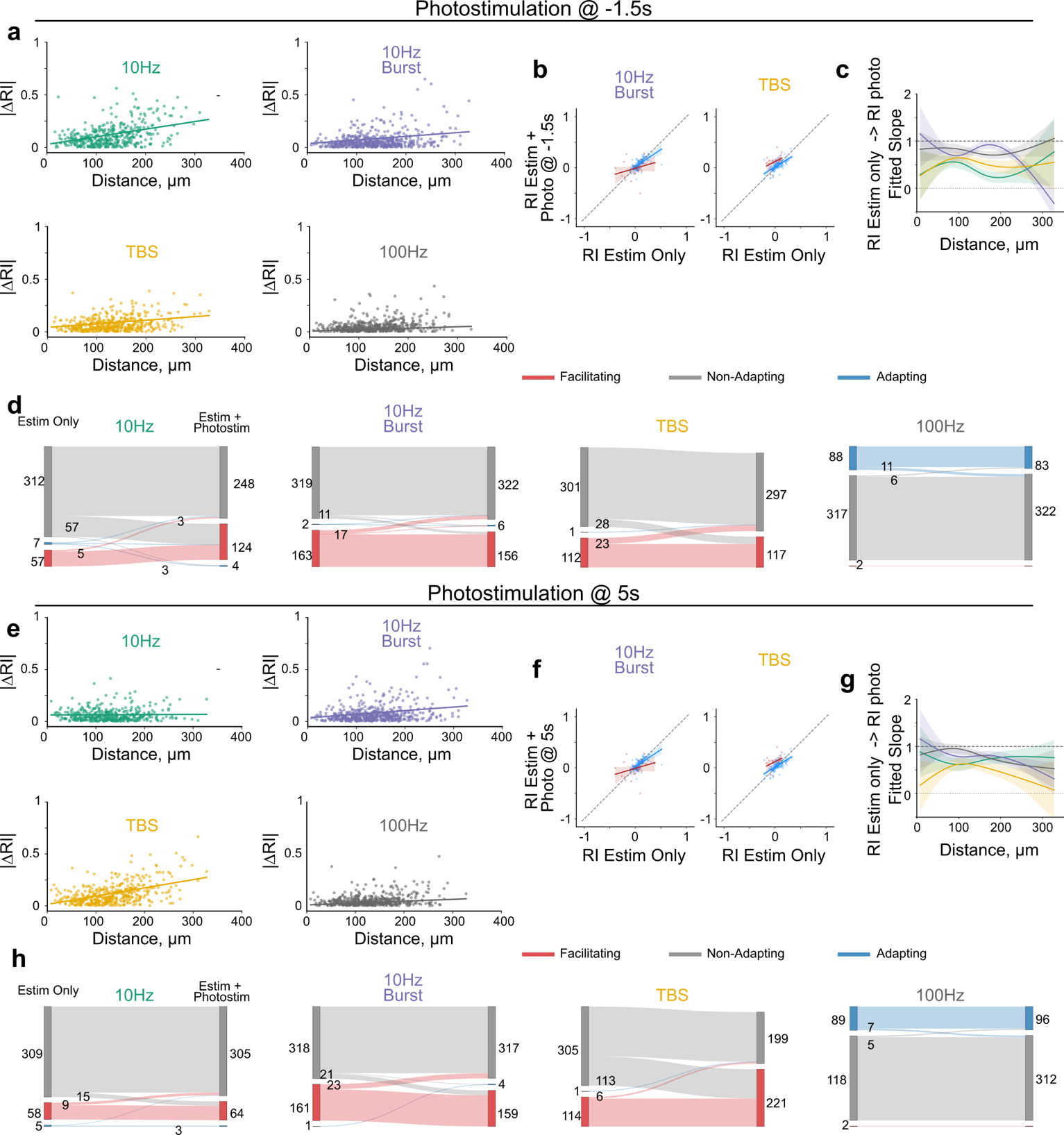

**Figure S13**. Photostimulation of astrocytes elicits heterogeneous effects on temporal profile of ICMS-induced neuronal activity. A. Magnitude of change of the response index with astrocyte photostimulation pre-ICMS increases with distance from the electrode depending on stimulation parameter. Scatter plots of fitted slopes in Fig. 5i (see Fig. 5i for stats). B-C. Fitted slope over distance of RI of neurons with photostimulation of astrocytes pre-ICMS over RI of neurons with ICMS only (see Fig. 5k-l for stats). D. Small populations of neurons change their response profile with photostimulation of astrocytes pre-ICMS. E. Magnitude of change of the response index with astrocyte photostimulation post-ICMS increases with distance from the electrode depending on stimulation parameter. Scatter plots of fitted slopes in Fig. 5j (See Fig. 5j for stats). f. Fitted slope over distance of RI of neurons with photostimulation of astrocytes post-ICMS over RI of neurons with ICMS only (See Fig. 5m-n for stats). f. Small populations of neurons change their response profile with photostimulation of astrocytes post-ICMS.

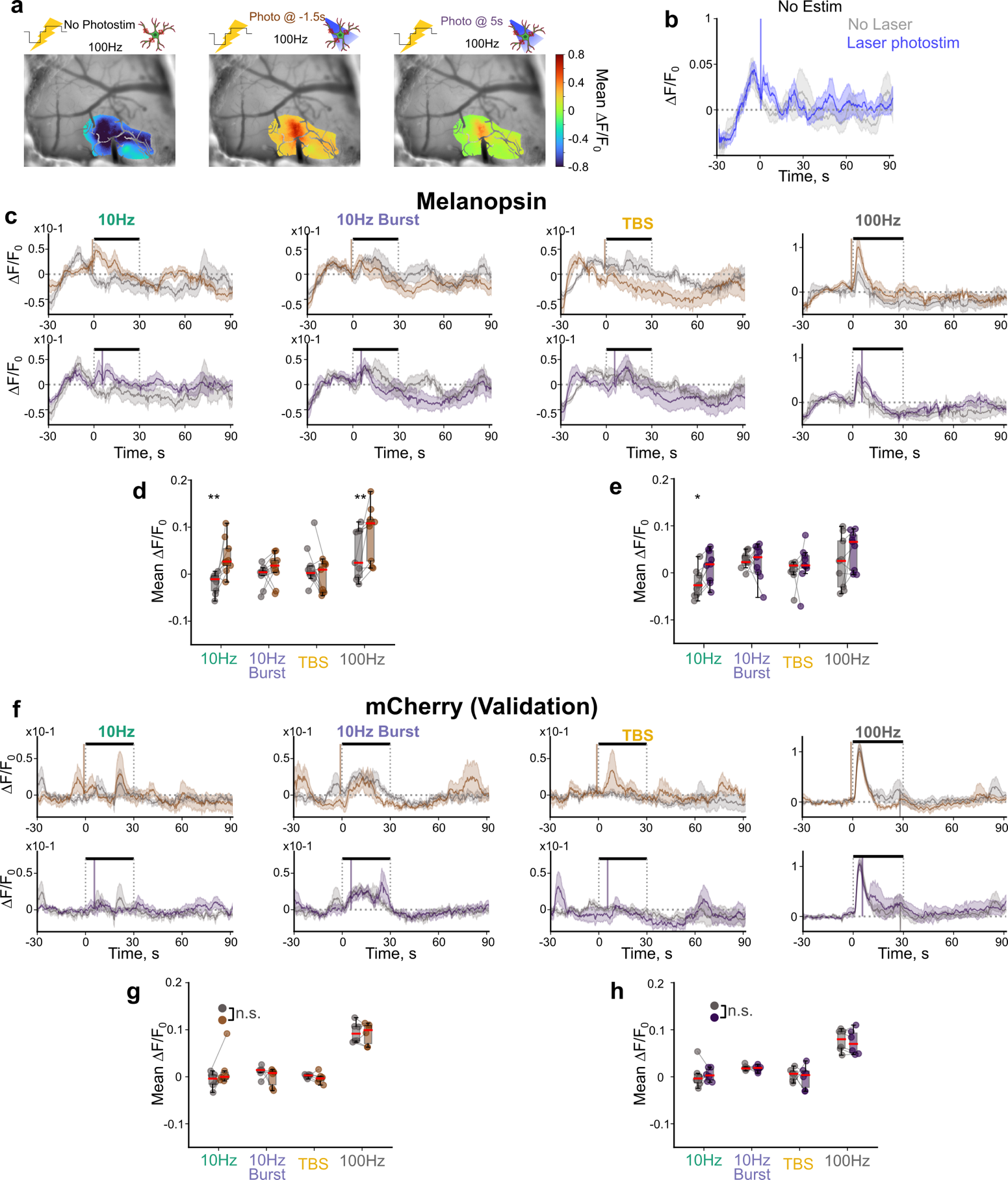

**Figure S14. Laser photostimulation of melanopsin transiently increases astrocyte calcium activity during ICMS. A**. Representative mean projection images during 100 Hz stimulation shows that astrocyte calcium activity is higher with laser photostimulation than without. **B**. The imaging modality is sufficient to increase astrocyte calcium activity during the baseline period before laser photostimulation. Laser photostimulation can still increase astrocyte calcium activity (blue trace) compared to without (gray) **C**. Laser photostimulation 1.5s before (orange) or 5s after (purple) transiently amplifies astrocyte calcium activity during stimulation more than electrical stimulation alone (gray). **D-E**. Astrocyte calcium activity is significantly higher 5s after photostimulation compared to no stimulation (gray) whether laser photostimulation is delivered 1.5s before electrical stimulation onset (d; LMM; Stim: p=9.9e-7, photostim: p= 4.9e-3, interaction: p=3.9e-3) or 5s after electrical stimulation onset (e; LMM; Stim: p=3.9e-3, photostim: p=0.019, interaction: p=0.25). **F**. Astrocyte calcium activity is comparable between trials without photostimulation (gray) and with laser photostimulation 1.5s before (orange) or 5s after (purple) electrical stimulation onset in validation animals. **G-H**. Photostimulation does not significantly affect astrocyte calcium activity in the 5s after photostimulation delivery in validation animals (g; LMM; Stim: p=1.74e-14, photostim: p=0.22, interaction: p=0.97; H; LMM; Stim: p=1.7e-11, photostim: p= 0.68, interaction: p=0.97). N=2-3 animals across 3 trials. *:p<0.05, **:p<0.01, ***:p<0.001.

**Supplemental Movie 1: 100 Hz ICMS elicits a propagating wave of calcium activity in astrocytes.** AAV-Flex-GCaMP8m and AAV-Syn-jRGECO1a were injected into the visual cortex of Aldh1l1-Cre mice. Four weeks later a single shank Michigan-style electrode was implanted into layer II/III of the infected region, and the window was sealed for awake in vivo imaging. The raw two-photon image series was downsampled to 3Hz and astrocyte activity was filtered with a 7-frame median filter, then converted to ∆F/F0, where F0 represents the average pixel intensity over the 30 s baseline period. The ∆F/F0 signal was clipped at 0.05 such that activity below this value is not shown. The ∆F/F0 signal is overlayed on top of a grayscale mean projection image to highlight the morphology of the astrocytes. Stimulation is delivered for 30 s following a 30s baseline period, and the video is played at 5x speed.

**Supplemental Movie 2: Two-photon recording of neurons and astrocytes during 100 Hz ICMS.** AAV-Flex-GCaMP8m and AAV-Syn-jRGECO1a were injected into the visual cortex of Aldh1l1-Cre mice. Four weeks later a single shank Michigan-style electrode was implanted into layer II/III of the infected region, and the window was sealed for awake in vivo imaging. Raw images were denoised with singular value decomposition to better highlight astrocyte (green) and neuronal (red) morphology and activity increases, and images were downsampled to 3 Hz by averaging frames together and the video is played at 5x speed. 100 Hz stimulation was delivered for 30s following a 30s baseline.
